# Co-infecting phages impede each other’s entry into the cell

**DOI:** 10.1101/2023.06.05.543643

**Authors:** Thu Vu Phuc Nguyen, Yuchen Wu, Tianyou Yao, Jimmy T. Trinh, Lanying Zeng, Yann R. Chemla, Ido Golding

## Abstract

Bacteriophage lambda tunes its propensity to lysogenize based on the number of viral genome copies inside the infected cell. Viral self-counting is believed to serve as a way of inferring the abundance of available hosts in the environment. This interpretation is premised on an accurate mapping between the extracellular phage-to-bacteria ratio and the intracellular multiplicity of infection (MOI). However, here we show this premise to be untrue. By simultaneously labeling phage capsids and genomes, we find that, while the number of phages landing on each cell reliably samples the population ratio, the number of phages entering the cell does not. Single-cell infections, followed in a microfluidic device and interpreted using a stochastic model, reveal that the probability and rate of individual phage entries decrease with MOI. This decrease reflects an MOI-dependent perturbation to host physiology caused by phage landing, evidenced by compromised membrane integrity and loss of membrane potential. The dependence of phage entry dynamics on the surrounding medium is found to result in a strong impact of environmental conditions on the infection outcome, while the protracted entry of co-infecting phages increases the cell-to-cell variability in infection outcome at a given MOI. Our findings demonstrate the previously unappreciated role played by entry dynamics in determining the outcome of bacteriophage infection.

---

Bacteriophage lambda serves as a paradigm for the decision made by temperate phages between rampant replication, resulting in release of viral progeny and cell death (lysis), and dormancy in a prophage state (lysogeny), which allows the host bacterium to live and reproduce (*1, 2*). The best-characterized factor affecting this choice is the number of lambda phages co-infecting the cell (multiplicity of infection, MOI) (*3, 4*). Infection by a single phage typically results in lysis, whereas a higher MOI results in lysogeny (*5, 6*). The effect of MOI is mediated by a phage-encoded circuit that chooses the transcriptional program to be executed based on the number of lambda genomes that entered the cell (*7, 8*). In terms of its utility, viral self-counting is believed to serve as a way of inferring the abundance of available hosts in the environment (*3, 9*). Specifically, simultaneous infection by multiple phages (i.e., MOI > 1) implies that phages outnumber bacteria, thus the release of new progeny via lysis will be futile and lysogeny should be chosen. This interpretation is consistent with other examples where the relative abundance of phages and bacteria impacts infection outcome (*10*). However, it is premised on an accurate mapping between the environmental phage-to-bacteria ratio and the intracellular MOI, a mapping that has not been directly tested. This accurate mapping is further called into doubt by the fact that, at the single-cell level, the relation between MOI and infection outcome is highly probabilistic rather than threshold-like (*6, 11*). Thus, the relationship between the population ratio and the number of internalized phage genomes merits careful examination.

To examine infection at the single-cell level, we utilized phages (λ *cI*857 *Pam*80 *stf*::P1*parS*-*kan*^*R*^, ref. (*8*)) whose capsid was decorated with gpD-mTurquoise2 or gpD-EYFP (*12*). Following infection (of host MG1655), the intracellular phage genome is detected via binding of mCherry-ParB or CFP-ParB to an engineered *parS* sequence (*13, 14*) (**Figure 1A** and ***Methods***). The infecting phage is unable to replicate on wild-type host, preventing intracellular genome number from increasing after entry (*8*). The labeling scheme enables reliable counting of both extracellular capsids and intracellular phage genomes (**Figures S1** and **S2**). We first asked how the numbers of adsorbed and intracellular phages changed as we infected cells with varying phage concentrations. We found that, over a ∼10-fold change in phage concentration, the average number of adsorbed phages per cell followed the phage-to-bacteria ratio (**Figure S3**), with the adsorption efficiency remaining constant at 0.71 ± 0.04 (**Figure 1B**, mean ± SE of the three samples, *N* = 201, 204, and 221 cells), consistent with reported values (*15*–*17*) (**Figure S4**). In contrast, the efficiency of phage entry decreased approximately twofold (from 0.36 ± 0.06 to 0.16 ± 0.01, **Figure 1B**). Consequently, the intracellular MOI did not reflect the bulk ratio of phages to bacteria.

**Figure 1.**
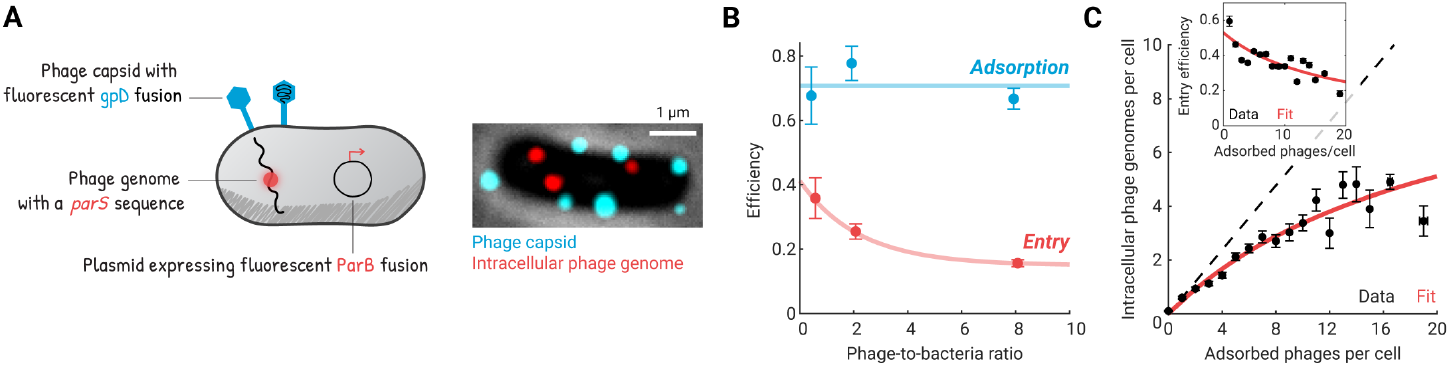
The intracellular viral copy number is not proportional to the extracellular phage-to-bacteria ratio. **(A)** Phage capsids and intracellular phage genomes were fluorescently labeled. Left, schematic of the labeling system. Right, an *E. coli* cell adsorbed by seven lambda phages (cyan spots), three of which have ejected their genomes (red spots). **(B)** The fraction of phages whose genomes entered the cell decreases at higher phage-to-bacteria ratios. Phages and bacteria were mixed at different ratios. The efficiencies of adsorption and entry were defined as the average numbers of adsorbed or intracellular phages per cell, respectively, divided by the phage-to-bacteria ratio. Markers, mean ± SE (*N* = 201, 204, and 221 cells for samples with mixing ratios of 0.5, 2, and 8, respectively). Cyan line, the average of the three samples. Red curve, fit to an exponential decay with a baseline, serving as a guide to the eye. **(C)** The number of intracellular phage genomes scales sublinearly with the number of phages adsorbed to the cell. Black markers, mean ± SE (*N* = 1437 cells, pooled from 7 independent experiments); cells at higher MOI were binned together to allow for at least 10 cells per bin. Red curve, fit to a Michaelis-Menten function, serving as a guide to the eye. Dashed line, linear scaling extrapolated from cells with one adsorbed phage. Inset, the efficiency of phage entry decreases with the number of adsorbed phages. The red curve was calculated using the same fit as in the main panel.

We next utilized the natural variability in the number of phages adsorbed to each cell (*6, 8*) (**Figure S5**) to probe the relation between the adsorbed and intracellular phage numbers at the single-cell level. We found that the efficiency of phage entry decreased from approximately 60% in cells adsorbed by a single phage (MOI = 1, consistent with (*17, 18*)) to approximately 30% at MOI = 10 (**Figure 1C**, *N* = 1437 cells, pooled from 7 independent experiments). In other words, as more phages adsorb to the cell, a smaller fraction of their genomes successfully enters the cell. As a result, the number of intracellular phage genomes scales sublinearly with MOI (**Figure 1C**).

To gain insight into the observed reduction in entry efficiency at higher MOI, we next aimed to examine the temporal kinetics of phage entries in individual cells. To that end, we performed infection in a microfluidic device, tracking phage adsorption and entry in real time (**Figures 2A, S6**, and ***Methods***). The experiments provided the time series of phage adsorption and entry events in each infected cell (**Figures 2A** and **S7**). To interpret these time series, we formulated a simple stochastic model where phage entry is governed by three parameters (**Figure 2B**): *η*, the entry probability (at infinite time); *k*, the rate (or probability per unit time) of initiating entry; and *τ*, the time between entry initiation and detection (see ***Methods***). For a cell adsorbed by *n* phages, the model is solved to yield the probability distribution for the number of intracellular phage copies at time *t*. We first tested the model on cells adsorbed by a single phage (*n* = 1). The model successfully captured the time-dependent average number of intracellular phages ⟨*λ*(*t, n* = 1)⟨ (**Figure 2C**) and the distribution of phage entry time *P*(*T*_1_|*n* = 1) (**Figure 2D**). The inferred parameter values (*η* = 0.56 ± 0.04, 1/*k* = 1.7 ± 0.2 min, *τ* = 35.9 ± 5.6 s, *N* = 208 cells pooled from 7 independent experiments, SE from bootstrapping) were consistent with the entry efficiency and mean entry time we observed in bulk and with reported values (*12, 17*–*20*) (**Figures S4** and **S8**), thus lending credence to the microfluidic-acquired data and the stochastic model used to interpret it.

**Figure 2.**
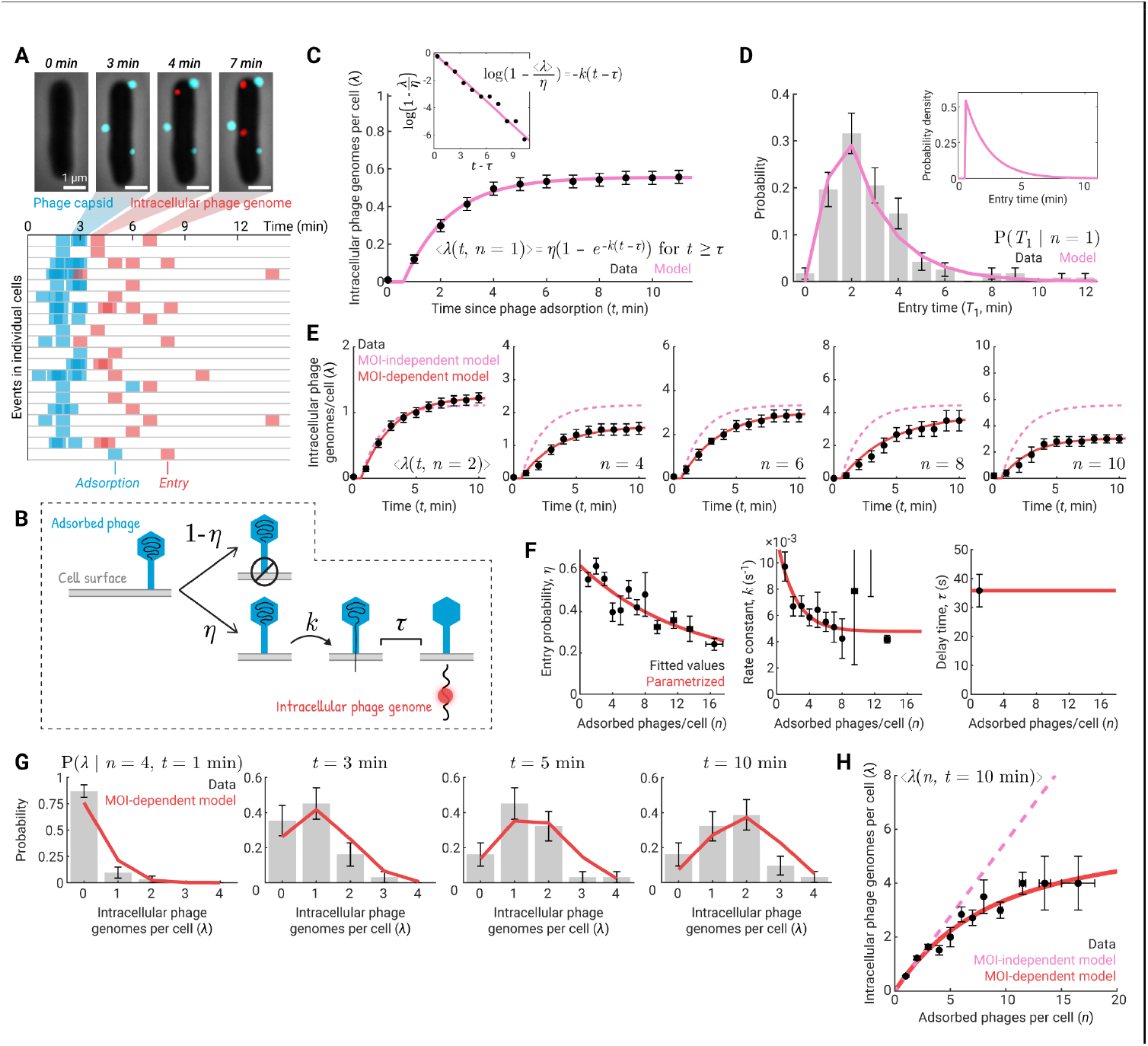
Time-lapse measurements indicate an MOI-dependent decrease in the probability and rate of phage entries. **(A)** Adsorption and entry of individual phages were followed in a microfluidic device. Top, an infected cell tracked over time. Bottom, for each cell, the times of adsorption and entry events were recorded. Data for 20 cells is shown. For all cells in this experiment, see **Figure S7.** **(B)** Schematic of the stochastic model for phage entry kinetics; see text for details. **(C)** The theoretical model captures the time-dependent average number of intracellular phages in cells adsorbed by one phage. Black markers, mean ± SE (*N* = 208 cells, pooled from 7 independent experiments). Pink curve, fit to the model. Inset, linearized data and model. **(D)** The model successfully predicts the distribution of entry times in cells adsorbed by one phage. Gray bars, histogram of the data, with error bars indicating SE. Pink curve, the probability distribution predicted by the model, binned by the imaging inter-frame interval. Inset, the theoretical probability distribution before binning. **(E)** A dependence of the entry parameters on MOI is required to capture the experimental data. Dashed curves in pink, predictions by a model in which *η* and *k* in cells with MOI > 1 are equal to those in cells with MOI = 1. Solid curves in red, predictions by a model in which *η* and *k* are allowed to vary with MOI. Black markers, data ± SE for MOI = 2, 4, 6, 8, and 10. For other MOIs, see **Figure S10.** **(F)** Inferred parameters for the MOI-dependent model. Black markers, fitted values ± SE from bootstrapping (*N* = 1030 cells, pooled from 7 independent experiments); cells at higher MOI were binned together to allow for at least 10 cells per bin. Red curves, parametrization: *η* and *k* as exponentially decaying functions of MOI, *τ* remaining the same as in cells with MOI = 1. **(G)** The MOI-dependent model successfully predicts the distributions of intracellular phage numbers at a given time. Gray bars, histograms of the data, with error bars indicating SE. Red curves, model predictions. Data for MOI = 4 at 1, 3, 5, and 10 minutes are shown. For other MOIs and other time points, see **Figure S12.** **(H)** The MOI-dependent model reproduces the sublinear relation between the numbers of adsorbed and intracellular phages. Black markers, mean ± SE; cells at higher MOI were binned together to allow for at least 10 cells per bin. Dashed line in pink, prediction by the MOI-independent model. Red curve, prediction by the MOI-dependent model. Data at 10 minutes following phage adsorption is shown. For other time points, see **Figure S13**.

Having calibrated our model based on singly infected cells, we next used it to predict the entry kinetics in cells adsorbed by two or more phages (**Figure 2E**). We found, however, that as the MOI increased, the prediction of intracellular phage number became poorer (**Figure S9**). In particular, the model overestimated the average number of intracellular phages at any given time (**Figures 2E** and **S10**). In light of our observations in **Figure 1C** above, we reasoned that the failure of the model reflected a decrease in phages’ ability to enter the cell at higher multiplicities. To test this hypothesis, we allowed model parameters to now vary with MOI. The revised model successfully reproduced the average kinetics (**Figure 2E**) and revealed a decrease in both entry probability *η* and rate *k* with MOI, and no MOI dependence for *τ* (**Figure 2F** and **S11**). Parametrizing *η* and *k* as exponentially decaying functions of MOI, the stochastic model also captured the distributions of intracellular phage numbers at a given time (**Figures 2G** and **S12**). Finally, the model successfully reproduced the sublinear relation between the numbers of intracellular and adsorbed phages, observed in both the microfluidic (**Figures 2H** and **S13**) and bulk experiments (**Figure S14**). Altogether, these results suggest an MOI-dependent impediment to cell entry by co-infecting phages.

How does coinfection impede phage entry into the cell? Despite decades of interrogation, the biophysical mechanisms driving the translocation of phage DNA from the capsid into the cytoplasm are still debated (*21*). Nevertheless, several aspects of phage entry, previously reported, are noteworthy. First, phage infection induces a strong ionic perturbation to the host cell, involving fluxes of solutes and a reduction or even loss of membrane potential (*22, 23*). Phage-induced perturbations were reported for diverse viral and bacterial species, including lambda and *E. coli* (*24, 25*). The severity of perturbation increases with MOI (*24, 26*) but does not require phage entry to occur, as evidenced by infection of “ghost” phage particles devoid of DNA (*26, 27*). Could these perturbations to the host cell underlie the diminished entry we find at higher MOI? Consistent with this idea, ionic conditions have been shown to impact the kinetics of DNA ejection in vitro (*21*), possibly by modulating the self-repulsion of the encapsidated DNA or altering the osmotic pressure that opposes ejection (*21*). In lambda, specifically, both the extent (*28*–*30*) and rate (*31, 32*) of DNA ejection are affected. Putting these elements together, we hypothesized that phage-induced perturbations to the host physiology, which occur following adsorption but prior to entry, impede phage entry into the cell in an MOI-dependent manner.

To directly probe the relation between phage-induced perturbation and DNA entry, we focused on two reported aspects of this perturbation: the compromise to membrane integrity and the reduction of membrane potential (*21*). To measure the membrane integrity, we sampled the infection mixture at different times and stained cells using propidium iodide (PI, ***Methods***). For each cell, we recorded the numbers of adsorbed and intracellular phages, as well as the intracellular PI fluorescence (**Figures 3A** and **S15**). Within 5 minutes of infection, cells became permeable to PI, with intracellular PI fluorescence rising linearly with the number of adsorbed phages (**Figures 3B** and **S16**). The MOI dependence of permeability is consistent with the idea that each adsorbed phage induces an opening in the cell membrane (*26, 33*). In agreement with reports that phage-induced perturbations do not require entry (*26, 27*), we found that PI permeation was not correlated with the intracellular phage numbers (**Figure S16**). We note that the permeation of PI into phage-infected cells did not reflect cell death or phage-induced lysis (***Methods*** and **Figure S15**). Examining PI fluorescence later in infection, we found that the degree of PI permeation per adsorbed phage decreased over time, indicating that the infected cells gradually recover membrane integrity (**Figure 3C**). Taken together, these results indicate that phage adsorption transiently permeabilizes the infected cell’s membrane in an MOI-dependent manner.

**Figure 3.**
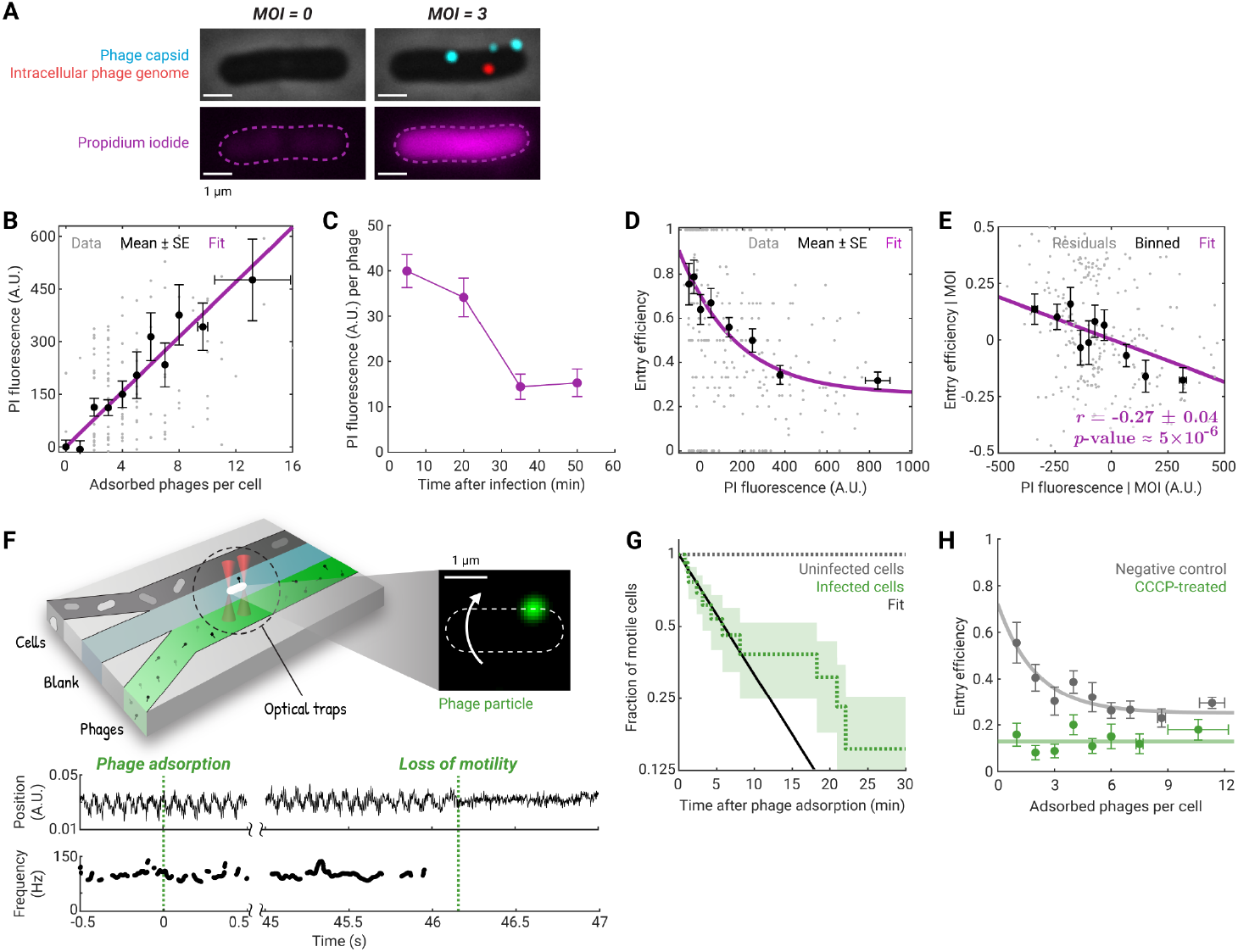
Phage adsorption causes cellular perturbation, resulting in impeded phage entry. **(A)** Following infection (phage capsids in cyan, intracellular genomes in red), cells were stained with propidium iodide (PI, shown in magenta). Dashed line, cell boundary. **(B)** Phages permeabilize the cell’s membrane in an MOI-dependent manner. The intracellular fluorescence of PI is plotted as a function of the number of adsorbed phages. Gray markers, single-cell values (*N* = 188 cells). Black markers, mean ± SE; cells at higher MOI were binned together to allow for at least 10 cells per bin. Magenta line, linear fit, the slope of which reflects the degree of membrane permeabilization per phage. Data at 5 minutes following infection is shown. For other time points, see **Figure S16.** **(C)** Infected cells recover membrane integrity. The slope of the linear fit between PI fluorescence and MOI at each time point is plotted. Error bars indicate SE from bootstrapping (*N* = 188, 132, 131, and 135 cells). **(D)** The efficiency of phage entry decreases in cells with stronger PI fluorescence. Gray markers, single-cell values (*N* = 298 cells pooled from *t* = 5 and 20 minutes). Black markers, mean ± SE (40 cells per bin). Magenta curve, fit to an exponential decay with a baseline. **(E)** Phage entry efficiency and PI fluorescence, conditioned on MOI, are negatively correlated. Gray markers, residuals obtained by linear regression of the PI fluorescence and of the entry efficiency on MOI (*N* = 298 cells). Black markers, mean ± SE of the residuals (30 cells per bin). Magenta line, linear fit. **(F)** The flagellar rotation frequency of phage-infected cells, indicating the membrane potential, is measured using optical traps. Top left, schematic of the trap setup and the flow chamber. Top right, a trapped cell with one adsorbed phage (stained using SYTOX Orange); the dashed line indicates the cell outline, and the white arrow indicates the cell body rotation. Bottom, the trapped cell position and the flagellar rotation frequency of the same cell. See also **Movie S1.** **(G)** Phage adsorption results in loss of motility. Dotted green line, the fraction of motile cells following phage adsorption (MOI = 1), with shading for SE (*N* = 13 cells). Black line, exponential fit to the data up to 10 minutes. Dotted gray line, the fraction of motile cells in the uninfected sample (*N* = 7 cells). **(H)** Complete membrane depolarization results in impeded phage entry. Green markers, mean (± SE) entry efficiency of cells depolarized using carbonyl cyanide *m*-chlorophenyl hydrazone (CCCP, *N* = 219 cells). Green line, the average over all cells in the CCCP-treated sample. Gray markers, mean ± SE for cells treated with 0.5% DMSO, serving as a negative control (*N* = 209 cells). Gray line, fit to an exponential decay with a baseline. For both samples, cells at higher MOI are binned together to allow for at least 5 cells per bin.

We next used the degree of PI permeation to test whether the compromise to membrane integrity impedes phage entry. Examining single-cell data up to 20 minutes post-infection (prior to the recovery of membrane integrity, **Figure 3C**), we found that cells with a stronger PI fluorescence exhibited lower efficiency of phage entry (ratio of intracellular phage genomes to adsorbed phages per cell, **Figure 3D**). In particular, the entry efficiencies found in the least and most permeabilized cells were comparable to the values observed above for low and high MOI, respectively (**Figure 1C**). These findings are consistent with the hypothesis that the impediment to entry reflects the severity of the infection-induced perturbation. To directly establish the causal link between the two phenomena, we calculated the correlation between entry efficiency and PI fluorescence, conditional on (i.e., for a given) MOI (*34*). Consistent with our hypothesis, the correlation coefficient, *r*(Entry efficiency, PI fluorescence | MOI), is negative (**Figures 3E** and **S17**). Although we cannot rule out the presence of additional mechanisms connecting phage adsorption and impeded entry (**Figure S18**), our causal inference analysis established that phage-induced compromise to membrane integrity causes entry impediment.

To examine phage-induced changes to *E. coli*’s membrane potential, we utilized as proxies the flagellar rotation frequency (*35*) and a fluorescent reporter, PROPS (*36*). We used dual-trap optical tweezers to trap cells in a flow chamber, into which chemicals and phages can be perfused (*37*). Trapped cells were fluorescently imaged, while their flagellar rotation frequency was inferred from the trap signal (*38*) (**Figure 3F** and ***Methods***). Exposing cells to carbonyl cyanide *m*-chlorophenyl hydrazone (CCCP), a protonophore that depletes the proton motive force, resulted in rapid loss of flagellar motility and an increase in fluorescence from PROPS (**Figures S19** and **S20**), as expected. A similar cessation of motility was exhibited following lambda adsorption (**Figures 3F** and **3G**). The measured half-life for cell motility, 5.6 ± 2.1 min (*N* = 13 cells, SE from bootstrapping), was consistent with the timescale of K^+^ efflux from lambda-infected cells (**Figure S21** and ref. (*25*)). The encapsidated DNA of the adsorbed phages could still be detected on cells that had lost motility (**Figure 3F** and **Movie S1**), indicating that membrane depolarization preceded phage entry. Phage-induced depolarization was also revealed by the changes to PROPS fluorescence, which mirrored the effect of CCCP treatment (**Figure S22**). Next, to test the prediction that membrane depolarization will in turn impede entry, we measured entry efficiency in CCCP-treated cells. While intracellular lambda genomes were still detected, the entry efficiency (∼15%, *N* = 219 cells) was lower than that of untreated cells (**Figure 3H**) and comparable to the value at high MOI (**Figures 3H, 2F**, and **1C** above). Depolarization-induced entry efficiency was also similar to that of highly permeabilized cells, as indicated by PI fluorescence (**Figure 3D** above). Thus, complete membrane depolarization emulates the perturbation caused by adsorption of multiple phages and results in significantly impeded phage entry.

Having illuminated some of the mechanistic underpinnings of impeded phage entry, we returned to the question motivating our investigation and tested whether entry dynamics impact lambda’s choice of lysogeny. Multiple reports indicate that the outcome of infection varies greatly with the environmental conditions (*39, 40*). Based on our findings above, we hypothesized that these conditions exert their influence, at least in part, by affecting the number of phages entering the cell, a number which in turn drives the choice of developmental program by the phage (*8*). The idea that media conditions affect entry was premised on the fact that the severity and duration of phage-induced perturbations strongly depend on these conditions (*41, 42*). And, indeed, performing infection in 12 different solutions used in microbiological studies revealed diverse relations between the numbers of adsorbed phages and intracellular phage genomes (**Figure S23**). Under some conditions, adsorption by as many as ten or more phages resulted, on average, in a single phage entry. We reasoned that this apparent limit on the number of intracellular phages has the potential to reduce the occurrence of lysogeny.

To test whether the diminished phage entry indeed affects cell fate, we assayed the frequency of lysogeny when infection takes place in two commonly used media with very different entry characteristics, LB (which enables multiple phage entries) and SM (which limits entry to a single phage, on average) (**Figures 4A** and **S23**). In both cases, cell growth pre- and post-infection took place in LB (*8*), and cells were not starved prior to infection (***Methods*** and **Figure S24**). Following ref. (*8*), we again used a replication-deficient phage, where the number of viral genomes remains constant following entry. Under each entry condition, we measured the fraction of cells undergoing lysogeny as a function of the phage-to-bacteria ratio. The two conditions resulted in distinct lysogeny-versus-MOI curves (**Figure 4B**). In particular, at higher MOI, when cells are likely absorbed by multiple phages, the frequency of lysogeny in SM saturated at a value ∼30-fold lower than that in LB. Thus, the choice of entry conditions dramatically impacted the propensity to lysogenize at high MOI.

**Figure 4.**
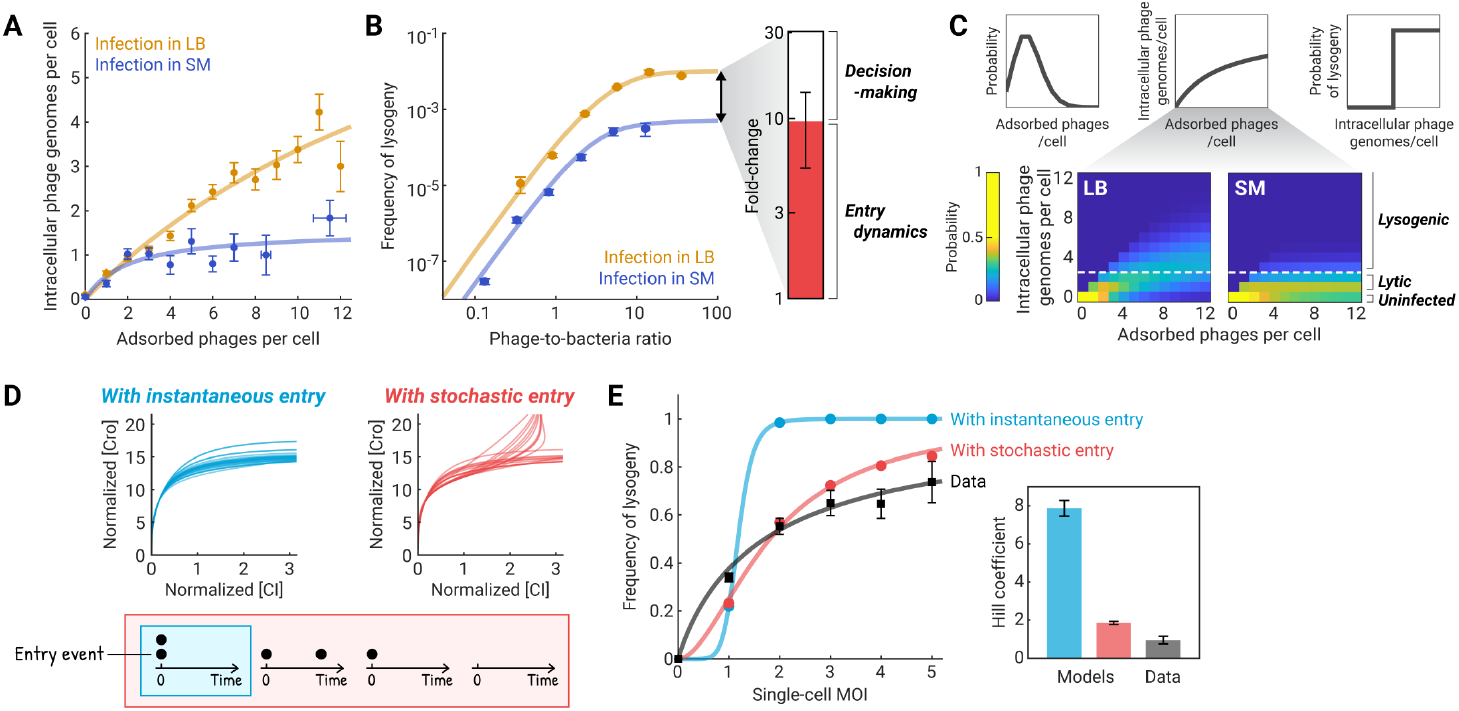
Entry dynamics impact the infection outcome and increase the variability in cell fate. **(A)** Infection media alter the degree to which lambda entry is impeded. Identically cultured cells were infected in different media, and the numbers of adsorbed and intracellular phages per cell were measured. Orange markers, mean ± SE for infection in LB (data reproduced from **Figure 1C**). Blue markers, infection in SM (*N* = 242 cells, at least 5 cells per bin). Orange and blue curves, fits to a Michaelis-Menten function, serving as a guide to the eye. For other infection media and buffers, see **Figure S23.** **(B)** Entry dynamics in different media accounts for most of the difference in lysogenization frequency. Left, the frequencies of lysogeny as a function of phage-to-bacteria ratios (corrected for adsorption efficiency) when infection is performed in LB (orange) and SM (blue). Markers, mean ± SE from technical duplicates. Orange and blue curves, fits to the model in **Panel C**. Right, entry dynamics (red) accounts for ∼10-fold of the observed ∼30-fold difference in the maximum frequency of lysogeny between the two media. The other ∼3-fold is due to differences in the single-cell probability of lysogeny (white, SE from bootstrapping). **(C)**Schematic of the model used to capture the data in **Panel B**. Top, the model accounts for phage adsorption, entry, and decision-making. Bottom, the predicted distributions of intracellular phage numbers in LB and SM, parametrized using the data in **Panel A**. Dashed lines, the minimum MOI required for lysogenization. **(D)** Stochastic phage entry results in different expression profiles of Cro (driving lysis) and CI (driving lysogeny). Top, predicted single-cell trajectories of Cro and CI concentrations in the Yao et al. (2021) model, assuming instantaneous (cyan) or stochastic (red) phage entry. Trajectories of 24 cells, each adsorbed by 2 phages, are shown for each case. For other MOIs, see **Figure S28**. Bottom, schematics of the temporal change in MOI for each model. **(E)** Entry dynamics increase the cell-to-cell variability in infection outcome at a given MOI. Black markers, data from Zeng et al. (2010). Cyan and red markers, predictions by the model of Yao et al. in the absence or presence of stochastic entry kinetics, respectively. Solid curves, fits to Hill functions. The Hill coefficients, which describe the degree of precision in the lysis vs. lysogeny decision, are shown on the right (SE from bootstrapping).

We interpreted the lysogenization curves in the two media using a simple model (**Figure 4C**) consisting of (i) random phage-bacteria collisions, resulting in a Poisson distribution of single-cell MOI (*8*); (ii) stochastic entry of phages into the cell, parametrized using the experimentally measured values (**Figure 4A**); and (iii) choice of lysogeny if three or more phage genomes entered the cell (*5, 8*) (***Methods***). The model was able to capture the experimental curves (**Figure 4B**). The inferred model parameters revealed that the difference in phage entry— specifically, the reduced probability in SM of having three or more intracellular phages, even at high MOI— accounted for most of the difference in lysogenization between the two media (**Figures 4B** and **S25**). In other words, when infecting in SM, the intracellular MOI poorly reflected the phage-to-bacteria ratio in the environment, resulting in a strong impact on infection outcome.

Finally, beyond its impact on the lysogeny phenotype as measured in bulk (**Figure 4B**), we asked whether entry dynamics can account for the reported cell-to-cell variability in infection outcome at a given MOI (*6, 43, 44*). While this heterogeneity has traditionally been attributed to the stochastic (“noisy”) expression of genes in the decision network (*45*), the idea lacks experimental evidence. We reasoned that, instead, the stochastic kinetics of phage entry may be key. This was motivated by recent works suggesting that, rather than simply measuring MOI, the lysis/lysogeny decision circuit weighs intracellular genomes by their arrival times, with latecomers contributing less to the outcome (*8, 16*). To test the consequences of stochastic entry, we first used our entry model (**Figure 2B** above) to create simulated time series of phage entries in many individual cells, under varying numbers of adsorbed phages (**Figures S26** and **S27**; see ***Methods***). We then used this series as the input to a deterministic model of the decision network (*8*), yielding the predicted kinetics for the concentration of the Cro and CI transcription factors, and thus the infection outcome, in each cell (**Figures 4D, S28**, and **S29**). To facilitate comparison with the single-cell data of ref. (*6*), we simulated infection by a replication-competent (wild-type) phage. When stochastic entry dynamics are ignored, i.e., all adsorbed phages are assumed to enter the cell instantaneously, cells with MOI = 2 show dominant CI expression, resulting in the choice of lysogeny (*8*). When entry dynamics is incorporated into the model, some cells adsorbed by two phages may have only one intracellular phage genome, or the second phage genome may arrive in the cell too late to contribute to the decision (*8, 16*). As a result, approx. 50% of the infected cells are Cro-dominant and will hence undergo lysis.

To quantify the heterogeneity in infection outcome, we calculated the resulting “decision curve”, which describes the fraction of cells undergoing lysogeny as a function of single-cell (extracellular) MOI. When entry dynamics is incorporated into the model, the predicted decision curve increased gradually rather than step-like, reminiscent of the data in ref. (*6*) (**Figure 4E**). The degree of precision of the decision can be quantified using the fitted Hill coefficient of this curve (*6, 46*). We found that incorporating entry dynamics into the model reduced the Hill coefficient from 7.9 ± 0.4 to 1.8 ± 0.1, thus considerably closer to the experimental value of 1.0 ± 0.1 in ref. (*6*). The “flattening” of the step-like response present in the original model (*8*) reflects the lower entry efficiency and slower entry in cells of higher MOI. That this effect is captured by a fully deterministic model of the decision suggests a diminished role for stochastic gene expression in explaining the observed heterogeneity of outcome among cells.

Considering that phage-induced perturbations and ion-modulated ejection, analogous to those found in lambda, have been reported for multiple phage-bacteria systems (*21*), it is likely that the MOI-dependent entry we report here extends beyond lambda. For temperate phages, the strong dependence on medium conditions provides a way for the environment to impact the choice of cell fate, by modulating the timing and number of phage genomes entering the cell. This previously unappreciated effect may help resolve the standing conflict between the simple MOI-to-lysogeny mapping observed in the lab and the complex, sometimes contradictory, relations found in natural phage habitats (*47*–*49*). Beyond the case of infection by multiple copies of the same phage, mutual impedance of entry may facilitate competition between phages of different species that coinfect the same host (*50*), where the effect may be seen as a form of superinfection exclusion (*51*).

The stochastic and protracted entry of co-infecting phages, demonstrated here, may also have implications for the continuous arms race between phages and bacteria (*52*). From the point of view of the infected cell, the sensitivity of phage entry to physiological parameters may provide an opportunity for the host to delay a critical step in the infection cycle, while its defenses are triggered (*25, 53, 54*), and offer regulatory opportunities to halt this entry altogether. Reversely, delayed entries may facilitate cooperation between co-infecting phages, with early-arriving phages counteracting the host’s defense system (*53, 55*), allowing later-arriving phages to survive and successfully propagate (*56*).

## ACKNOWLEDGEMENTS

We are grateful to the following people for their generous advice: M. Gruebele, M. Goulian, C. Herman, K. Maxwell, I. Molineux, T. Pilizota, A. Sokac, K. Venken, T. Wensel, Z. Yu, C. Zong, and all members of the Golding lab. Work in the Golding lab is supported by the National Institutes of Health grant R35 GM140709 and by the Alfred P. Sloan Foundation. Work in the Chemla lab is supported by the National Science Foundation Physics Frontiers Center (PFC) “Center for the Physics of Living Cells” (CPLC), grant PHY 1430124, and by the National Institutes of Health grant R35 GM144125. Work in the Zeng lab is supported by the National Science Foundation grant MCB 2013762. We gratefully acknowledge the computing resources provided by the Computational and Integrative Biomedical Research Center of Baylor College of Medicine.

## SUPPLEMENTARY INFORMATION

Methods

Figures S1 to S34 Movie S1

Tables S1 to S5

References for Supplementary Information

