## Supplementary Information for "Co-infecting phages impede each other’s entry into the cell"

***This file includes:***

**SUPPLEMENTARY METHODS ........ S3**
1. Bacterial strains, phages, and plasmids ........ S3
2. Chemical reagents, growth media, and buffers ........ S3
3. Phage preparation ........ S3
4. Bacterial growth conditions ........ S4
5. Measuring the numbers of adsorbed phages and intracellular phage genomes following bulk infection ........ S5
6. Measuring phage-induced membrane permeabilization using propidium iodide ........ S6
7. Measuring the time of the first phage entry following bulk infection ........ S8
8. Measuring the kinetics of phage entries in the microfluidic device ........ S9
9. Validating the ParB-*parS* labeling system using SYTOX Orange ........ S11
10. Validating the CCCP treatment protocol using PROPS ........ S12
11. Microscopy ........ S12
12. Image analysis ........ S13
13. Optical trap assay ........ S13
14. Preparation of representative images ........ S15
15. Potassium efflux assay ........ S15
16. Measuring the frequency of lysogeny following infection in different media ........ S15
17. Modeling bulk lysogenization following phage entry in different media ........ S17
18. Stochastic model of phage entry kinetics ........ S18
19. Simulating the stochastic model ........ S24
20. Stochastic simulation of infection outcome in individual cells ........ S25

**SUPPLEMENTARY FIGURES ........ S28**
Figure S1. Efficiency of phage labeling using fluorescent capsids and SYTOX Orange. ........ S28
Figure S2. Efficiency of genome detection using the ParB-*parS* system. ........ S29
Figure S3. Relation between the numbers of adsorbed and intracellular phages per cell and the phage-to-bacteria ratio. ........ S30
Figure S4. Comparison with literature-reported values for the efficiency and rate of phage adsorption and entry. ........ S31
Figure S5. Distribution of the number of phages adsorbed to the cell. ........ S32
Figure S6. The number of adsorbed phages as a function of perfusion time and phage concentration. ........ S33
Figure S7. Kinetics of phage adsorption and entry in the microfluidic infection assay. ........ S34
Figure S8. Kinetics of the first phage entry following bulk infection. ........ S35
Figure S9. The MOI-dependent $\eta, k$ model captures the time-lapse infection data most closely. ........ S36
Figure S10. The MOI-dependent model successfully fits to the time-dependent average number of intracellular phage genomes. ........ S37
Figure S11. Fitted parameter values in the MOI-dependent $\eta, k, \tau$ model. ........ S38
Figure S12. The MOI-dependent model successfully predicts the distributions of intracellular phage numbers ........ S39

Figure S13. The MOI-dependent model reproduces the sublinear relation between the numbers of intracellular and adsorbed phages following infection in the microfluidic device. ....S40
Figure S14. The MOI-dependent model captures the sublinear relation between the numbers of intracellular and adsorbed phages following bulk infection. ....S41
Figure S15. Controls for experiments involving propidium iodide (PI). ....S42
Figure S16. Correlation between intracellular PI fluorescence and MOI. ....S43
Figure S17. Inferring a causal link between membrane permeabilization and impaired phage entry. ....S44
Figure S18. Inferring additional causal links between phage adsorption and impaired phage entry. ....S45
Figure S19. Validating the CCCP treatment protocol using PROPS. ....S46
Figure S20. Response of optically-trapped cells to CCCP treatment. ....S47
Figure S21. Potassium efflux following phage infection. ....S48
Figure S22. Membrane depolarization of optically-trapped cells following phage adsorption, measured using PROPS. ....S49
Figure S23. Relation between the numbers of intracellular phage genomes and adsorbed phages in various infection media. ....S50
Figure S24. Comparing the lysogenization assay in this study vs. the literature. ....S51
Figure S25. Frequency of lysogeny as a function of phage-to-bacteria ratio, predicted using different models of entry dynamics. ....S52
Figure S26. Simulated time series of phage entry events. ....S53
Figure S27. Distributions of intracellular phage numbers, simulated using different models of entry kinetics. ....S54
Figure S28. Trajectories of Cro and CI concentrations, simulated using different models of entry kinetics. ....S55
Figure S29. Distributions of infection outcomes, simulated using different models of entry kinetics. ....S56
Figure S30. Comparison of the stochastic simulation and analytical predictions for the time-dependent average number of intracellular phage genomes. ....S57
Figure S31. Comparison of the stochastic simulation, analytical predictions, and phenomenological fit for the relation between the numbers of intracellular phage genomes and adsorbed phages ....S58
Figure S32. Comparison of the stochastic simulation, analytical predictions, and phenomenological fit for the efficiency of phage entry. ....S59
Figure S33. Comparison of the stochastic simulation, analytical predictions, and phenomenological fit for the distribution of the intracellular phage number. ....S60
Figure S34. Comparison of the stochastic simulation and analytical predictions for the distribution of phage entry time. ....S61

**SUPPLEMENTARY MOVIE ....S62**
Caption for Movie S1: Phage adsorption results in membrane depolarization. ....S62

**SUPPLEMENTARY TABLES ....S63**
Table S1: Bacterial strains, phages, and plasmids used in this study ....S63
Table S2: Chemical reagents used in this study ....S64
Table S3: Growth media and buffers used in this study ....S65
Table S4: Filter sets used for fluorescence microscopy ....S66
Table S5: Variables and parameters in the mathematical models ....S67

**REFERENCES FOR SUPPLEMENTARY INFORMATION ....S68**

***Other Supplementary Materials for this manuscript include the following:***

Movie S1

### SUPPLEMENTARY METHODS

#### 1. Bacterial strains, phages, and plasmids

All strains and plasmids used in this study are listed in **Table S1**. The phage strain used in this study, $\lambda_{TY11}$ (λ *cI*857 *Pam*80 *stf*::P1*parS-kan*$^R$) (*1*), is unable to replicate in wild-type (MG1655) host. Capsids of the infecting phages were labeled using gpD-mTurquoise2 or gpD-EYFP fusions (*2*). Intracellular phage genomes, each harboring a *parS* sequence, were labeled using mCherry-ParB or CFP-ParB fusions (*1*, *3*). Plasmid transformation (using a Bio-Rad MicroPulser Electroporator), colony plating, and phage titering followed standard protocols (*4*, *5*).

#### 2. Chemical reagents, growth media, and buffers

Reagents used in this study, excluding those involved in the phage purification protocol (**Section 3**) and the optical trap assay (**Section 13**), are listed in **Table S2**. Stock solutions were prepared using Milli-Q water (MilliporeSigma) or DEPC-treated water (Invitrogen).

Growth media and buffers used in this study are described in **Table S3**. The growth medium in most experiments was LB (Lennox formulation) (*4*). The medium in optical trap experiments was tryptone broth (TB). Plaque assays to titer phages were performed using NZYM (*1*, *6*). Supplements to growth media are specified below for each experiment. Agar plates and soft agar were prepared by supplementing the medium with 1.5% and 0.7% (w/v) agar (BD Biosciences), respectively. All media were autoclaved at 121°C for at least 25 minutes in a liquid cycle for sterilization.

#### 3. Phage preparation

##### *3.1. Phage purification*

This procedure was performed as described in (*6*). Briefly, fluorescently labeled phages were produced from LE392 $\lambda_{TY11}$ lysogens harboring plasmids that express gpD-mTurquoise2 or gpD-EYFP (**Table S1**). Crude phage lysates were obtained using heat induction, followed by precipitation using polyethylene glycol (PEG), CsCl ultracentrifugation, and dialysis into SM buffer. For the ultracentrifugation step, the step and equilibrium gradients were both set up using 14 mL ultracentrifuge tubes, and spun at 4°C using a Beckman SW40Ti rotor for 4–6 hours at 24,000 rpm, or 20–24 hours at 35,000 rpm, respectively.

The concentration of the purified phage stock (typically $10^{11}$–$10^{12}$ plaque-forming units [PFU]/mL) was measured using LE392 indicator cells. To measure the efficiency of capsid labeling, the phages were stained using 5 µg/mL DAPI (see **Table S2**) (*6*). Then, 1 µL of stained phages was mounted between a 24×50 mm coverslip no. 1 and a 22×22 mm coverslip no. 1 (Fisher Scientific), and imaged as described in **Section 11**. The efficiency of capsid labeling is defined to be the fraction of detected phages with both capsid and DAPI signals (97.0 ± 1.0%, mean and standard error, SE, from 3 independent phage purification runs, **Figure S1A**).

##### *3.2. SYTOX Orange staining*

In some experiments, phage particles were stained with SYTOX Orange (see **Table S2**), a DNA-intercalating fluorescent dye. Here, we followed the protocols from (*7*, *8*). The purified phage stock was first diluted as necessary using the same medium or buffer as the subsequent infection step. Phages were mixed with 500 nM SYTOX Orange and incubated in the dark at room temperature (RT) for 3 hours. Excess SYTOX Orange was removed using Slide-A-Lyzer MINI Dialysis Devices (2K MWCO, ThermoScientific) in two rounds of dialysis, each round for 2 hours at RT in at least 500× the phage volume.

The efficiency of SYTOX Orange labeling, obtained by imaging as described above, is defined to be the fraction of detected capsids with colocalized SYTOX Orange signal (91.9% ± 1.6%, mean and SE from 3 independent staining replicates, **Figure S1B**). We note that, following dialysis, the fraction of stained phages declined by approx. 20% within one day, and by approx. 50% within one week (**Figure S1C**). Therefore, this staining and dialysis protocol was always performed on the same day as the infection experiment.

### 4. Bacterial growth conditions

##### *4.1. Equipment*

Bacterial agar plates were incubated in Isotemp Microbiological Incubators (Fisherbrand). Liquid cultures were grown with aeration (using 220 rpm shaking) in a MaxQ 4000 Benchtop Orbital Shaker (Thermo Scientific), a MaxQ 7000 Water Bath Orbital Shaker (Thermo Scientific), or an Excella E24 Incubator Shaker (New Brunswick Scientific). Optical density (OD) was measured using a SmartSpec Plus spectrophotometer (Bio-Rad).

##### *4.2. Plating and overnight cultures*

Cells were streaked from 15% glycerol stocks (stored at -80°C) onto LB agar plates (supplemented with 100 µg/mL ampicillin and/or 50 µg/mL kanamycin when applicable) and incubated at 30°C for approx. 16 hours. For overnight cultures, fresh colonies were inoculated into 2 mL of growth medium (supplemented with antibiotics at the same concentration as the agar plates when applicable) in 14 mL round-bottom test tubes (Falcon). Overnight cultures were grown for approx. 16 hours at 30°C with aeration.

##### *4.3. Experimental cultures*

To prepare cells expressing mCherry-ParB or CFP-ParB (MG1655 harboring p2973 or pALA3047), we followed the protocol of (*1*). Experimental cultures (“overday” cultures) were prepared by diluting the overnight cultures by at least 1:500 into LBMM supplemented with IPTG in baffled Erlenmeyer flasks; the culture volume was 10–20% of the flask volume. The medium was supplemented with 100 µM or 10 µM IPTG to induce the expression of mCherry-ParB or CFP-ParB from p2973 or pALA3047, respectively. We note that inducing p2973 using 10 µM IPTG resulted in poor fluorescence, whereas inducing pALA3047 using 100 µM IPTG may result in aggregates (*9*). The cultures were grown at 37°C with aeration to $OD_{600}$ ≈ 0.3–0.4 and harvested as described in **Sections 5–9**.

To prepare cells expressing the proteorhodopsin optical proton sensor (PROPS) for all experiments except for those involving the optical traps (**Section 13**), we followed the protocol of (*10*). Cultures of MG1655 harboring pJMK001 were grown in LB at 33°C with aeration. When the culture reached $OD_{600}$ ≈ 0.3, 0.2% *L*-arabinose (for $P_{araBAD}$ induction) and 5 µM all-*trans* retinal (the chromophore of PROPS) were added to the cultures. Growth was resumed in the dark for 3 hours, after which the cultures were harvested as described in **Section 10**. We note that MG1655 harboring pJMK001, when cultured with 10 mM $MgSO_4$, showed no detectable PROPS fluorescence. Hence, $MgSO_4$ was omitted during growth and only supplemented before the infection step to enable phage adsorption (*11*). Because PROPS is expressed from an arabinose-induced promoter, we also omitted maltose to avoid interference with arabinose uptake. Previous studies have shown that adsorption of lambda phages still occurred in the absence of maltose in the growth medium (*11*, *12*).

To prepare motile cells for optical trap experiments (**Section 13**), we followed the protocol of (*13*). Cultures of MG1655 or MG1655 harboring pJMK001 (expressing PROPS) were grown in TB at 30°C with aeration. For MG1655, the cultures were harvested upon reaching $OD_{600}$ ≈ 0.5 as described in **Section 13**. For MG1655 harboring pJMK001, induction of PROPS was performed as described above.

To prepare cells for the potassium efflux assay or the lysogenization assay (**Sections 15** and **16**), cultures of MG1655 were grown in LBMM or LBM, respectively, at 37°C with aeration, and harvested upon reaching $OD_{600}$ ≈ 0.3–0.4 as described in **Sections 15** and **16**.

#### 5. Measuring the numbers of adsorbed phages and intracellular phage genomes following bulk infection

##### *5.1. Preparation of cells and phages*

Cultures of MG1655 harboring p2973 or pALA3047 were grown at 37°C in LBMM supplemented with IPTG as described in **Section 4**. When the cultures reached $OD_{600} \approx 0.3$–0.4, 10 mL of cells were transferred to a centrifuge tube (Corning) and spun at 2000×g for 10 minutes at 4°C using a Thermo Scientific Sorvall Legend XTR centrifuge, and the supernatant was removed. The cell pellet was resuspended in 1 mL of an ice-cold medium or buffer (dictated by the subsequent infection step) and transferred to a 1.5 mL microcentrifuge tube (Eppendorf). The cell suspension was centrifuged at 21130×g for 30 seconds at RT using an Eppendorf 5420 centrifuge, and the supernatant was removed. The cell pellet was resuspended in 100 μL of the ice-cold medium or buffer and used for infection immediately.

The purified phage stock ($\lambda_{TY11}$ with fluorescently labeled capsids, described in **Section 3**) was diluted to $2\times10^{10}$ PFU/mL using the same medium or buffer as the cell resuspension. In most experiments, gpD-mTurquoise2 phages were used to infect MG1655 harboring p2973 (expressing mCherry-ParB). In some experiments, including those involving SYTOX Orange, gpD-EYFP phages were used to infect MG1655 harboring pALA3047 (CFP-ParB).

##### *5.2. Infection*

Infection mixtures comprising 10 μL of cells and 10 μL of phages (with phage-to-bacteria ratio ≈ 3), and a negative control comprising cells and blank solution were prepared. In experiments where the phage-to-bacteria ratio was varied **(Figure 1B)**, phages at different concentrations were used but the volume ratio between cells and phages remained constant. The samples were cooled at 4°C for 30 minutes in a dry block incubator (ThermoFisher/VWR) to allow phages to adsorb to cells, then heated at 35°C for 5 minutes to trigger phage ejection. Using wide pipette tips, the samples were gently diluted 1:10 to decrease the cell density before imaging. During this step, if the infection medium was LBMM or TBM, we used PBSM for dilution to reduce the autofluorescence background (*14*). Otherwise, the same solution as the infection step was used for dilution.

##### *5.3. Sample mounting and imaging*

Samples were mounted for imaging as described in (*15*). Briefly, a 1.5% agarose pad (made using the same solution as the one used to dilute the infection mixture) was laid on top of a 22×22 mm coverslip no. 1 (Fisher Scientific). Then, 1 μL of the diluted sample was gently pipetted onto the pad. When the 1 μL droplet was no longer visible (1–2 minutes), a 24×50 mm coverslip no. 1 (Fisher Scientific) was gently laid on top of the pad. Samples were imaged as described in **Section 11**, followed by analysis (described in **Section 12**) to obtain the numbers of adsorbed phages and intracellular phage genomes for each cell.

##### *5.4. Treatment with carbonyl cyanide m-chlorophenyl hydrazone (CCCP)*

In experiments where the efficiency of phage entry in depolarized cells was examined (**Figure 3H**), cell washing and resuspension, phage dilution, and infection were performed using LBMM supplemented 200 μM of CCCP (similar concentration to that in (*10*)). This CCCP treatment protocol was validated using a control experiment with PROPS-expressing cells (**Section 10**). A no-treatment control was prepared using LBMM supplemented with 0.5% dimethylsulfoxide (DMSO). Following infection, the samples were diluted, mounted, and imaged as described above. Consistent with (*16*), entry of lambda phages still occurred in cells treated with CCCP, but the efficiency of phage entry was severely reduced (**Figure 3H**).

##### *5.5. Data analysis*

We used the stochastic model of phage entry kinetics (derived in **Section 18** below; see **Table S5** for variable definitions) to capture the sublinear relation between the numbers of intracellular and adsorbed phages following bulk infection in LBMM. To do so, we used the parameters $(\eta_n, k_n, \tau_n)$ calibrated using the microfluidic assay in LBMM (**Section 8.3.3**) and **Equation 18.3** to calculate $\langle \lambda(n, t) \rangle$. To account for the additional time of sample dilution and mounting before the microscopy images were taken, we scanned for $t$ between 5 and 12 minutes at a 0.1-minute increment. At each time point, the root-mean-squared-error (RMSE) between the model prediction and the data was calculated. The time at which RMSE was minimized was 9.1 minutes (**Figure S14**).

This model prediction was possible because the kinetic parameters were already inferred from microfluidic infection data in LBMM (**Section 8**). When kinetic parameters were not available (e.g., in other infection media surveyed in **Figure S23**), we used the following phenomenological expression to capture the average number of intracellular phage genomes:

$$\langle \lambda(n) \rangle = \frac{a \cdot n}{K + n}. \tag{5.1}$$

**Equation 5.1** shares the same form with the Michaelis-Menten function, which is also a Hill function with a coefficient of 1. The fitted parameter values for each infection medium are given in the caption of **Figure S23**.

To capture the asymptotic value of entry efficiency in cells adsorbed by many phages (**Figure 3H**, negative control in the CCCP experiment), we used the following expression:

$$\langle \lambda / n \rangle = a \cdot e^{-b(n-1)} + c\,, \tag{5.2}$$

which describes an exponential decay with a baseline. This expression provides the entry efficiency in cells adsorbed by one phage ($= a + c$ when $n = 1$) and in cells adsorbed by many phages ($= c$ when $n \to \infty$).

To capture the distribution of intracellular phage numbers (e.g., **Figure 4C**), we used the following expression:

$$P(\lambda) = \left( \sum_{j=0}^{n} \frac{\mu^j e^{-\mu}}{j!} \right)^{-1} \frac{\mu^\lambda e^{-\mu}}{\lambda!}. \tag{5.3}$$

**Equation 5.3** describes a truncated Poisson distribution for the random variable $\lambda$, controlled by a parameter $\mu$. Truncation is necessary because there cannot be more intracellular phage genomes ($\lambda$) than the number of adsorbed phages ($n$).

In **Figures S31–S33**, we show that **Equations 5.1–5.3** capture the corresponding predictions by the stochastic model over the experimentally relevant range of MOI and time.

##### *5.6. Fitting procedure*

Fitting to single-cell data was performed with bootstrapping (1000 iterations). In each iteration, a bootstrapped sample with the same number of cells as the original dataset was randomly drawn with replacement (implemented using the MATLAB \`datasample\` function). Fitting to each bootstrapped sample was implemented with the MATLAB \`fit\` function, using nonlinear least-squares optimization and the trust-region algorithm. Estimates and SE of the parameters were calculated from the mean and standard deviation (SD) of the fitted parameters' sampling distributions.

#### 6. Measuring phage-induced membrane permeabilization using propidium iodide

##### *6.1. Infection, staining, and imaging*

MG1655 cells harboring pALA3047 (expressing CFP-ParB) were grown and concentrated as described in **Section 5**. An infection mixture, using $\lambda_{TY11}$ labeled with gpD-EYFP, was prepared at a phage-to-bacteria ratio of approx. 3, incubated for 30 minutes at 4°C, then shifted to 37°C. At each time point (5, 20, 35, and 50 min), an aliquot of the infection mixture was taken, diluted 1:10 in PBSM, and mixed with propidium iodide (PI, final concentration 10 μM, following (*17*)). The sample was incubated in the dark at RT for 5 minutes, then mounted and imaged as described in **Section 11**. Image analysis was performed as described in **Section 12** to obtain the number of adsorbed phages, the number of intracellular phage genomes, and the intracellular PI fluorescence of each cell.

##### *6.2. Controls*

Because PI was previously reported to perturb productive phage infection (*18*), here, we only added PI to an aliquot of the infection mixture before imaging, while phage adsorption and entry both took place in the absence of PI. As a result, phage entry was not perturbed by PI. Indeed, the efficiency of phage entry in PI-stained samples was similar to that in the unstained sample, and the sublinear relation between the numbers of intracellular and adsorbed phages was reproduced (**Figure S15A**).

For the negative control, cells that had gone through the same centrifugation and washing protocol as the infected sample were mixed with blank medium, incubated at 4°C for 30 minutes, and shifted to 37°C for 60 minutes (longer than the last time point of the infected sample). The sample was then stained with PI and imaged. The PI fluorescence in the negative control was similar to that in cells with MOI = 0 in the infected sample (**Figure S15B**).

For the positive control, cells were permeabilized by incubation with 70% ethanol at RT for 30 minutes (*19*), then washed and resuspended in PBS. Permeabilized cells were stained with PI and imaged. The PI fluorescence in ethanol-treated cells was approx. 50× higher than that in cells with MOI = 10 (**Figure S15B**). This suggested that the PI permeation observed in the infected sample did not reflect cell death (in contrast to, e.g., (*20*)). In addition, since the phage strain used in this experiment, $\lambda_{TY11}$, is incapable of the lytic pathway (*1*, *21*), permeation of PI into the cell, even at 50 minutes after infection, was not due to the onset of cell lysis (in contrast to, e.g., (*17*)).

##### *6.3. Data analysis*

We fitted the following expression to the intracellular PI fluorescence ($F$) as a function of the number of adsorbed phages ($n$):

$$F(n) = a \cdot n \,. \tag{6.1}$$

In **Equation 6.1**, $F(n = 0) = 0$ reflects our image analysis procedure (described in **Section 12**), in which the PI fluorescence in all cells was corrected by that in cells with MOI = 0. The slope $a$ provides an estimate for the degree of membrane permeabilization per infecting phage. **Equation 6.1** was fitted to data at each time point (5, 20, 35, and 50 minutes), with bootstrapping as described in **Section 5.6**. The fitting results are shown in **Figure 3B** and **Figure S16A.** The values of the slope $a$ are shown in **Figure 3C**. A similar fitting procedure was performed for the intracellular PI fluorescence as a function of the number of intracellular phage genomes, $F(\lambda)$, shown in **Figure S16B**.

Calculations of the Pearson correlation coefficients between the PI fluorescence and the number of adsorbed phages, and between the PI fluorescence and the number of intracellular phage genomes were performed with bootstrapping (1000 iterations, implemented using the MATLAB `corrcoef` function). We found that the PI fluorescence was positively correlated with the number of adsorbed phages and was not correlated with the number of intracellular phage genomes (**Figure S16C**).

We fitted the following expression to the efficiency of phage entry as a function of the PI fluorescence:

$$\langle \lambda / n \rangle = a \cdot e^{-bF} + c \,. \tag{6.2}$$

**Equation 6.2** provides an estimate of the entry efficiency in non-permeabilized cells ($= a + c$ when $F = 0$) and the asymptotic value in highly permeabilized cells ($= c$ when $F \to \infty$). **Equation 6.2** was fitted to the pooled data from 5 and 20 minutes with bootstrapping as described in **Section 5.6**. Fitting results are shown in **Figure 3D**, and the fitted parameters are $a = 0.45 \pm 0.05$, $b = 0.004 \pm 0.002$, and $c = 0.26 \pm 0.08$.

##### *6.4. Causal inference*

Because the number of adsorbed phages is negatively correlated with the efficiency of phage entry (**Figure 1C**) and positively correlated with the intracellular PI fluorescence (**Figure 3B**), the observed correlation between the entry efficiency and the PI fluorescence (**Figure 3D**) could be due to the confounding effect of MOI. To control for MOI and infer whether there is a causal link between entry efficiency and PI fluorescence, we followed (*22*) and calculated the conditional correlation, $r$, between entry efficiency and PI fluorescence when the effect of MOI is removed.

Using the pooled data from 5 and 20 minutes, we performed linear regressions of entry efficiency on MOI, and of PI fluorescence on MOI (**Figure S17**). Cells with no adsorbed phages, in which the efficiency of phage entry is undefined, were not included in this analysis. The residuals following linear regressions represent variations in the entry efficiency and in the PI fluorescence for a given MOI, i.e., due to sources other than MOI. These residuals were denoted as $\text{Entry efficiency}|\text{MOI}$ and $\text{PI fluorescence}|\text{MOI}$. If there is no causal link between PI fluorescence and entry efficiency, the correlation between these residuals will be zero. However, we found that the Pearson correlation coefficient between $\text{Entry efficiency}|\text{MOI}$ and $\text{PI fluorescence}|\text{MOI}$ is negative (**Figure 3E**). Calculation of $r(\text{Entry efficiency}, \text{PI fluorescence}|\text{MOI})$ was performed with bootstrapping (1000 iterations), yielding $r = -0.27 \pm 0.04$. Performing the Student's t-test, we found that this negative correlation is statistically significant ($p\text{-value} \approx 5 \times 10^{-6}$). Therefore, compromise to membrane integrity was inferred to be a cause of impeded phage entry.

To investigate the presence of additional mechanisms connecting phage adsorption and impeded entry, we performed linear regression of entry efficiency on PI fluorescence, and of MOI on PI fluorescence (**Figure S18**). The correlation coefficient between the residuals $\text{Entry efficiency}|\text{PI fluorescence}$ and $\text{MOI}|\text{PI fluorescence}$ is also negative, $r(\text{Entry efficiency}, \text{MOI}|\text{PI fluorescence}) = -0.19 \pm 0.05$ (SE from bootstrapping, $p\text{-value} \approx 8 \times 10^{-4}$). This suggests that among cells with the same degree of membrane permeabilization, the efficiency of phage entry is still lower in cells with more adsorbed phages. Thus, we could not rule out the presence of other mechanisms connecting phage adsorption and impeded entry.

#### 7. Measuring the time of the first phage entry following bulk infection

##### *7.1. Infection*

MG1655 cells harboring pALA3047 (expressing CFP-ParB) were grown and concentrated as described in **Section 5**. An infection mixture was prepared at a phage-to-bacteria ratio of approx. 0.5 and incubated at 4°C for 30 minutes. To trigger phage ejection and to prevent additional adsorptions, the infection mixture was diluted 1:1000 into prewarmed LBGM supplemented with IPTG in a baffled Erlenmeyer flask. The diluted infection mixture was shaken at 220 rpm in a 37°C water bath for 10 minutes, during which samples were extracted for fixation. A negative control (cells and blank solution) was also prepared and extracted at the end of the experiment.

##### *7.2. Fixation and imaging*

Chemical fixation was performed as described in (*1*). Briefly, at each time point (immediately before the dilution to 37°C, then at 1, 2, 3, 4, 5, and 10 minutes), samples were aliquoted and mixed with formaldehyde (final concentration 3.7%) in PBS (final concentration 1×) for 30 minutes at RT. The samples were washed twice and concentrated in PBS, mounted as in (*19*), and imaged as described in **Section 11**. Image analysis was performed as described in **Section 12** to obtain the number of intracellular phage genomes in each cell.

##### *7.3. Data analysis*

At this phage-to-bacteria ratio (≈ 0.5), we assumed that all cells in the infection mixture were adsorbed by either 0 or 1 phage (*23*). Hence, measurements obtained from the infected cells reflected the kinetics of phage entry in cells adsorbed by one phage. We fitted the following expression to the time-dependent average number of intracellular phage genomes among all cells:

$$\langle \lambda(t) \rangle = a\left(1 - e^{-k(t-\tau)}\right) \text{ for } t \geq \tau, \text{and 0 otherwise.} \quad (7.1)$$

**Equation 7.1** was derived using the stochastic model (see **Equation 18.3** in **Section 18** below). Here, the parameter $a$ describes the fraction of cells with intracellular phage genomes among all cells in the infection mixture, thus representing both the entry ($\eta$) and adsorption efficiencies. Fitting was performed with bootstrapping as described in **Section 5.6**. Both the data and the fit in **Figure S8** are shown after rescaling, i.e., $\langle {}^{1}/_{a} \cdot \lambda(t) \rangle$ vs. $t$. The measured kinetics from this bulk assay is compared with that in the microfluidic assay and the literature in **Figure S4**.

#### 8. Measuring the kinetics of phage entries in the microfluidic device

##### *8.1. Preparation of cells and phages*

Cultures of MG1655 harboring p2973 or pALA3047 were grown at 37°C in LBMM supplemented with IPTG and 100 µg/mL ampicillin as described in **Section 4**. Upon reaching $OD_{600} \approx 0.3$–0.4, the culture was diluted 5× in LBMM supplemented with IPTG, and introduced into the microfluidic device as described below. The purified phage stock ($\lambda_{TY11}$ with fluorescently labeled capsids, described in **Section 3**) was diluted to approx. $2\times10^{10}$ PFU/mL using LBMM supplemented with IPTG. In this assay, gpD-mTurquoise2 or gpD-EYFP phages were used to infect MG1655 harboring p2973 or pALA3047 (expressing mCherry-ParB or CFP-ParB), respectively.

##### *8.2. Setting up and operating the microfluidic device*

Microfluidic experiments were performed in B04A plates of the CellASIC ONIX or ONIX2 microfluidics system (MilliporeSigma), controlled using the ONIX FG software. The microfluidic system was incubated in a temperature-controlled enclosure (Okolab), set at 37°C. Cells, phages, and blank medium (LBMM supplemented with IPTG) were pipetted into inlets of a microfluidic plate (pre-purged as described in the instruction manual). The observation chamber was first primed with the blank medium. Cells were then loaded into the chamber and grown for 30 minutes (approx. one generation). Fresh medium was constantly exchanged during this time using a flow pressure of 1 psi. To begin infection, phages were perfused into the chamber at 10 psi for 2 minutes, followed by the blank medium at 10 psi for 4 minutes to wash out excess phages. Constant medium exchange at 1 psi was resumed until the end of the experiment. Image acquisition was performed throughout this perfusion protocol (described in **Section 11**). Image analysis was performed as described in **Section 12** to obtain the numbers of adsorbed phages and intracellular phage genomes for each cell at every time point. The time series and the average kinetics of phage adsorptions and entries from a representative experiment are shown in **Figure S7**.

In this assay, the phage concentration and flow time were chosen to obtain a countable number of adsorbed phages per cell while maximizing synchrony in adsorption time. The average numbers of adsorbed phages per cell, as a function of phage concentration and flow time, are shown in **Figure S6**.

##### *8.3. Data analysis*

In the following analyses (**Sections 8.3.1–8.3.3**), the expressions used for model fitting are derived in **Section 18**, and the variables are defined in **Table S5**. Model fitting was implemented with the MATLAB `lsqcurvefit` function, using nonlinear least-squares optimization and the trust-region algorithm. Fitting was performed with bootstrapping (1000 iterations for each single-cell dataset), as described in **Section 5.6**.

###### *8.3.1. Estimation of model parameters for MOI = 1*

We first examined the entry kinetics in cells with one adsorbed phage (7 experiments acquired at 1-minute imaging frequency, each yielding 6–49 cells with $n = 1$). For each experiment, we calculated the time-dependent average number of intracellular phages in cells adsorbed by one phage, $\langle\lambda(n=1,t\rangle$, and fitted **Equation 18.34** to this data. The inferred parameters are $\eta_1 = 0.56 \pm 0.04$, $k_1 = 0.011 \pm 0.004\ \mathrm{s}^{-1}$, and $\tau_1 = 43.0 \pm 10.5\ s$ (Mean and SE of the fitted parameters of the 7 experiments; the subscript "1" denotes parameters for $n = 1$). Using the pooled dataset from all 7 experiments yielded similar values: $\eta_1 = 0.56 \pm 0.04$, $k_1 = 0.010 \pm 0.001\ s^{-1}$, and $\tau_1 = 35.9 \pm 5.6\ s$ (SE from bootstrapping); this fitting result is shown in **Figure 2C**. The inset of **Figure 2C** shows $\log(1 - \langle\lambda\rangle/\eta)$ vs. $t - \tau$; the linearity of this graph supports the analytical expression in **Equation 18.34**. The inferred parameters are compared with those obtained in the bulk assay and the literature in **Figure S4**.

We next used **Equation 18.31** and the fitted parameters $(\eta_1, k_1, \tau_1)$ above to calculate the probability density of phage entry time, $f(T_1|n=1)$. To compare with the experimental data (**Figure 2D**), we binned the predicted probability density by the imaging inter-frame interval (calculated using the MATLAB `trapz` function):

$$P(T_1 = t|n = 1) = \int_{t-1}^{t} f(T_1 = t'|n = 1)\, dt' \,, \tag{8.1}$$

*8.3.2. Estimation of model parameters for MOI > 1*

Next, we assumed that the parameters fitted using cells with $n = 1$ were also applicable to cells with $n > 1$. Following this assumption (termed the MOI-independent model), we used **Equation 18.3** and the parameters $(\eta_1, k_1, \tau_1)$ from **Section 8.3.1.** to predict the time-dependent average number of intracellular phage genomes in cells adsorbed by multiple phages, $\langle\lambda(n,t)\rangle$, and compared these predictions with the data for $n \in [2,10]$ (beyond which the number of cells for a given MOI is less than 5). As $n$ increases, the model prediction becomes poorer (**Figures 2E** and **S10**). In particular, the root-mean-squared-error (RMSE) between the data and the model increases with $n$ (**Figure S9**). We concluded that the MOI-independent model was inappropriate, and the parameters for $n = 1$ cannot be applied to $n > 1$.

Instead, we allowed all three parameters of the model to vary with MOI. To do so, we divided the pooled dataset into subsets of cells with different $n$, and fitted **Equation 18.3** to each of these data subsets, yielding separate sets of parameters $(\eta_n, k_n, \tau_n)$ for each $n$ value. This model (termed the MOI-dependent $\eta, k, \tau$ model), captured the time-dependent average number of intracellular phages well (**Figure S10**). The average RMSE of the MOI-dependent $\eta, k, \tau$ model is 0.0862, a considerable improvement compared to the MOI-independent model with RMSE = 0.887 (**Figure S9**). Examining how each parameter varies with $n$, we noted that both the fitted $\eta_n$ and $k_n$ appeared to decrease with $n$, whereas $\tau_n$ showed no consistent trend (**Figure S11**). We hypothesized that a dependence of $\eta$ and $k$ on the MOI ($n$) is necessary to capture the data, while $\tau$ can remain MOI-independent.

To test this hypothesis, we fitted a model in which only $\eta$ and $k$ were allowed to vary with $n$, while $\tau$ was constrained at $\tau_1$, and obtained the parameters $(\eta_n, k_n, \tau_1)$ for each $n$ value. Predictions by this model (termed the MOI-dependent $\eta, k$ model) were indistinguishable from those of the MOI-dependent $\eta, k, \tau$ model (**Figure S10**). The average RMSE of the MOI-dependent $\eta, k$ model (= 0.0825) is lower than that of the MOI-dependent $\eta, k, \tau$ model **(Figure S9**). When we investigated the other 5 models in which one or two of the parameters were allowed to vary with the MOI, none of them achieved a lower RMSE than when only $\eta$ and $k$ were allowed to vary (**Figure S9**). Therefore, we concluded that the MOI-dependent $\eta, k$ model is the most appropriate one to describe the observed data.

*8.3.3. Parametrization of the MOI-dependent model*

To describe the overall behaviors of $\eta$ and $k$ as a function of $n$, and to infer parameter values when data of a given $n$ was not available, we parametrized $\eta_n$ and $k_n$ as exponentially decaying functions of $n$:

$$\eta_n = a_\eta \cdot e^{-b_\eta(n-1)} + c_\eta \,, \tag{8.2}$$

$$k_n = a_k \cdot e^{-b_k(n-1)} + c_k \,. \tag{8.3}$$

The fitted values of $\eta$ and $k$ from the MOI-dependent $\eta, k$ model above (**Section 8.3.2.**) were used to fit **Equations 8.2** and **8.3**, respectively. Fitting was weighted by the natural logarithm of the number of cells with a given $n$. The results of this parametrization are shown in **Figure 2F.** For $\eta$, $a_\eta = 0.458$, $b_\eta = 0.0762$, and $c_\eta = 0.131$. For $k$, $a_k = 0.00450$, $b_k = 0.456$, and $c_k = 0.00479$. As for $\tau$, we simply set $\tau_n = \tau_1 = 35.9 \pm 5.6$ s. This set of inferred parameters $(\eta_n, k_n, \tau_n)$ constitutes the MOI-dependent model of entry dynamics, used below and in **Sections 19** and **20**.

To predict the distribution of intracellular phage numbers at a given time after adsorption, we used **Equation 18.29** and these inferred parameters $(\eta_n, k_n, \tau_n)$ to calculate $P(\lambda|n, t)$. These predictions are shown in **Figures 2G** and **S12** for $t \in [0,10]$ min and $n \in [1,4]$ (beyond which the sample size is smaller than 30 cells).

We also used the inferred parameters to predict the relation between the numbers of adsorbed and intracellular phages at different times. This was calculated using **Equation 18.3** and plotted as $\langle\lambda(n)\rangle$ separately for each time point $t \in [0,10]$ min. These predictions are shown in **Figures 2H** and **S13**. Using the same model parameters, we also captured the values observed in bulk infection assays (**Figure S14**, described above in **Section 5**).

#### 9. Validating the ParB-*parS* labeling system using SYTOX Orange

##### *9.1. Experimental protocol*

To measure the detection efficiency of the ParB-*parS* system (**Figure S2**), we used SYTOX Orange to stain the DNA of phages whose capsids are labeled with gpD-EYFP (produced as described in **Section 3**), and infected cells expressing CFP-ParB. The microfluidic infection protocol was performed as described in **Section 8**. Imaging was performed at the end of the experiment (described in **Section 11)**, followed by image analysis as described in **Section 12** to obtain the numbers of adsorbed phage capsids, encapsidated phage DNA, and intracellular phage genomes in individual cells.

##### *9.2. Data analysis*

An entry event was defined to have occurred for every phage capsid without encapsidated DNA (i.e., gpD-YFP foci without SYTOX Orange signal). If the ParB-*parS* system faithfully detects intracellular phage genomes, there will be a ParB-*parS* spot for every corresponding entry event. In other words, we predicted the following relation:

$$N_{\text{ParB-}parS} = N_{\text{fluorescent capsids}} - N_{\text{SYTOX Orange}}, \tag{9.1}$$

in which $N_i$ is the number of fluorescent spots of type $i$ in each cell. The fitted slope of $N_{\text{ParB-}parS}$ vs. $N_{\text{fluorescent capsids}} - N_{\text{SYTOX Orange}}$ is 0.89 ± 0.04 (**Figure S2**, SE from bootstrapping as described in **Section 5.6**). This suggests the detection efficiency of ParB-*parS* is approx. 90%.

##### *9.3. Correcting for SYTOX Orange labeling*

If SYTOX Orange fails to stain the encapsidated phage DNA, the capsid would appear to be empty regardless of whether ejection has occurred. As a result, an entry event would be falsely registered, and the detection efficiency of ParB-*parS* would be underestimated.

To correct this underestimation, we performed a simple stochastic simulation as follows. For each cell, given $N_{\text{fluorescent capsids}}$, we generated a random number from the following binomial distribution (implemented using the MATLAB `binornd` function):

$$N_{\text{stained particles}} \sim \text{Binom}\left(N_{\text{fluorescent capsids}}, \eta_{\text{SYTOX Orange}}\right), \tag{9.2}$$

in which $\eta_{\text{SYTOX Orange}}$ is the labeling efficiency of SYTOX Orange (91.9% ± 1.6%, measured as described in **Section 3**). For each cell, $N_{\text{stained particles}}$ simulates the number of phage particles that were stained with SYTOX Orange prior to infection. This simulation was performed for all cells in the original dataset, yielding a simulated dataset of the same size. The fitted slope of $N_{\text{ParB-}parS}$ vs. $N_{\text{stained particles}} - N_{\text{SYTOX Orange}}$ provided a corrected estimate of the detection efficiency of the ParB-*parS* system. The distribution of this simulated slope value ($N$ = 1000 realizations) is shown in **Figure S2**. The mean and SD of this distribution is 0.93 ± 0.02, suggesting the detection efficiency of the ParB-*parS* system was approx. 93%.

#### 10. Validating the CCCP treatment protocol using PROPS

Our protocol for carbonyl cyanide *m*-chlorophenyl hydrazone (CCCP) treatment (**Sections 5** and **13**) was validated using PROPS-expressing cells. Cultures of MG1655 pJMK001 were grown as described in **Section 4**. Following PROPS induction, cells were washed and concentrated 5× in LBM supplemented with either 200 µM CCCP or 0.5% DMSO (the same residual concentration of DMSO in the CCCP-treated sample, serving as a negative control). Samples were mounted and imaged as described in **Section 11**, and the PROPS fluorescence was quantified as described in **Section 12**.

The PROPS fluorescence in cells depolarized using CCCP was approx. 3× higher than that in the DMSO control (**Figure S19**). This fold-change is in agreement with (*10*), thus validating our protocol for CCCP treatment.

#### 11. Microscopy

##### *11.1. Equipment and setup*

For all experiments except for those involving the optical traps (**Section 13**), either one of the following two inverted epifluorescence microscopes was used. The first one is an Eclipse T*i* (Nikon) system, equipped with a mercury lamp (Intensilight C-HGFIE, Nikon), a CMOS camera (Prime 95B, Photometrics), and a 100×, NA 1.45, oil-immersion phase-contrast objective lens (Plan Apo, Nikon). The second one is an Eclipse T*i*2 (Nikon) system equipped with an LED light source (X-Cite XYLIS); the camera and the objective lens were identical to the first setup. The microscopes were installed on a pneumatically-supported vibration-isolation table (CleanBench) and placed in a temperature-controlled enclosure (Okolab). Microscope operation and image acquisition were performed using the NIS-Elements software (Nikon). For phase contrast imaging, we used an exposure time of 100 ms. For fluorescence imaging, we used the filter sets listed in **Table S4**, with exposure times ranging between 50 ms and 400 ms.

##### *11.2. Snapshot imaging*

Samples were mounted using coverslips and an agarose pad (described in **Sections 5**–**7** and **10**), and placed on a universal specimen holder (TI2-S-HU, Nikon). Images were acquired at multiple fields of view (*xy*-positions), located at least 300 µm apart to minimize photobleaching across fields of view. The different fields of view were located using motorized stage control. For each field of view, the phase contrast channel was imaged first, followed by the fluorescence channels. For each channel, images were taken at 7 focal planes (*z*-slices), with steps of 300 nm apart.

##### *11.3. Time-lapse imaging*

CellASIC ONIX B04A plates were prepared as described in **Sections 8** and **9** and placed on a well-plate holder (TI2-S-HW, Nikon). The imaging frequency was either 1 minute (most experiments), 2 minutes, or 10 minutes (pilot or control experiments). For each time point, all of the fields of view were imaged. For each field of view, all of the phase contrast and fluorescence channels were imaged. For each channel, images were taken at 3 focal planes, with steps of 400 nm apart. Image acquisition was initiated at least 1 minute before phage perfusion and terminated no earlier than 20 minutes afterwards.

#### 12. Image analysis

Analysis was performed on the original images (before contrast adjustment was performed to produce representative images, as described in **Section 14**), using the Nikon NIS-Elements software, the ImageJ2/Fiji software, and custom MATLAB (MathWorks) codes. In all analyses, the phase contrast channel provides information regarding the outline, size, and morphology of bacterial cells.

##### *12.1. Measuring single-cell MOI*

To quantify the numbers of extracellular phages (using DAPI, gpD-mTurquoise2, gpD-EYFP, or SYTOX Orange) and intracellular phage genomes (using CFP-ParB or mCherry-ParB), we manually counted the number of diffraction-limited fluorescent foci (spots) on the cell surface or inside the cell, respectively, in the corresponding fluorescence channels.

In time-lapse assays, for each cell tracked over the course of the experiment, the numbers of adsorbed phages and of intracellular phage genomes were recorded at each time point. Using this time series with respect to the beginning of phage perfusion (**Figure S7**), the times of phage adsorptions and entries were inferred. For example, if the numbers of adsorbed phages at (0, 1, 2, 3, 4,…) minutes were (0, 0, 1, 3, 3,…), the 1$^{st}$ adsorption event was considered to have occurred between 1 and 2 minutes, while the 2$^{nd}$ and 3$^{rd}$ adsorptions both took place between 2 and 3 minutes. Only cells with synchronized phage adsorptions—defined to be those with the first and last adsorption events within 2 minutes of one another—were used for model fitting. For each cell, the average time at which phages adsorbed to the cell was calculated. The time-dependent numbers of intracellular phage genomes and the entry times of individual phages were calculated with respect to this cell-specific average adsorption time (i.e., time since phage adsorption, set to be $t = 0$).

##### *12.2. Measuring propidium iodide (PI) fluorescence*

Using phase contrast, we performed manual segmentation to identify a region of interest (ROI) corresponding to each cell and measured the average fluorescence of PI (in arbitrary units, A.U., per pixel) within the ROI. The PI fluorescence was corrected for the average background outside the cells in the same field of view and for the average intensity in cells with MOI = 0. Cells with overwhelming PI fluorescence (< 0.5% of the dataset, comparable to that in ethanol-treated cells as measured in **Section 6.2**) were excluded from the analysis.

##### *12.3. Measuring PROPS fluorescence*

PROPS fluorescence in individual cells was quantified in the same manner as PI, as described above. PROPS fluorescence was corrected for the background outside the cell in the same field of view, then normalized using the average intensity of cells in the negative control. Hence, the PROPS fluorescence following CCCP treatment or phage infection was reported in fold-change, as in (*10*).

#### 13. Optical trap assay

##### *13.1. Preparation of cells and phages*

For this assay, we followed the protocol from (*24*). Briefly, cultures of MG1655 or MG1655 harboring pJMK001 (expressing PROPS) were grown as described in **Section 4**. Cells were centrifuged at 1300×g for 10 minutes at RT, resuspended in the same volume in MB, and diluted 5× in TMB or TMBM as specified below (See **Table S3** for the composition of these buffers). When required, phages were stained with SYTOX Orange as described in **Section 3**.

##### *13.2. Setup of the optical traps and the flow chamber*

All trap-related experiments were performed at room temperature. The optical traps and epifluorescence setup were described in (*25*). The three-channel flow chamber (**Figure 3F**) was made as described in . Solutions were perfused at a flow rate of 0.33 µL/min, resulting in a linear speed of 35 µm/s. The positions of the optical traps were recorded at 666 Hz, and the fluorescence of SYTOX Orange or PROPS was imaged using a 532 nm excitation laser.

##### *13.3. Measuring the membrane potential following CCCP treatment*

In this experiment, the three channels of the flow chamber contained (i) PROPS-expressing cells in TMB, (ii) blank TMB, and (iii) TMB supplemented with 200 µM CCCP. Individual cells were first trapped from the cell channel, then moved into the blank channel. Each cell was kept in the blank channel for approx. 1 minute to record the basal flagellar rotation frequency, then moved into the CCCP channel, where the loss of motility and the increase in PROPS fluorescence were observed within ~5 seconds (**Figure S20**).

##### *13.4. Measuring the membrane potential following phage adsorption*

In this experiment, the three channels of the flow chamber contained (i) cells in TMBM, (ii) blank TMBM, and (iii) phages at approx. $1\times10^{10}$ PFU/mL in TMBM. Two combinations of phages and cells were used: (i) Phages stained with SYTOX Orange, infecting MG1655 cells; and (ii) Unstained phages, infecting PROPS-expressing cells.

Individual cells were first trapped from the cell channel, then moved into the blank channel for approx. 1 minute to record the trap positions. Each cell was then moved into the phage channel for at least 5 minutes. During this time, if a phage (visualized using SYTOX Orange) stably adsorbed to the cell, the cell was considered to be infected and was moved back into the blank channel. Cells with no adsorbed phage were used as a negative control. Cells (with MOI = 0 or 1) were monitored in the blank channel for at least 30 minutes or until motility was lost. In this experiment, $t$ = 0 was defined to be the time the phage adsorbed to the cell, or when a cell with no adsorbed phage was moved back into the blank channel.

In experiments with PROPS-expressing cells, cells were kept in the phage channel for at least 30 minutes or until motility was lost. In this case, $t$ = 0 was defined to be the time the cell was moved into the phage channel. Because phages were not stained in this experiment, the adsorption time and the MOI could not be measured.

##### *13.5. Data analysis*

To extract the flagellar rotation frequency, the trap positions were analyzed using wavelet analysis as described in (*13*). A typical cell in this assay had a flagellar rotation frequency of approx. 100 Hz. A cell was determined to have lost motility, indicating membrane depolarization, when the flagellar rotation frequency decreased to zero.

**Figure 3G** shows a "survival curve" for the fraction of motile cells over time following phage adsorption, $f_{\mathrm{motile}}(t)$. Data within the first 10 minutes was used to fit the following expression:

$$f_{\mathrm{motile}}(t) = e^{-\kappa t}. \tag{13.1}$$

To estimate $\kappa$, fitting of **Equation 13.1** to data was performed with bootstrapping as described in **Section 5.6**. The fitting result is shown in **Figure 3G**, with $\kappa = 0.12 \pm 0.05\ \mathrm{min}^{-1}$. The half-life, $t_{1/2} = \log 2/\kappa = 5.6 \pm 2.2\ \mathrm{min}$, describes the time it takes for 50% of the cells to lose motility.

In experiments involving PROPS, the PROPS fluorescence was corrected for the background fluorescence, then normalized using the basal fluorescence prior to CCCP treatment or phage adsorption. Hence, the PROPS fluorescence was reported in fold-change, as in (*10*) and **Section 12**.

#### 14. Preparation of representative images

To prepare representative images (**Figures 1A**, **2A**, **3A**, **3F**, **S1**, **S2**, and **S19**), we used Nikon NIS-Elements and ImageJ2/Fiji, to first perform a maximum intensity projection across the focal planes. Then, to remove background and non-specific fluorescence, we applied contrast adjustments to the entire image and to all samples. Pseudo-coloring of the fluorescence channels was then performed. Extracellular phage capsids (gpD-mTurquoise2 or gpD-EYFP) were pseudo-colored in cyan, and intracellular phage genomes (mCherry-ParB or CFP-ParB) were pseudo-colored in red. DAPI, SYTOX Orange, PI, and PROPS were shown in other colors. When appropriate, the relevant channels were combined into a single image, and cropped to highlight the cell or area of interest. Scale bars were provided for all images (80 nm per pixel for images obtained with the optical traps, and 44 nm per pixel otherwise).

#### 15. Potassium efflux assay

For this assay, we followed the protocols from (*18*, *26*). Cultures of MG1655 were grown at 37°C in LBMM as described in **Section 4**. When the culture reached $OD_{600} \approx 0.3–0.4$, cells were centrifuged, washed once in SM3 buffer (described in (*18*)), and resuspended in SM3 at the original concentration. The cell suspension was incubated at 37°C for at least 5 minutes before infection was performed. The purified phage stock (**Section 3**), diluted to $1\times10^{10}$ PFU/mL in SM3, was added to the cells at a phage-to-bacteria ratio of approx. 100. The concentration of $K^+$ ions in the medium was measured using an Orion $K^+$ ISE Electrode (ThermoScientific) and a FiveEasy pH/mV meter (Mettler Toledo). The voltage readings were manually recorded once every 10 seconds from 5 minutes before phage addition (the baseline) to 15 minutes after. Voltage readings were converted to molarity using a calibration curve, obtained using $K^+$ standard solutions as prescribed by the electrode's instruction manual. The total $K^+$ content in the cells was determined by treating the same amount of cells with the BugBuster 10× Reagent (Millipore). This value was used for normalization to calculate the time-dependent fraction of $K^+$ released due to phage infection (**Figure S21**).

#### 16. Measuring the frequency of lysogeny following infection in different media

##### *16.1. Infection and selection*

This assay was an adaptation of (*1*), but instead of using agar plates, we used a plate reader to count the number of lysogens. MG1655 cells, grown as described in **Section 4**, were centrifuged, washed in LBM or SM, and concentrated 20× in LBM or SM (similar to **Section 5**). Two 2.5× dilution series of the phage stock (produced as described in **Section 3**) were prepared in LBM or SM (from $\sim7.5\times10^7$ to $\sim1.2\times10^{11}$ PFU/mL). Two infection series in LBM and SM (phage-to-bacteria ratio ranging from ~0.05 to ~50) were prepared, incubated at 4°C for 30 minutes, and shifted to 35°C for 5 minutes. Then, 2 µL of each infection mixture was diluted into duplicate wells containing 500 µL LBM, in a clear 48-well flat-bottom microplate (COSTAR). Samples were incubated for 1 hour at 30°C with shaking (orbital mode, 1 mm amplitude) in a TECAN F200 Pro plate reader to allow the lysogenic cells to express kanamycin resistance. Each culture was then supplemented with 50 µg/mL kanamycin, and incubation with shaking was resumed. For each infection series, an uninfected control without kanamycin selection was also prepared. The plate reader was set to measure the optical density (OD) of the cultures once every 5 minutes for ~24 hours.

Our SM infection procedure is different from protocols involving pre-infection starvation (**Figure S24**), which were reported to result in a higher frequency of lysogeny (*23*, *27–29*). In these studies, starvation prior to infection was achieved by either growing cells into stationary phase (*23*, *29*) or by incubating exponentially growing cells in a solution without nutrients for at least one generation (in one version, with shaking at 37°C (*28*)) before adding phages (*27*). Here, we resuspended exponentially growing cells in cold SM buffer and immediately added phages to begin adsorption. Thus, our cells were not starved before infection, and the observed lower frequency of lysogeny following infection in SM (analyzed as described below) does not contradict previous studies.

We also note that in (*27*), after starved cells had been incubated with phages in a 10 mM $MgSO_4$ solution for 30 minutes at 4°C, the infection mixture was diluted into tryptone-maltose broth at 32°C to trigger phage ejection. Our survey of different entry media (**Figure S23**) showed that in solutions containing tryptone and/or maltose, the average number of intracellular phage genomes does not saturate at one (as in SM, used in this study). Thus, the high frequency of lysogeny found in (*27*) was conceivably due to the combined effect of starvation and a medium permissive of multiple phage entries.

##### *16.2. Data analysis*

###### 16.2.1. Calculating the frequency of lysogeny

Under kanamycin selection, only lysogenic cells, which harbor prophages with the resistance cassette, can grow (*1*, *15*, *23*). As a result, the OD curves of infected cultures can be used to infer the number of lysogens. The frequency of lysogeny, $f_{\text{lysogeny}}$, is defined to be the fraction of lysogenic cells $(L_0)$ among total cells $(T_0)$, calculated as follows:

$$f_{\text{lysogeny}} = \frac{L_0}{T_0} = 2^{-\frac{1}{g}(t_L^* - t_T^*)} = 2^{-\frac{\Delta t^*}{g}}. \tag{16.1}$$

Here, $g$ is the doubling time of the cell cultures (approx. 35 min). $t_L^*$ and $t_T^*$ are the times the infected culture under selection and the uninfected culture without selection, respectively, reached a threshold OD (= 0.1) during the exponential growth phase. Hence, $\Delta t^*/g$ is the difference in the number of elapsed generations between the two samples.

Following this calculation, the maximum frequencies of lysogeny observed under the two conditions are $f_{\text{max,LBM}} = 0.009 \pm 0.001$ and $f_{\text{max,SM}} = 0.0003 \pm 0.0002$. Our model in **Section 17** aimed to use the medium-specific entry dynamics to account for this difference ($f_{\text{max,LBM}}/f_{\text{max,SM}} \approx 30.5$).

###### 16.2.2. Model fitting

We fitted **Equation 17.7** (derived in **Section 17** below) to the frequency of lysogeny as a function of the phage-to-bacteria ratio, $f_{\text{lysogeny}}(M)$, obtained following infection in LBM and SM. Only $a$ and $q_{\text{max}}$ were fitted parameters. Because the phage strain used in this assay was replication-deficient, following (*1*, *27*), we set $\text{MOI}^*$ to be 3. Parametrization of $\mu_n = \langle \lambda(n) \rangle$ was also pre-determined using microscopy data (see **Figure S23** for the parameters following infection in LBMM and SM).

While **Equation 17.7** includes a summation from 0 to infinity for $n$, for numerical purposes, the sum was taken from 0 to either 4-fold of $M$ or to 10, whichever is larger. Beyond this range, the probability $P(n > 4M)$ or $P(n > 10)$ becomes vanishingly small and was ignored. We confirmed that increasing the upper limit of this sum added computational time but resulted in no change in the fitted values.

Fitting was implemented with the MATLAB `lsqcurvefit` function, using nonlinear least-squares optimization and the trust-region algorithm. Because the sample size was small (6 data points for each series in LBM or SM), we bootstrapped each series by fitting to all 6 data points, then fitting to two subsets of the data (the 1st, 3rd, 5th points, and the 2nd, 4th, and 6th points). Estimates and SE of the parameters were calculated from the mean and SD of these three fitting runs.

This fitting procedure yielded the parameters $a$ and $q_{\text{max}}$ specific for each medium. In LBM, $q_{\text{max,LBM}} = 0.010 \pm 0.001$, and in SM, $q_{\text{max,SM}} = 0.003 \pm 0.001$. We interpreted these fitting results as follows (**Figure 4B**). The experimentally observed $f_{\text{max,LBM}}/f_{\text{max,SM}} \approx 30.5$ was captured using an inferred ratio $q_{\text{max,LBM}}/q_{\text{max,SM}}$ of only 3.2. This suggests that the remaining 9.5-fold could be attributed to the other difference between the two media: the specific parametrization of the distribution of intracellular phage numbers, $P(\lambda|n)$, as described in **Section 17**.

Differences in the fitted parameter $a$ ($a_{\mathrm{LBM}} = 0.51 \pm 0.03$, $a_{\mathrm{SM}} = 0.11 \pm 0.03$) only shifted the lysogeny-vs.-MOI curves horizontally when plotted in log-log scale and did not contribute to the maximum frequency of lysogeny. Data and the fits in **Figure 4B** are shown with the phage-to-bacteria ratios rescaled, $f_{\mathrm{lysogeny}}(a \cdot M)$, as done in (*1*).

### 17. Modeling bulk lysogenization following phage entry in different media

##### *17.1. Description of the model*

Following (*1*, *15*, *27*), our model mapped the phage-to-bacteria ratio in the infection mixture to a single-cell MOI, and the MOI to a probability at which the infected cell is lysogenized. However, while previous models omitted the entry stage, thus implicitly assuming all adsorbed phages enter the cell instantaneously, ours includes an explicit term for the entry dynamics. The model is depicted in **Figure 4C**.

The frequency of lysogeny as a function of the phage-to-bacteria ratio, $f_{\mathrm{lysogeny}}(M)$, is described using the following function:

$$f_{\mathrm{lysogeny}}(M) = \sum_{n=0}^{\infty} \left[ P(n|M) \sum_{\lambda=0}^{n} P(\lambda|n) \cdot Q(\lambda) \right]. \qquad (17.1)$$

**Equation 17.1** consists of three terms. The first term, $P(n|M)$, is the probability a cell is adsorbed by $n$ phages, given a phage-to-bacteria ratio $M$. Following (*1*, *27*), we assumed phage-bacteria collisions follow Poisson statistics. Hence, the number of adsorbed phages per cell follows the distribution below:

$$P(n|M) = \frac{(aM)^n e^{-aM}}{n!}, \qquad (17.2)$$

in which a scaling factor $a$ accounts for both the adsorption efficiency and the accuracy in measuring phage and cell concentrations.

In **Equation 17.1**, $P(\lambda|n)$ is the probability a cell has $\lambda$ intracellular phage genomes, given $n$ adsorbed phages. This distribution is described below (**Section 17.2**).

Finally, $Q(\lambda)$ is the probability a cell with $\lambda$ intracellular phage genomes is lysogenized. Following (*1*, *27*), we assumed that coinfection by $\lambda \geq \mathrm{MOI}^*$ phages is required for lysogeny. Accordingly, $Q(\lambda)$ is described using the following step-function:

$$Q(\lambda) = \begin{cases} 0 & \text{for } \lambda < \mathrm{MOI}^* \\ q_{\max} & \text{for } \lambda \geq \mathrm{MOI}^*, \end{cases} \qquad (17.3)$$

in which $q_{\max}$ is the maximum probability of lysogenization.

Using this parametrization of $Q(\lambda)$, **Equation 17.1** can be rewritten as follows**:**

$$f_{\mathrm{lysogeny}}(M) = q_{\max} \sum_{n=0}^{\infty} \left[ P(n|M) \sum_{\lambda=\mathrm{MOI}^*}^{n} P(\lambda|n) \right]. \qquad (17.4)$$

##### *17.2. Parametrization of the distribution of intracellular phage numbers*

If we assume that all adsorbed phages enter the cell instantaneously, $P(\lambda|n)$ can be parametrized as:

$$P(\lambda|n) = \begin{cases} 0 & \text{for } \lambda \neq n \\ 1 & \text{for } \lambda = n. \end{cases} \qquad (17.5)$$

As a result, **Equation 17.4** is rewritten as follows.

$$f_{\text{lysogeny}}(M) = q_{\max} \sum_{n=\text{MOI}^*}^{\infty} P(n|M) = q_{\max} \left( 1 - \sum_{n=0}^{\text{MOI}^*-1} \frac{(aM)^n e^{-aM}}{n!} \right). \tag{17.6}$$

We note that **Equation 17.6** has been reduced to the expression used to fit the bulk lysogenization data in (*1*), and is applicable when instantaneous entry is assumed.

To incorporate stochastic phage entry into the model, we parametrized $P(\lambda|n)$ using **Equation 5.3**: For cells adsorbed by $n$ phages, the intracellular phage number $\lambda$ is assumed to follow a truncated Poisson distribution controlled by a parameter $\mu_n$. With this parametrization of $P(\lambda|n)$, **Equation 17.4** is rewritten as follows:

$$f_{\text{lysogeny}}(M) = q_{\max} \sum_{n=0}^{\infty} \left[ \frac{(aM)^n e^{-aM}}{n!} \left( \sum_{j=0}^{n} \frac{\mu_n^j e^{-\mu_n}}{j!} \right)^{-1} \sum_{\lambda=\text{MOI}^*}^{n} \frac{\mu_n^\lambda e^{-\mu_n}}{\lambda!} \right]. \tag{17.7}$$

For simplicity, we set $\mu_n$, the mean of the Poisson distribution before truncation, to be equal to $\langle \lambda(n) \rangle$, the average number of intracellular phages in cells adsorbed by $n$ phages. $\langle \lambda(n) \rangle$ is parametrized by **Equation 5.1**, using experimental data obtained in each infection media (**Figure S23**). Therefore, $P(\lambda|n)$ is also specific to each medium (e.g., **Figure 4C**).

The values of $f_{\text{lysogeny}}(M)$ predicted using **Equations 17.6** and **17.7**, the latter parametrized using $\mu_n = \langle \lambda(n) \rangle$ in LBMM and SM, are shown in **Figure S25** ($\text{MOI}^*$, $a$, and $q_{\max}$ were kept constant). Given the same $q_{\max}$, the reduced probability of $P(\lambda \geq \text{MOI}^*)$ in SM results in a lower predicted $f_{\text{lysogeny}}(M)$ as compared to that in LBM.

#### 18. Stochastic model of phage entry kinetics

##### *18.1. Setup of the model*

The model is depicted in **Figure 2B**, and the variables used in the following derivations are listed in **Table S5**. For cells adsorbed by $n$ phages, the model outputs, at time $t$, the probability distribution of the number of intracellular phage genomes, $P(\lambda|n, t)$. In addition, the model also predicts the probability distribution of the time of the $i$-th phage entry, $P(T_i|n)$.

Our model is governed by three parameters: The entry probability of each adsorbed phage at infinite time ($\eta$); the rate (or probability per unit time) of entry initiation by each phage ($k$); and the time between entry initiation and detection ($\tau$). We note that to capture the experimental data, these three parameters were further parametrized as functions of $n$ (**Section 8.3.3**).

In the following derivations, the basic rules of probability were applied as described in (*30*, *31*). Symbolic integration was performed using Wolfram Mathematica, and the analytical solutions were confirmed using the stochastic simulation (**Section 19**), shown in **Figures S30–34**.

##### *18.2. Model assumptions*

For a cell adsorbed by $n$ phages, we assumed that only $m \leq n$ phages are capable of entering the cell. Biologically, a phage may not be entry-capable because of faulty capsid assembly or DNA packaging, improper docking to the cell's receptors, ionic conditions not conducive for DNA ejection, or other reasons (*32*–*37*). The probability per phage of being entry-capable is designated $\eta$. Thus, with $n$ adsorbed phages on the cell, $m$ follows a binomial distribution with a mean of $\eta n$.

We assumed that each of the entry-capable phages has a probability of initiating entry per unit of time (i.e., a rate) equal to $k$. For a cell with $m$ entry-capable phages, the probability per unit of time that an entry event is initiated by any of the phages is $mk$. The waiting time until the first entry initiation, $t_{0,1}$, thus follows an exponential distribution with a rate $mk$.

Following entry initiation, phage DNA is assumed to take a time $\tau$ to translocate into the cell and become labeled by the ParB-*parS* system. The time of the first phage entry, as measured, is thus $T_1 = t_{0,1} + \tau$.

After one phage has initiated entry, the number of remaining entry-capable phages is $m - 1$. The waiting time between the first and the second entry initiations, $t_{1,2}$, thus follows an exponential distribution with a rate $(m-1)k$. The time of the second phage entry is thus $T_2 = t_{0,1} + t_{1,2} + \tau$.

The same assumptions apply to the subsequent phages until all $m$ entry-capable phages have entered the cell. Once all entries have occurred, the number of intracellular phage genomes is equal to the initial number of entry-capable phages, $\lim_{t\to\infty} \lambda(t) = m$.

##### *18.3. Time-dependent average number of intracellular phage genomes*

Following the assumptions above, an expression for the average number of intracellular phage genomes, $\langle \lambda(n,t) \rangle$, can be derived as follows. The rate of change in the average number of entry-capable phages that have not yet ejected their genomes ("unejected"), $\langle U(t) \rangle$, is given by:

$$\frac{d\langle U \rangle}{dt} = -k \cdot \langle U(t) \rangle\,. \tag{18.1}$$

The solution of this ordinary differential equation, using the initial condition of $\langle U(t=0) \rangle = \langle m \rangle = \eta n$, is:

$$\langle U(n,t) \rangle = n\eta e^{-kt}\,. \tag{18.2}$$

Accounting for the delay between entry initiation and detection ($\tau$), the time-dependent average number of detected intracellular phage genomes in cells adsorbed by $n$ phages is:

$$\langle \lambda(n,t) \rangle = \langle m \rangle - \langle U(t) \rangle = n\eta\left(1 - e^{-k(t-\tau)}\right) \text{ for } t \geq \tau, \text{and 0 otherwise.} \tag{18.3}$$

**Equation 18.3** was used to predict the average number of intracellular phage genomes in **Figures 2E, 2F, S10, S13**, and **S14**.

##### *18.4. The probability distribution of phage entry time, given $m$*

We first aimed to derive the distribution of the time of the first phage entry, $T_1(m)$. The probability density function (PDF) of the waiting time until the first entry initiation is:

$$f\left(t_{0,1} = t|m\right) = f_{t_{0,1}|m}(t) = mke^{-mkt} \text{ for } t \geq 0 \text{ and } m \geq 1, \text{and 0 otherwise.} \tag{18.4}$$

The PDF in **Equation 18.4** is normalized for $t \in [0, \infty)$. The time of the first phage entry is $T_1 = t_{0,1} + \tau$. Because $\tau$ is a constant, the PDF of $T_1$ can be written by modifying **Equation 18.4**:

$$f(T_1 = t|m) = f_{T_1|m}(t) = mke^{-mk(t-\tau)} \text{ for } t \geq \tau \text{ and } m \geq 1, \text{and 0 otherwise.} \tag{18.5}$$

The PDF in **Equation 18.5** is normalized for $t \in [\tau, \infty)$. The cumulative distribution function (CDF) of $T_1$ is:

$$\begin{aligned} F(T_1 \leq t|m) = F_{T_1|m}(t) &= \int_{\tau}^{t} f_{T_1|m}(t')\,dt' = \int_{\tau}^{t} mke^{-mk(t'-\tau)}\,dt' \\ &= 1 - e^{-mk(t-\tau)} = m\left(\frac{1 - e^{-mk(t-\tau)}}{m}\right) \text{ for } t \geq \tau \text{ and } m \geq 1, \text{and 0 otherwise.} \end{aligned} \tag{18.6}$$

The last expression in **Equation 18.6** was rewritten in a way that enabled generalization to **Equation 18.20** below.

The average time of the first phage entry is the first moment of $f_{T_1|m}(t)$ (**Equation 18.5**):

$$\langle T_1(m)\rangle = \int_\tau^\infty t\cdot f_{T_1|m}(t)\,dt = \int_\tau^\infty t\cdot mke^{-mk(t-\tau)}\,dt = \frac{1}{mk} + \tau \text{ for } m \geq 1. \tag{18.7}$$

We note that after integration and rearrangement, **Equation 18.7** is simply the sum of the mean of the exponential distribution in **Equation 18.4**, $\langle t_{0,1}\rangle = {}^1/_{mk}$ and $\tau$. This matches our expectation for $T_1 = t_{0,1} + \tau$.

Next, we aimed to derive the distribution of the time of the second phage entry, $T_2(m)$. The PDF of the waiting time between the first and the second entry initiations is:

$$f(t_{1,2} = t|m) = f_{t_{1,2}|m}(t) = (m-1)ke^{-(m-1)kt} \text{ for } t \geq 0 \text{ and } m \geq 2, \text{ and 0 otherwise.} \tag{18.8}$$

$t_{0,1}$ and $t_{1,2}$ are two independent random variables, each describing a separate waiting time. Because the PDF of the sum of two independent random variables is the convolution of the two individual PDFs (given by **Equations 18.4** and **18.8**), the PDF of $t_{0,1} + t_{1,2}$ is:

$$\begin{aligned} f(t_{0,1} + t_{1,2} = t|m) &= (f_{t_{0,1}|m} * f_{t_{1,2}|m})(t) = \int_0^t f_{t_{0,1}|m}(t')\, f_{t_{1,2}|m}(t-t')\,dt' \\ &= \int_0^t mke^{-mkt'}(m-1)ke^{-(m-1)k(t-t')}\,dt' \\ &= m(m-1)ke^{-(m-1)kt}(1-e^{-kt}) \text{ for } t \geq 0 \text{ and } m \geq 2, \text{ and 0 otherwise.} \end{aligned} \tag{18.9}$$

Adding $\tau$ to $t_{0,1} + t_{1,2}$, the PDF of $T_2 = t_{0,1} + t_{1,2} + \tau$ is:

$$\begin{aligned} f(T_2 = t|m) = f_{T_2|m}(t) &= m(m-1)ke^{-(m-1)k(t-\tau)}(1-e^{-k(t-\tau)}) \\ &\text{for } t \geq \tau \text{ and } m \geq 2, \text{ and 0 otherwise.} \end{aligned} \tag{18.10}$$

Using the PDF in **Equation 18.10**, we can write the CDF of $T_2$:

$$\begin{aligned} F(T_2 \leq t|m) = F_{T_2|m}(t) &= \int_\tau^t f_{T_2|m}(t')\,dt' = \int_\tau^t m(m-1)ke^{-(m-1)k(t'-\tau)}(1-e^{-k(t'-\tau)})\,dt' \\ &= m(m-1)\left(-\frac{1-e^{-mk(t-\tau)}}{m} + \frac{1-e^{-(m-1)k(t-\tau)}}{m-1}\right) \text{ for } t \geq \tau \text{ and } m \geq 2, \text{ and 0 otherwise.} \end{aligned} \tag{18.11}$$

While the average time of the second phage entry could be found using the first moment of $f_{T_2|m}(t)$ (**Equation 18.10**), we note that $\langle T_2\rangle = \langle t_{0,1}\rangle + \langle t_{1,2}\rangle + \tau$. With $\langle t_{0,1}\rangle = {}^1/_{mk}$ (from **Equation 18.4**) and $\langle t_{1,2}\rangle = {}^1/_{(m-1)k}$ (from **Equation 18.8**), $\langle T_2(m)\rangle$ is equal to the following:

$$\langle T_2(m)\rangle = \frac{1}{mk} + \frac{1}{(m-1)k} + \tau \text{ for } m \geq 2. \tag{18.12}$$

To derive $T_3(m)$, we followed a similar approach. We first expressed the PDF of the waiting time between the second and the third entry initiations, $f_{t_{2,3}|m}(t)$, and used convolution to find the PDF of the time of the third phage entry, $T_3 = t_{0,1} + t_{1,2} + t_{2,3} + \tau$. The PDF of $T_3$ is as follows:

$$\begin{aligned} f(T_3 = t|m) = f_{T_3|m}(t) &= \frac{1}{2}m(m-1)(m-2)ke^{-(m-2)k(t-\tau)}(1-e^{-k(t-\tau)})^2 \\ &\text{for } t \geq \tau \text{ and } m \geq 3, \text{ and 0 otherwise,} \end{aligned} \tag{18.13}$$

and the corresponding CDF:

$$\begin{aligned}F(T_3 \leq t|m) &= F_{T_3|m}(t)\\ &= \frac{1}{2}m(m-1)(m-2)\left[\frac{1-e^{-mk(t-\tau)}}{m} - 2\left(\frac{1-e^{-(m-1)k(t-\tau)}}{m-1}\right) + \frac{1-e^{-(m-2)k(t-\tau)}}{m-2}\right]\\ &\text{for } t \geq \tau \text{ and } m \geq 3, \text{and 0 otherwise,}\end{aligned} \quad (18.14)$$

as well as the average entry time:

$$\langle T_3(m)\rangle = \frac{1}{mk} + \frac{1}{(m-1)k} + \frac{1}{(m-2)k} + \tau \text{ for } m \geq 3. \quad (18.15)$$

Using the same approach, we found the following PDF, CDF, and average value for $T_4(m)$:

$$\begin{aligned}f(T_4 = t|m) = f_{T_4|m}(t) = \frac{1}{6}m(m-1)(m-2)(m-3)ke^{-(m-3)k(t-\tau)}\left(1-e^{-k(t-\tau)}\right)^3\\ \text{for } t \geq \tau \text{ and } m \geq 4, \text{and 0 otherwise,}\end{aligned} \quad (18.16)$$

$$\begin{aligned}F(T_4 \leq t|m) &= F_{T_4|m}(t)\\ &= \frac{1}{6}m(m-1)(m-2)(m-3)\left[-\frac{1-e^{-mk(t-\tau)}}{m} + 3\left(\frac{1-e^{-(m-1)k(t-\tau)}}{m-1}\right)\right.\\ &\left. - 3\left(\frac{1-e^{-(m-2)k(t-\tau)}}{m-2}\right) + \frac{1-e^{-(m-3)k(t-\tau)}}{m-3}\right]\\ &\text{for } t \geq \tau \text{ and } m \geq 4, \text{and 0 otherwise,}\end{aligned} \quad (18.17)$$

$$\langle T_4(m)\rangle = \frac{1}{mk} + \frac{1}{(m-1)k} + \frac{1}{(m-2)k} + \frac{1}{(m-3)k} + \tau \text{ for } m \geq 4. \quad (18.18)$$

Taken together, **Equations 18.5**, **18.10**, **18.13**, and **18.16** can be generalized to the following form for the PDF of the time of the $i$-th phage entry, given $m$ entry-capable phages:

$$\begin{aligned}f(T_i = t|m) = f_{T_i|m}(t) &= \frac{1}{(i-1)!}\left[\prod_{j=1}^{i} m-(j-1)\right] ke^{-(m-(i-1))k(t-\tau)}\left(1-e^{-k(t-\tau)}\right)^{i-1}\\ &= \binom{m}{i} ike^{-(m-(i-1))k(t-\tau)}\left(1-e^{-k(t-\tau)}\right)^{i-1} \text{ for } t \geq \tau \text{ and } m \geq i, \text{and 0 otherwise.}\end{aligned} \quad (18.19)$$

**Equations 18.6**, **18.11**, **18.14**, and **18.17** can be generalized to the CDF of the time of the $i$-th phage entry, given $m$ entry-capable phages:

$$\begin{aligned}F(T_i \leq t|m) = F_{T_i|m}(t) &= \frac{1}{(i-1)!}\left[\prod_{j=1}^{i} m-(j-1)\right](-1)^i \sum_{j=1}^{i}(-1)^j \binom{i-1}{j-1}\frac{1-e^{-(m-(j-1))k(t-\tau)}}{m-(j-1)}\\ &= \binom{m}{i} i(-1)^i \sum_{j=1}^{i}(-1)^j \binom{i-1}{j-1}\frac{1-e^{-(m-(j-1))k(t-\tau)}}{m-(j-1)} \text{ for } t \geq \tau \text{ and } m \geq i, \text{and 0 otherwise.}\end{aligned} \quad (18.20)$$

**Equations 18.7**, **18.12**, **18.15**, and **18.18** can be generalized to the average time of the $i$-th phage entry, given $m$ entry-capable phages:

$$\langle T_i(m)\rangle = \tau + \sum_{j=1}^{i}\frac{1}{\left(m-(j-1)\right)k} \text{ for } m \geq i\,. \quad (18.21)$$

These generalizations were confirmed using the stochastic simulation (**Section 19**). For up to at least $m = 20$ (the experimentally relevant range), these analytical expressions (**Equations 18.19–18.21**) successfully captured the behavior of the simulation.

##### *18.5. The probability distribution of phage entry time, given $n$*

The probability mass function (PMF) of the number of entry-capable phages is:

$$P(m|n) = \binom{n}{m} \eta^m (1-\eta)^{n-m} . \tag{18.22}$$

The joint PDF for a cell to have $m$ entry-capable phages and the $i$-th entry time to occur at $T_i = t$, given $n$ adsorbed phages, is the product of $f_{T_i|m}(t)$ (**Equation 18.19**) and $P(m|n)$ (**Equation 18.22**):

$$\begin{aligned} f(T_i = t, m|n) &= f_{T_i,m|n}(t) = f_{T_i|m}(t)\, P(m|n) \\ &= C^{-1} \binom{n}{m} \eta^m (1-\eta)^{n-m} \binom{m}{i} ike^{-(m-(i-1))k(t-\tau)} \left(1 - e^{-k(t-\tau)}\right)^{i-1} \\ &\text{for } t \geq \tau, m \geq i, \text{and } n \geq i, \mathit{and}\ 0 \text{ otherwise,} \end{aligned} \tag{18.23}$$

in which $C$ is a normalization constant:

$$C = \sum_{j=i}^{n} \binom{n}{j} \eta^j (1-\eta)^{n-j} . \tag{18.24}$$

This re-normalization was necessary because the supports of $P(m|n)$, defined for $m \in [0, n]$, and of $f_{T_i|m}(t)$, defined for $m \geq i$, are different. As a result, the PMF of $m$ must be truncated.

The PDF of the time of the $i$-th phage entry, given $n$ adsorbed phages (regardless of how many of which are entry-capable), was found by summing the joint PDF $f_{T_i,m|n}(t)$ in **Equation 18.23** over all supported values of $m \in [i, n]$:

$$\begin{aligned} f(T_i = t|n) &= f_{T_i|n}(t) = \sum_m f_{T_i,m|n}(t) \\ &= C^{-1} \sum_{m=i}^{n} \binom{n}{m} \eta^m (1-\eta)^{n-m} \binom{m}{i} ike^{-(m-(i-1))k(t-\tau)} \left(1 - e^{-k(t-\tau)}\right)^{i-1} \\ &\text{for } t \geq \tau \text{ and } n \geq i, \text{and } 0 \text{ otherwise.} \end{aligned} \tag{18.25}$$

To calculate the average time of the $i$-th phage entry given $n$ adsorbed phages, we used **Equation 18.21** and **Equation 18.22**, as follows:

$$\langle T_i(n) \rangle = \sum_m \langle T_i(m) \rangle\, P(m|n) = \tau + C^{-1} \sum_{m=i}^{n} \left[ \binom{n}{m} \eta^m (1-\eta)^{n-m} \sum_{j=1}^{i} \frac{1}{\big(m-(j-1)\big)k} \right] \text{for } n \geq i \tag{18.26}$$

##### *18.6. The probability distribution of the intracellular phage number at time t, given $n$*

To derive $P(\lambda|n, t)$, we utilized the fact that, at time $t$, a cell would have exactly $\lambda = i$ intracellular phage genomes if the $i$-th phage entry has occurred by time $t$, but the $(i+1)$-th phage entry has not occurred yet. Therefore, given $m$ entry-capable phages, the probability that a cell has $\lambda = i$ intracellular phage genomes at time $t$ is:

$$P(\lambda = i|m, t) = F_{T_i|m}(t) - F_{T_{i+1}|m}(t) , \tag{18.27}$$

in which $F_{T_i|m}(t)$, given by **Equation 18.20**, describes the probability that the entry time of the $i$-th phage is less than or equal to $t$.

The joint probability that a cell with $m$ entry-capable phages has $\lambda = i$ intracellular phage genomes at time $t$, given $n$ adsorbed phages, is the product of $P(\lambda = i|m,t)$ (**Equation 18.27**) and $P(m|n)$ (**Equation 18.22**):

$$
\begin{aligned}
P(\lambda = i, m|n,t) &= P(\lambda = i|m,t)\, P(m|n) \\
&= \binom{n}{m} \eta^m (1-\eta)^{n-m} \left(F_{T_i|m}(t) - F_{T_{i+1}|m}(t)\right).
\end{aligned}
\tag{18.28}
$$

The probability that a cell has $\lambda = i$ intracellular phage genomes at time $t$, given $n$ adsorbed phages, was found by summing the joint probability in **Equation 18.28** over all values of $m \in [0,n]$:

$$
\begin{aligned}
P(\lambda = i|n,t) &= \sum_m P(\lambda = i, m|n,t) \\
&= \sum_{m=0}^{n} \binom{n}{m} \eta^m (1-\eta)^{n-m} \left(F_{T_i|m}(t) - F_{T_{i+1}|m}(t)\right) \\
&= \sum_{m=i}^{n} \binom{n}{m} \eta^m (1-\eta)^{n-m} \left(F_{T_i|m}(t) - F_{T_{i+1}|m}(t)\right).
\end{aligned}
\tag{18.29}
$$

The last simplification in **Equation 18.29** was possible because $F_{T_i|m}(t) = 0$ for $m < i$ (**Equation 18.20**). **Equation 18.29** was used to predict the theoretical distributions of intracellular phage numbers in **Figures 2G** and **S12**.

Using **Equation 18.29**, we arrived at the expression for the average number of intracellular phage genomes at time $t$, given $n$ adsorbed phages:

$$
\begin{aligned}
\langle \lambda(n,t) \rangle &= \sum_{i=0}^{n} i \cdot P(\lambda = i|n,t) \\
&= \sum_{i=0}^{n} i \cdot \left[ \sum_{m=i}^{n} \binom{n}{m} \eta^m (1-\eta)^{n-m} \left(F_{T_i|m}(t) - F_{T_{i+1}|m}(t)\right) \right].
\end{aligned}
\tag{18.30}
$$

Substituting $F_{T_i|m}(t)$ from **Equation 18.20** and integer values of $n \in [0,20]$ into **Equation 18.30**, we recovered the same expression for $\langle \lambda(n,t) \rangle$ obtained from the mean-field model in **Section 18.3** above, **Equation 18.3**.

##### *18.7. The case of cells with a single adsorbed phage*

For convenience, some simple expressions for the case of $n = 1$ are written below. From **Equation 18.25**, the PDF of the phage entry time in cells adsorbed by one phage (shown in **Figure 2D**) is:

$$
f(T_1 = t|n = 1) = k e^{-k(t-\tau)} \text{ for } t \geq \tau, \text{and 0 otherwise.} \tag{18.31}
$$

From **Equation 18.26**, the average time of phage entry in cells adsorbed by one phage is:

$$
\langle T_1(n = 1) \rangle = \tau + \frac{1}{k}. \tag{18.32}
$$

In both **Equations 18.31** and **18.32**, because the entry time is only defined in cells with phage entry, the parameter $\eta$ (controlling whether the adsorbed phage is entry-capable) does not appear in the expressions.

From **Equations 18.20** and **18.29**, the probability that a cell with one adsorbed phage has one intracellular phage genome at time $t$ (shown in **Figure S12**) is:

$$
P(\lambda = 1|n = 1,t) = \eta\left(1 - e^{-k(t-\tau)}\right) \text{ for } t \geq \tau, \text{and 0 otherwise.} \tag{18.33}
$$

Because $\lambda$ is either 0 or 1 for $n = 1$, from **Equation 18.33**, the time-dependent average number of intracellular phage genomes in cells with one adsorbed phage (shown in **Figure 2C**) is simply:

$$\langle \lambda(n = 1, t) \rangle = \eta\left(1 - e^{-k(t-\tau)}\right) \text{ for } t \geq \tau, \text{and 0 otherwise.} \tag{18.34}$$

### 19. Simulating the stochastic model

##### *19.1. Description of the simulation*

This stochastic simulation was based on the mathematical model described in **Section 18** and utilized the Gillespie algorithm (*38*). Below, we briefly summarize the pertinent parts of the model in the language of the simulation.

We assumed that an entry-capable phage exists in either one of the following two states: Adsorbed on the cell but unejected ("Outside"), or having ejected its DNA, now intracellular ("Inside"). There is only one forward reaction: A phage can transition from the "Outside" state to the "Inside" state using an "Ejection" reaction at a rate constant of $k$. For a cell with $m$ entry-capable phages and $\lambda(t)$ "Inside" phages at time $t$, the time-dependent number of "Outside" phages is $m - \lambda(t)$. Hence, the propensity for the "Ejection" reaction is:

$$a_{\text{ejection}} = (m - \lambda)k\,. \tag{19.1}$$

Because there is no reverse reaction, eventually, all entry-capable phages will be "Inside", i.e., $\lim_{t-\infty} \lambda(t) = m$. This serves as the terminal condition in the simulation.

##### *19.2. Simulation algorithm*

Prior to simulation, the MATLAB random number generator was seeded using `rng('shuffle')`.

For a cell adsorbed by $n$ phages, with $t = 0$ defined as the time of phage adsorption:

(1) Determine the parameters $(\eta, k, \tau)$ that govern the dynamics of phage entry in this cell. If these parameters are to be MOI-dependent, use the parametrization described in **Section 8.3.3**.
(2) Draw $m$, the number of entry-capable phages, randomly from a binomial distribution with parameters $n$ and $\eta$ (implemented using the MATLAB `binornd` function).
(3) Initialize the system: $t = 0$ and $\lambda(t = 0) = 0$.
(4) Simulate the waiting time between phage entries:
    (4.1) Calculate the ejection propensity (using **Equation 19.1**).
    (4.2) Draw a random number $r$ from a uniform distribution in the unit interval (implemented using the MATLAB `rand` function).
    (4.3) Calculate the waiting time until the next entry initiation:
$$t_{\text{waiting}} = \frac{1}{a_{\text{ejection}}} \log \frac{1}{1 - r}\,. \tag{19.2}$$
    (4.4) Record $t_{\text{waiting}}$ into a vector of waiting times between entry initiations, $\mathbf{t}_{\text{between}}$.
    (4.5) Update the system: $t \leftarrow t + t_{waiting}$, and $\lambda \leftarrow \lambda + 1$.
    (4.6) If $\lambda = m$, terminate the waiting-time simulation and proceed to step 5. Else, return to step 4.1.
(5) Determine the time from adsorption to the initiation of each phage entry, $\mathbf{t}_{\text{initiation}}$, equal to the cumulative sum of $\mathbf{t}_{\text{between}}$.
(6) Determine the time from adsorption to when the intracellular phage genome is detected, $\mathbf{t}_{\text{entry}} = \mathbf{t}_{\text{initiation}} + \tau$. This is equivalent to the phage entry times as experimentally measured.

##### *19.3. Processing of simulation results*

The simulation in **Section 19.2** was performed for $n \in [0{,}20]$, with 10000 cells for each $n$ value. For each simulated cell, the vector of simulated entry times was processed as a series of entry events following synchronized adsorptions (**Figure S26**). Comparisons between the stochastic simulation and the analytical expressions of the model (**Section 18**) are shown in **Figures S30–S34**.

#### 20. Stochastic simulation of infection outcome in individual cells

##### *20.1. Summary of the mathematical model in Yao et al.*

Our simulation for the infection outcome was based on the model in Yao et al. (*1*). Below, we briefly describe this model where pertinent to the current work.

Yao et al. modeled the cell-fate decision of phage lambda using a set of ordinary differential equations (ODEs) that track the mRNA and protein concentrations of three genes governing the decision: *cI*, *cII*, and *cro*, as well as the concentration of the phage genomes in the infected cell. The cell volume was assumed to grow exponentially over time.

The input of the model is the initial viral concentration, defined to be the MOI divided by the initial cell volume. The MOI is the initial intracellular number of phage genomes; the model in Yao et al. was agnostic to the number of adsorbed phages or the phage-to-bacteria ratio in the environment. The initial cell volume was set to be 1 $\mu m^3$, uniform for all cells.

Depending on whether infection of a replication-competent or replication-deficient phage is modeled, the copy number of the phage genomes may increase over time. The ODEs were solved numerically to yield the kinetics of Cro (driving lysis) and CI (driving lysogeny), whose concentrations were compared to respective thresholds to determine the cell fate. Four outcomes are possible: Failed infection, lysis, lysogeny, and mixed outcome.

Because the initial cell volume is assumed uniform, the predicted cell fate for a given MOI is deterministic, and the frequency of lysogeny is a step-function of MOI. In particular, for infection by replication-competent phages, cells with MOI = 1 are lytic, while cells with MOI ≥ 2 are lysogenic.

##### *20.2. Summary of the single-cell data in Zeng et al.*

We aimed to compare our simulation results with the single-cell data from Zeng et al. (*15*). Below, we briefly describe this data where pertinent to this study.

Zeng et al. measured the fate of individual cells following infection by a replication-competent phage strain. Capsids of the infecting phages were fluorescently labeled using a scheme similar to the current study, and the single-cell MOI was defined as the number of phages adsorbed to the cell (Zeng et al. did not measure the number of intracellular phage genomes). After 30 minutes of incubation at low temperature to allow phages to adsorb to the cell and 5 minutes at 35°C to trigger phage ejection, the infection mixtures in Zeng et al. were diluted using room-temperature (RT) medium, mounted, and imaged in a time-lapse at RT. The infected cells harbored fluorescence reporters that enabled the detection of cell fate. In Zeng et al., the measured frequency of lysogeny increased gradually with the number of adsorbed phages.

In addition, Zeng et al. found cell size to affect the infection outcome. For a given MOI, smaller cells had a higher chance of being lysogenized. The distribution of normalized cell size in Zeng et al. was approximated as a log-normal distribution ($\mu = 1, \sigma = 0.28$).

##### *20.3. Description of our simulation*

We performed the following simulation for cells adsorbed by $n \in [0,5]$ phages, with 1000 cells for each $n$ value.

For a cell adsorbed by $n$ phages:

(1) Determine the initial cell size (drawn from a log-normal distribution, described in **Section 20.4** below).
(2) Determine the single-cell MOI at 5 minutes (accounting for stochastic phage entries, described in **Section 20.5** below).

(3) Calculate the initial viral concentration.
(4) Apply the Yao et al. model to determine the cell fate.

The system of ODEs from Yao et al. was solved for infection by replication-competent phages as described in the original paper. The frequency of lysogeny was calculated as the fraction of lysogenic cells among lytic or lysogenic cells (i.e., failed infection and mixed outcome were not included). Following how Zeng et al. reported the data, the decision curve (**Figure 4E**) depicts the frequency of lysogeny as a function of the number of adsorbed phages, $f_{\text{lysogeny}}(n)$, regardless of how many intracellular phage genomes are in the cell.

##### *20.4. Incorporating variations in cell size*

We first aimed to introduce variations in cell size into the Yao et al. model. For each cell with a given number of adsorbed phages ($n$), we randomly generated the initial cell size by drawing from a log-normal distribution with $\mu = 1, \sigma = 0.28$ (as described in Zeng et al.), implemented using the MATLAB `lognrnd` function. This initial cell size was used to calculate the initial viral concentration, with the MOI equal to the number of adsorbed phages.

Variations in cell size introduced some variability to the cell fate. In particular, while Yao et al. deemed all cells with MOI = 1 to be Cro-dominant, thus lytic, variations in cell size rendered approx. 20% of the cells lysogenic. This fraction reflected infection in very small cells, such that the initial concentration of phage genomes was higher, facilitating increased CI concentration (**Figures 4D** and **S28**), thus lysogeny. Because infection in very large cells reduced the initial viral concentration, for MOI = 2, the frequency of lysogeny decreased from 100% in Yao et al. to approx. 98%. Variations in cell size did not change the frequency of lysogeny for MOI higher than 2. We used these predictions by the model with cell size variations as a baseline to determine the effect of phage entry dynamics on the infection outcome.

##### *20.5. Incorporating stochastic phage entry*

Next, we introduced stochastic phage entries into the model with cell size variation. For each cell with a given number of adsorbed phages ($n$), we implemented the simulation of stochastic phage entries (described in **Section 19**) to determine the time series of phage entries in this cell. On average, the simulated number of intracellular phage genomes was lower than the number of adsorbed phages, and the phage genomes entered the cell non-simultaneously. Because Zeng et al. diluted the infection mixture using RT medium and imaged the cells at RT, effectively halting additional phage entries after 5 minutes, we set the MOI that drives the lysis vs. lysogeny decision to be the number of phage genomes that have entered the cell within 5 minutes. The initial viral concentration is thus $\lambda(t = 5\ \text{min})$ divided by the initial cell size (drawn from a log-normal distribution as above).

Stochastic phage entries had a strong effect on the decision curve (**Figures 4D** and **4E**). For example, among cells adsorbed by $n = 2$ phages, approx. 50% of the cells had only one intracellular phage genome by 5 minutes (see **Figure S27** for the distributions of intracellular phage numbers). In such cells, the increased concentration of Cro (**Figures 4D** and **S28**) rendered the cell lytic. Similarly, the frequency of lysogeny also decreased for MOI higher than 2, and even in cells adsorbed by 5 phages, not all of the infected cells were lysogenic.

We note that within our simulation framework, stochastic phage entries had no effect on the infection outcome of MOI = 1. For this MOI, failed or delayed phage entry, which would result in failed infection (**Figure S29**), did not change the decision curve, which was calculated based on the ratio between the lytic and lysogenic cells only.

##### *20.6. Fitting to the Hill function*

The Hill equation (below) was fitted to the frequency of lysogeny as a function of the number of adsorbed phages, $f_{\text{lysogeny}}(n)$ for the data from Zeng et al. and our simulation results.

$$f_{\text{lysogeny}}(n) = \frac{n^h}{K^h + n^h}. \tag{20.1}$$

In **Equation 20.1**, $h$ is the Hill coefficient, which describes the degree of precision in the decision curve (*39*), and $K$ is the MOI at which the frequency of lysogeny reaches half its maximum. For Zeng et al., data for "all cells" in Figure 2C of the original paper was extracted using plotdigitizer.com, and bootstrapping was performed by resampling the mean values. For our simulation results, bootstrapping was performed on the single-cell level as described in **Section 5.6**.

Fitting results are shown in **Figure 4E**. The simulation with cell size variation alone gave $h = 7.87 \pm 0.41$, $K = 1.18 \pm 0.01$. The simulation with both cell size variation and stochastic phage entries gave $h = 1.84 \pm 0.08$, $K = 1.81 \pm 0.05$. Refitting Zeng et al. data gave $h = 0.94 \pm 0.20$, $K = 1.71 \pm 0.29$, similar to the published values, $h = 1.0 \pm 0.1$, $K = 1.8 \pm 0.1$.

#### SUPPLEMENTARY FIGURES

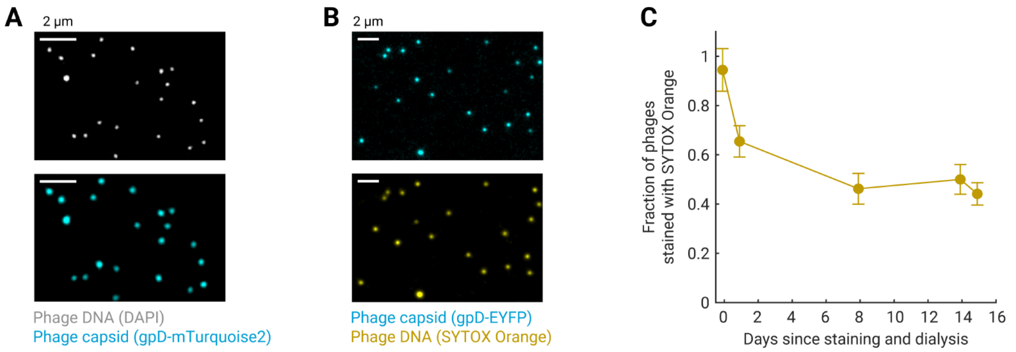


**Figure S1. Efficiency of phage labeling using fluorescent capsids and SYTOX Orange.**

**(A)** Labeling efficiency of fluorescent phage capsids. Phage capsids and encapsidated phage DNA were labeled using fluorescent gpD fusions and 4′,6-diamidino-2-phenylindole (DAPI), respectively, and the phage solution was imaged as described in **Methods, Section 3.1**. The efficiency of capsid labeling, defined to be the fraction of phage foci with both capsid and DAPI signals, was 97.0 ± 1.0% (mean and SE from 3 phage purification replicates; *N* = 152–442 phages counted per replicate).

**(B)** Labeling efficiency of SYTOX Orange. Encapsidated phage DNA was labeled using SYTOX Orange, and the phage solution was imaged as described in **Methods, Section 3.2**. The efficiency of SYTOX Orange labeling, defined to be the fraction of capsid foci with colocalized SYTOX Orange signal, was 91.9% ± 1.6% (mean and SE from 3 staining and dialysis replicates; *N* = 127–291 phages counted per replicate).

**(C)** The labeling efficiency of SYTOX Orange declined over time. The same phage solution, stained using SYTOX Orange, was imaged over 15 days, and the fraction of stained phages on each day was measured. Markers, mean ± SE (*N* = 119–213 phages counted on each day).

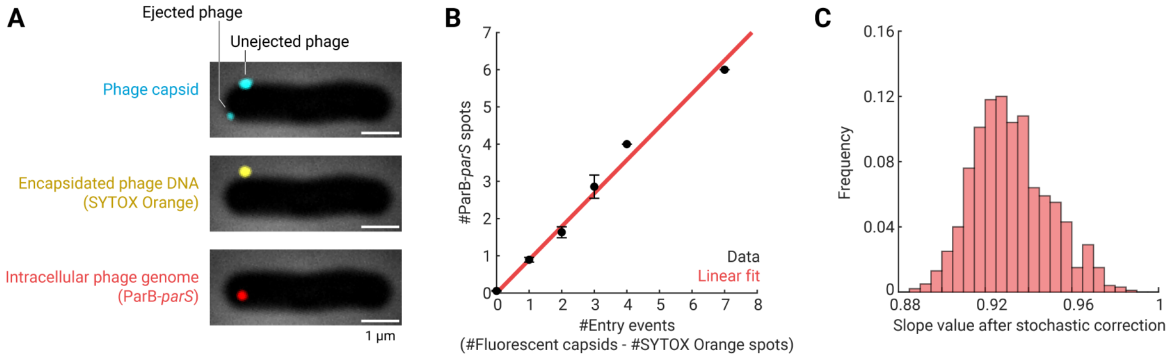


**Figure S2. Efficiency of genome detection using the ParB-*parS* system.**

**(A)** Co-labeling of phages using fluorescent capsids, SYTOX Orange (for encapsidated phage DNA), and ParB-*parS* (for intracellular phage genomes), as described in **Methods, Section 9**.

**(B)** The detection efficiency of the ParB-*parS* system is ~90%. Black markers, mean ± SE of the number of ParB-*parS* spots versus the number of fluorescent capsids without SYTOX Orange signals in the same cell ($N$ = 554 cells). Red line, linear fit with the $y$-intercept forced to zero, the slope of which reflects the detection efficiency of the ParB-*parS* system (= 0.89 ± 0.04, SE from bootstrapping).

**(C)** Correcting for the staining efficiency of SYTOX Orange. The number of phages stained with SYTOX Orange in each cell prior to infection was stochastically simulated as described in **Methods, Section 9.3**. Red bars, histogram of the detection efficiency of the ParB-*parS* system in each simulated dataset ($N$ = 1000 realizations, mean ± SD = 0.93 ± 0.02).

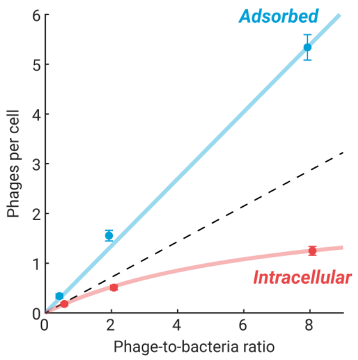


**Figure S3. Relation between the numbers of adsorbed and intracellular phages per cell and the phage-to-bacteria ratio.**

The average number of adsorbed phages and intracellular phage genomes were measured in infection mixtures with different phage-to-bacteria ratios, using the bulk infection assay as described in **Methods, Section 5**. Markers, mean ± SE ($N$ = 201–221 cells counted for each of the three samples at different phage-to-bacteria ratios). Cyan line, linear fit to the numbers of adsorbed phages per cell. Red curve, fit to a Michaelis-Menten function for the numbers of intracellular phage genomes, serving as a guide to the eye. Dashed line, linear scaling of the intracellular phage numbers, extrapolated from the sample with the phage-to-bacteria ratio of 0.5.

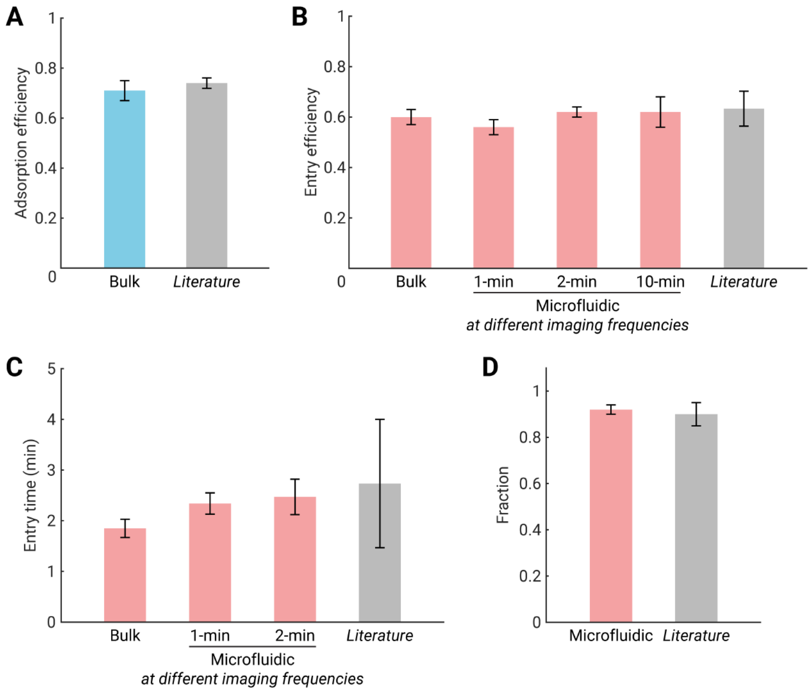


**Figure S4. Comparison with literature-reported values for the efficiency and rate of phage adsorption and entry.**

**(A)** Efficiency of phage adsorption. Cyan bar, mean ± SE of the adsorption efficiency in samples with phage-to-bacteria ratio of 0.5, 2, and 8, measured using the bulk infection assay (**Methods, Section 5**). Gray bar, mean ± SE of values reported in the literature, 0.70–0.75 (*11*), ~0.70 (*40*), and 0.77 ± 0.03 (*37*).

**(B)** Efficiency of phage entry. Red bars, mean ± SE of the entry efficiency of the first phage, measured in bulk infection (**Methods, Section 5**) and in the microfluidic device at different imaging frequencies (**Methods, Section 8**). The value at 1-minute frequency was compared to those at 2- and 10-minute frequencies to control for phototoxicity**.** Gray bar, mean ± SE of values reported in the literature, 0.51–0.64 (*41*), and 0.75 ± 0.01 (*37*).

**(C)** Average entry time of the first phage. Red bars, mean ± SE of when the first phage genome was detected in the cell, measured in bulk infection (**Methods, Section 7** and **Figure S8**) and in the microfluidic device at different imaging frequencies (**Methods, Section 8**). Gray bar, mean ± SE of values reported in the literature, 2–3 minutes (*41*), 1–2 minutes (*12*), and 5.2 ± 4.2 minutes (*7*).

**(D)** Fraction of phages having ejected their genomes by 5 minutes after adsorption. Red bar, mean ± SE for infection in the microfluidic device (**Methods, Section 8**); the corresponding distribution is shown in **Figure 2D**. Gray bar, mean ± SE of values reported in the literature, ~0.95 (*2*), and ~0.85 (*37*).

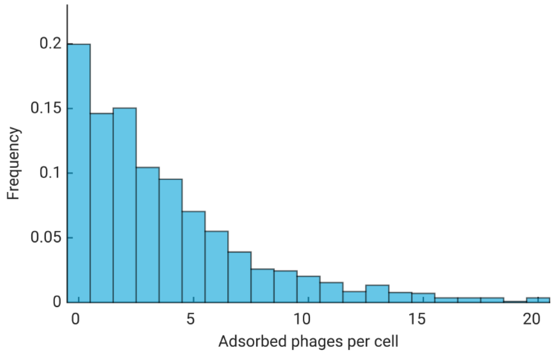


**Figure S5. Distribution of the number of phages adsorbed to the cell.**

Cyan bars, histogram of the numbers of adsorbed phages per cell following bulk infection in LBMM as described in **Methods, Section 5** (*N* = 1437 cells, pooled from 7 independent experiments).

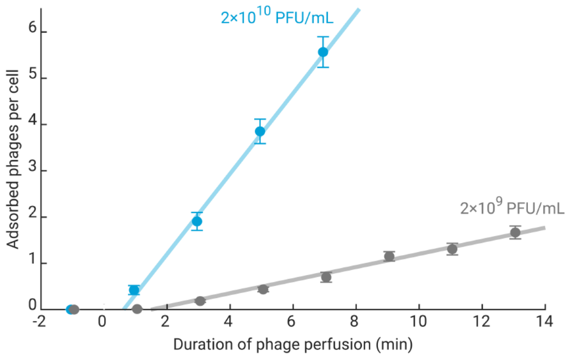


**Figure S6. The number of adsorbed phages as a function of perfusion time and phage concentration.**

A phage solution at either $2\times10^{10}$ PFU/mL (in cyan) or $2\times10^{9}$ PFU/mL (in gray) was perfused into the microfluidic device as described in **Methods, Section 8**, and the time-dependent numbers of adsorbed phages per cell were measured. Markers, mean ± SE (*N* = 57–247 cells for each data point). Cyan and gray lines, linear fits, the slope of which describes the rate of phage accumulation (approx. 1 and 0.1 phages/cell per minute, respectively). The phage concentration of $2\times10^{10}$ PFU/mL and the flow time of 2 minutes were used for most microfluidic experiments in this study.

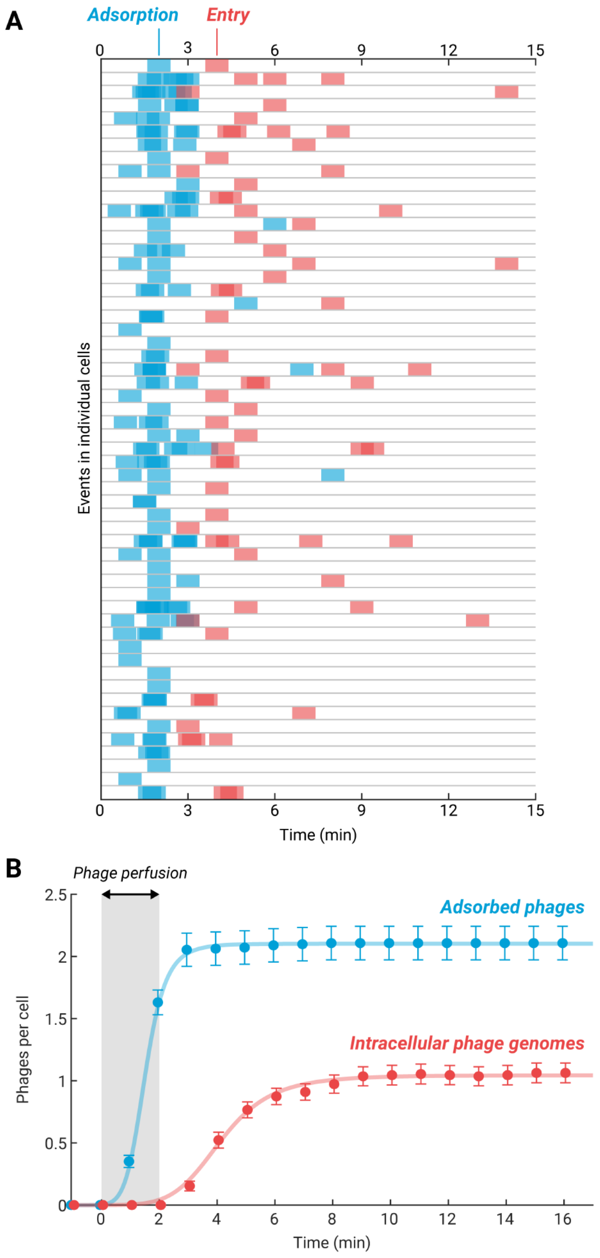


**Figure S7. Kinetics of phage adsorption and entry in the microfluidic infection assay.**

**(A)** The time series of phage adsorption and entry events from a representative microfluidic experiment, performed as described in **Methods, Section 8**. In this experiment, 222 cells were tracked, of which $N$ = 111 cells with at least one adsorbed phage are shown here. When multiple adsorption or entry events were observed within the same imaged time point, the cyan or red boxes, respectively, were slightly shifted for clarity. A subset of these time series is reproduced in **Figure 2A**.

**(B)** The time-dependent average numbers of adsorbed phages and intracellular phage genomes from the experiment shown in Panel A. Phages were perfused into the cell chamber between $t$ = 0 and 2 minutes (indicated by the gray shading), after which excess phages were washed out. Markers, mean ± SE. Cyan and red curves, fits to Hill equations, serving as a guide to the eye.

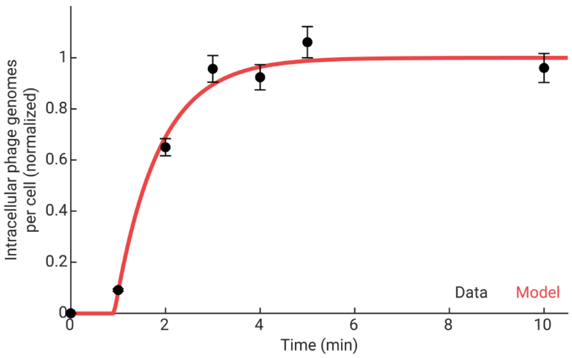


**Figure S8. Kinetics of the first phage entry following bulk infection.**

The time-dependent numbers of intracellular phage genomes were measured following bulk infection at a phage-to-bacteria ratio of 0.5 (**Methods, Section 7**). Black markers, mean ± SE ($N$ = 253–376 cells counted for each time point, 2019 cells in total). Red curve, fit to **Equation 7.1**. The inferred parameters are $k$ = 1.08 ± 0.19 per min, and $\tau$ = 0.92 ± 0.08 min (SE from bootstrapping). Both the data and the fit are shown after normalization, as described in **Methods, Section 7.3**.

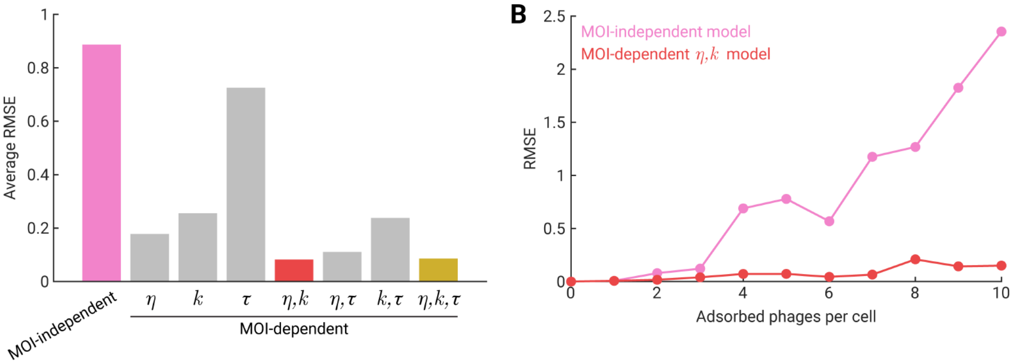


**Figure S9. The MOI-dependent $\eta, k$ model captures the time-lapse infection data most closely.**

**(A)** Average root-mean-squared-error (RMSE) of the 8 model variations described in **Methods, Section 8.3.2**. The RMSE was calculated using the time-dependent average number of intracellular phage genomes in cells adsorbed by $n$ = 0–10 phages, obtained using the microfluidic infection assay (**Methods, Section 8**). Pink bar, RMSE of the MOI-independent model in which the parameters $\eta, k, \tau$ in cells with MOI > 1 were the same as that in cells with MOI = 1. Red bar, RMSE of the model in which $\eta$ and $k$ were allowed to vary with the MOI, while $\tau$ remained the same as MOI = 1. Yellow bar, RMSE of the model in which all three parameters were allowed to vary with the MOI. Gray bars, the other 5 model variations in which one or two of the parameters were allowed to vary with the MOI.

**(B)** RMSE of the time-dependent average number of intracellular phage genomes as a function of the number of adsorbed phages. In pink, the RMSE of the MOI-independent model increases with the MOI. In red, the RMSE of the MOI-dependent model (i.e., $\eta$ and $k$ were allowed to vary with the MOI) remained low at high MOI.

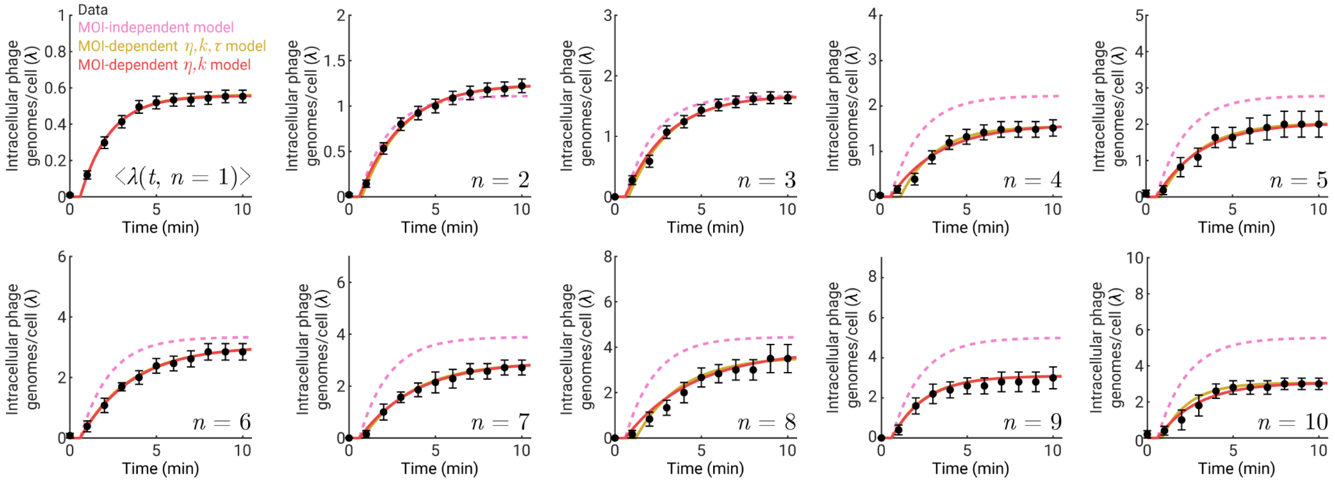


**Figure S10. The MOI-dependent model successfully fits to the time-dependent average number of intracellular phage genomes.**

Black markers, the time-dependent average number of intracellular phage genomes in cells adsorbed by $n$ = 1–10 phages (*N* = 208, 90, 58, 31, 11, 13, 7, 6, 5, and 5 cells for each MOI, respectively), obtained using the microfluidic infection assay (**Methods, Section 8**). Dashed curves in pink, predictions by a model in which all three parameters ($\eta$, $k$, and $\tau$) in cells with MOI > 1 were equal to those in cells with MOI = 1. Solid curves in yellow, fits by a model in which all three model parameters were allowed to vary with the MOI. Solid curves in red, fits by a model in which $\eta$ and $k$ were allowed to vary with the MOI. The yellow curves are largely indistinguishable from the red curves. Model predictions were calculated using **Equation 18.3**. Data and model predictions for MOI = 1, 2, 4, 6, 8, and 10 are reproduced in **Figures 2C** and **2E**.

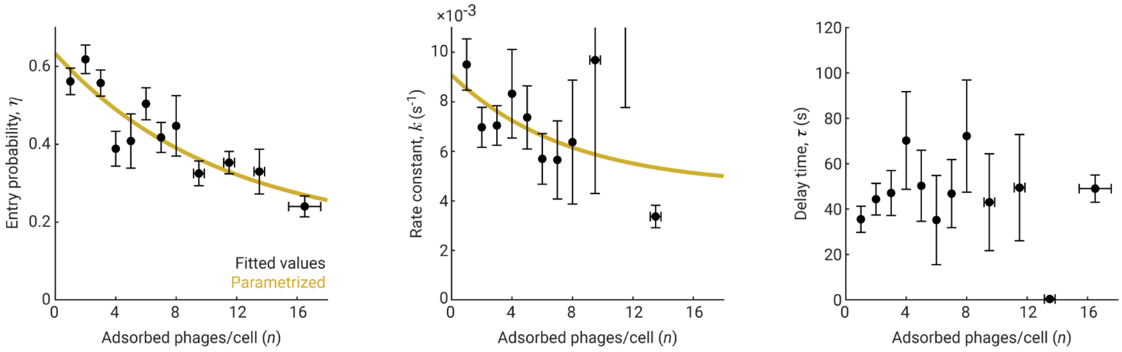


**Figure S11. Fitted parameter values in the MOI-dependent $\eta, k, \tau$ model.**

Black markers, fitted values ± SE (from bootstrapping) of the model parameters when all three parameters ($\eta$, $k$, and $\tau$) were allowed to vary with MOI ($n$); cells at higher MOI were binned together to allow for at least 10 cells per bin. Yellow curves, parametrization: $\eta$ and $k$ as exponentially decaying functions of MOI. The fitted values of $\tau$ showed no clear trend as a function of the MOI. See **Methods, Section 8** for details.

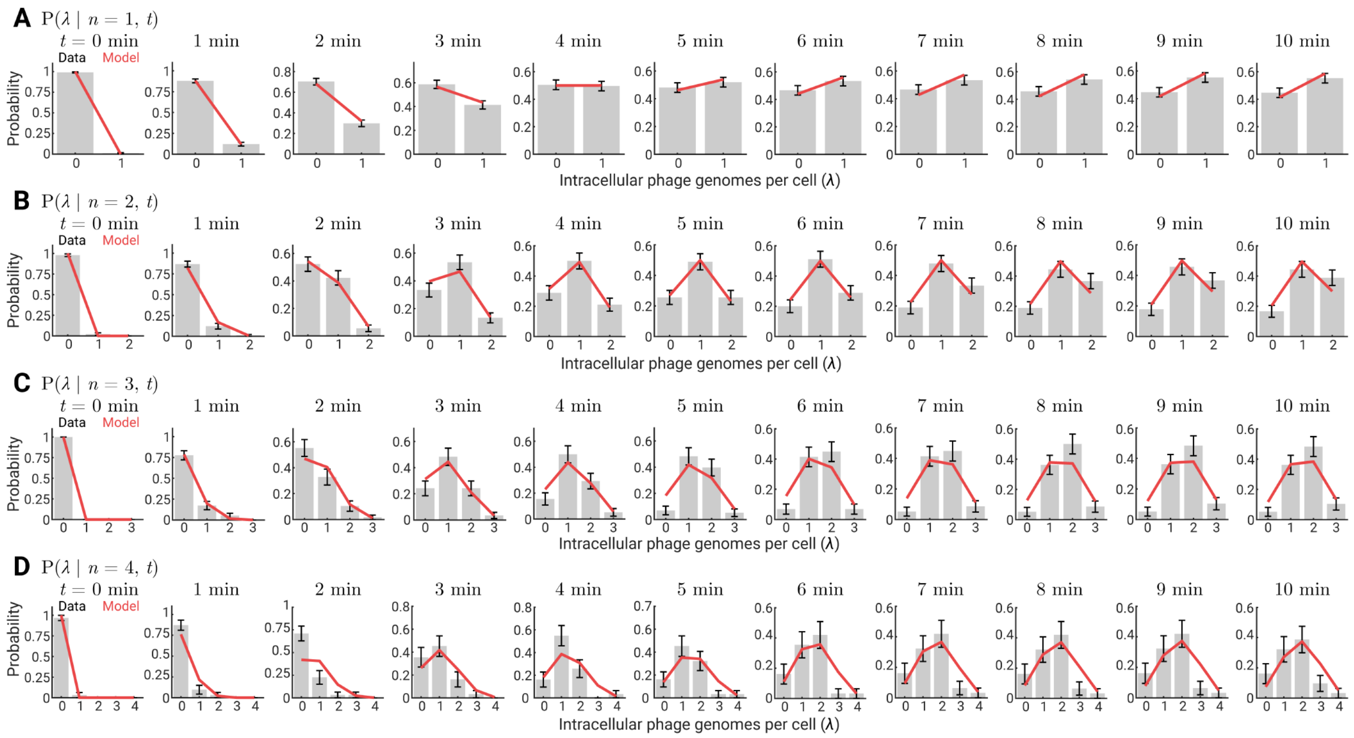


**Figure S12. The MOI-dependent model successfully predicts the distributions of intracellular phage numbers.**

Data and model predictions for the distributions of intracellular phage numbers at a given time ($t$ = 0–10 minutes) in cells adsorbed by (**A**–**E**) $n$ = 1–5 phages, respectively, obtained using the microfluidic infection assay (**Methods, Section 8**). Gray bars, histograms of the data, with error bars indicating SE ($N$ = 208, 90, 58, and 31 cells for each MOI, respectively). Red curves, model predictions, calculated using **Equation 18.29**. Data and model predictions for MOI = 4 at 1, 3, 5, and 10 minutes are reproduced in **Figure 2G**.

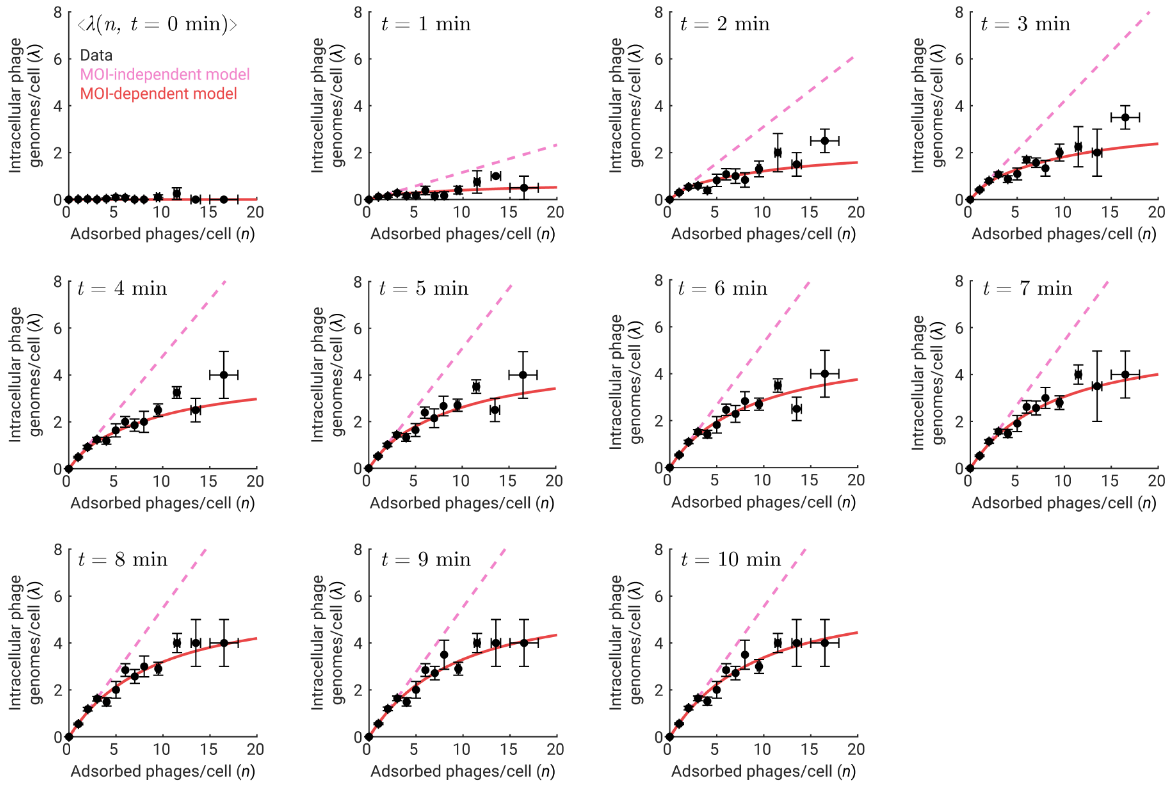


**Figure S13. The MOI-dependent model reproduces the sublinear relation between the numbers of intracellular and adsorbed phages following infection in the microfluidic device.**

Black markers, mean ± SE ($N$ = 1030 cells tracked at each time point) of the average number of intracellular phage genomes in cells with a given number of adsorbed phages, obtained using the microfluidic infection assay (**Methods, Section 8**). Cells at higher MOI were binned together to allow for at least 10 cells per bin. Dashed line in pink, prediction by the MOI-independent model. Red curve, prediction by the MOI-dependent model. Model predictions were calculated using **Equation 18.3**. Data and model predictions at 10 minutes are reproduced in **Figure 2H**.

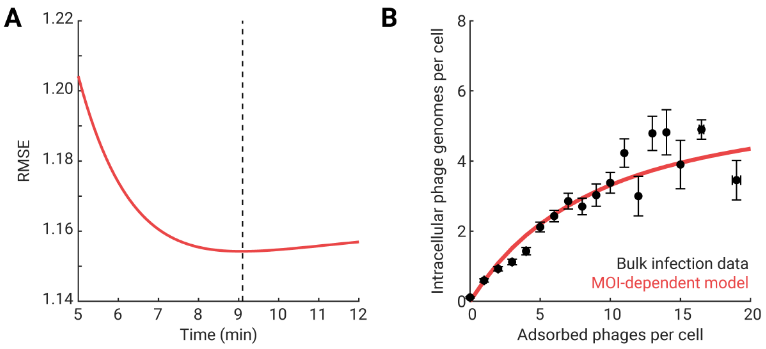


**Figure S14. The MOI-dependent model captures the sublinear relation between the numbers of intracellular and adsorbed phages following bulk infection.**

**(A)** Goodness-of-fit of the model prediction for the number of intracellular phage genomes as a function of the number of adsorbed phages, obtained using the bulk infection assay (**Methods, Section 5**). The time input to the model was scanned between 5 minutes and 12 minutes at a 0.1-minute increment (**Methods, Section 5.5**). Dashed line, 9.1 minutes, at which the root-mean-squared-error (RMSE) was minimized.

**(B)** Data and model predictions. Black markers, mean ± SE ($N$ = 1437 cells, pooled from 7 independent experiments); cells at higher MOI were binned together to allow for at least 10 cells per bin. Red curve, predictions by the MOI-dependent model at $t$ = 9.1 minutes, calculated using **Equation 18.3**.

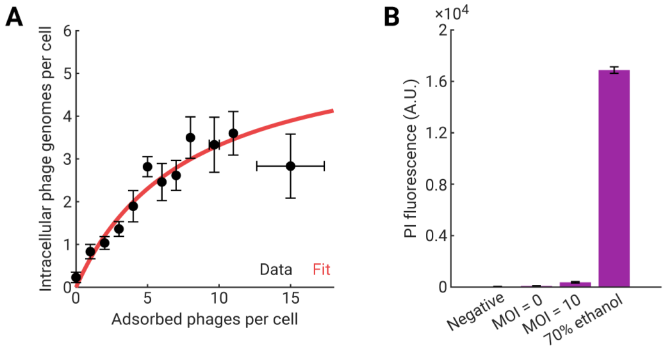


**Figure S15. Controls for experiments involving propidium iodide (PI).**

**(A)** The sublinear relation between the numbers of intracellular and adsorbed phages was reproduced when cells were stained with PI after phage infection (performed as described in **Methods, Section 6**). Black markers, mean ± SE (*N* = 188 cells; PI was added to an aliquot of the infection mixture after 5 minutes of 37°C incubation). Red curve, fit to a Michaelis-Menten function, serving as a guide to the eye.

**(B)** Comparing the intracellular PI fluorescence in the infected sample and the controls (error bars indicate SE). Data for the negative control (not mixed with phages, *N* = 39 cells), MOI = 0 and MOI = 10 (*N* = 22 and 30 cells, respectively), and the positive control (treated with 70% ethanol, *N* = 25 cells) are shown. See **Methods, Section 6** for details of each sample.

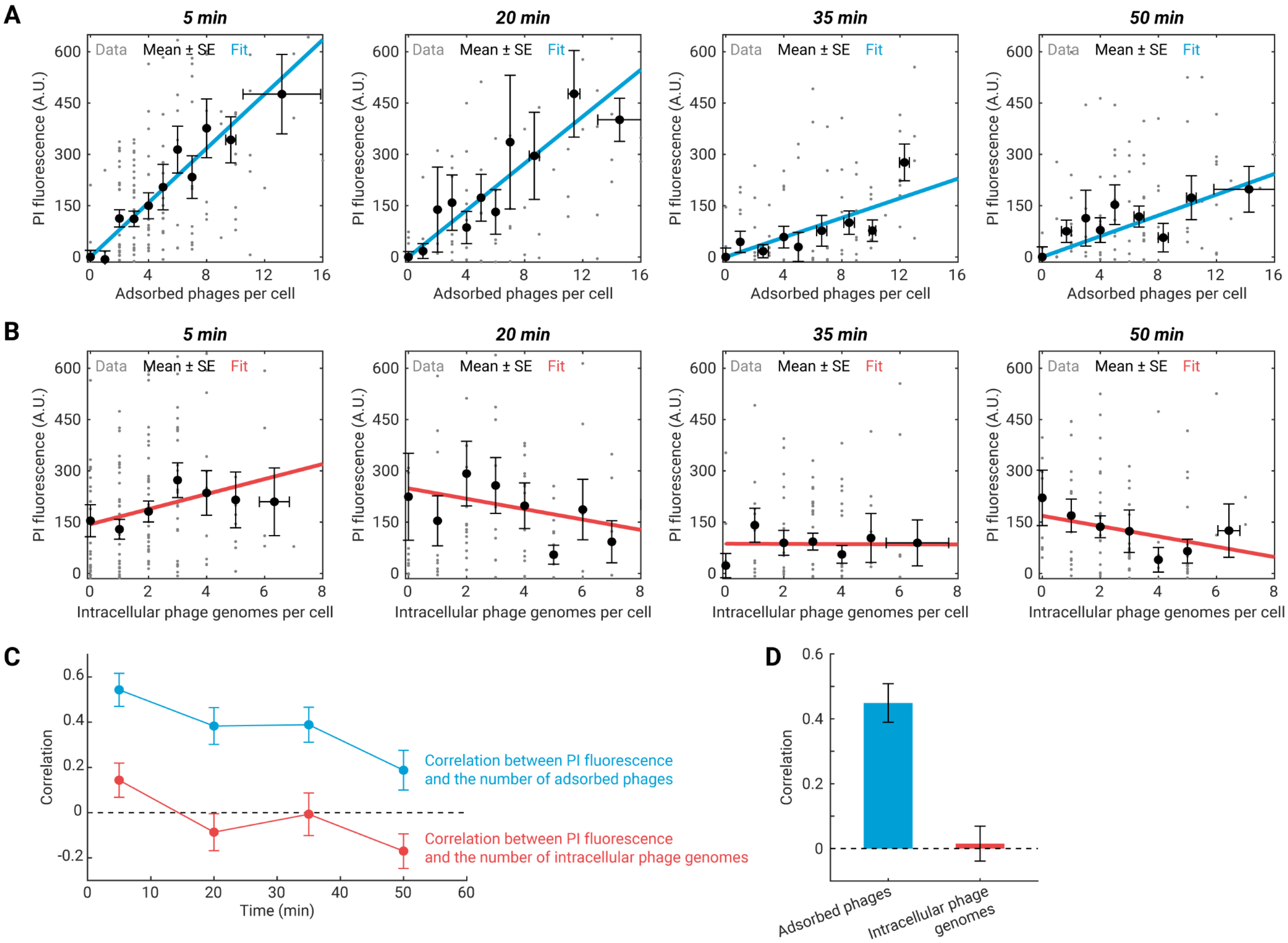


**Figure S16. Correlation between intracellular PI fluorescence and MOI.**

**(A)** The intracellular PI fluorescence as a function of the number of adsorbed phages. Data at different time points following infection (described in **Methods, Section 6**) are shown (*N* = 188, 132, 131, and 135 cells at 5, 20, 35, and 50 minutes, respectively). Gray markers, single-cell values. Black markers, mean ± SE; cells at higher MOI were binned together to allow for at least 10 cells per bin. Cyan line, linear fit with the *y*-intercept forced to zero. Data at 5 minutes is reproduced in **Figure 3B**.

**(B)** The intracellular PI fluorescence as a function of the number of intracellular phage genomes. The sample size at each time point is the same as in **Panel A**. Gray markers, single-cell values. Black markers, mean ± SE; cells at higher MOI were binned together to allow for at least 5 cells per bin. Red line, linear fit.

**(C)** Pearson correlation coefficients between intracellular PI fluorescence and the number of adsorbed phages (cyan) or the number of intracellular phage genomes (red) at each time point. Error bars indicate SE from bootstrapping.

**(D)** Intracellular PI fluorescence is not correlated with the number of intracellular phage genomes. The Pearson correlation coefficients between PI fluorescence and the number of adsorbed phages (cyan bar, $p\text{-value} \approx 4 \times 10^{-17}$) or the number of intracellular phage genomes (red bar, $p\text{-value} \approx 0.80$) are plotted. Data at 5 and 20 minutes post-infection (prior to the recovery of membrane integrity, **Figure 3C**) were pooled together for this calculation. Error bars indicate SE from bootstrapping (*N* = 320 cells).

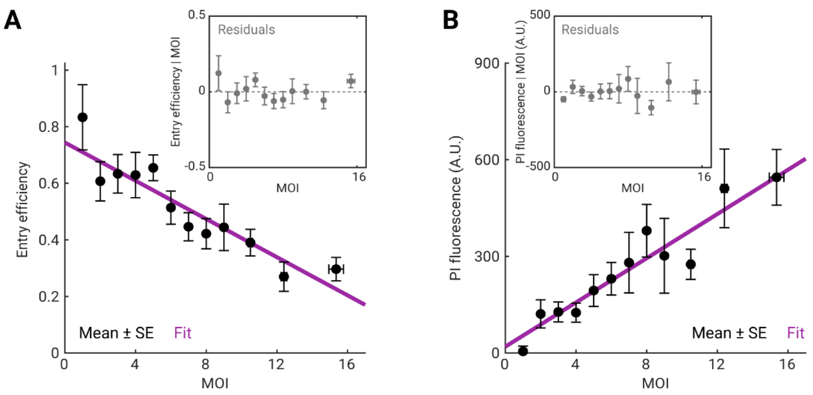


**Figure S17. Inferring a causal link between membrane permeabilization and impaired phage entry.**

**(A)** Linear regression of entry efficiency on the number of adsorbed phages (MOI) in individual cells, performed as described in **Methods, Section 6.4**. Black markers, mean ± SE (*N* = 298 cells, pooled from data at 5 and 20 minutes following infection). Magenta line, linear fit. Inset, the residuals $\text{Entry efficiency}|\text{MOI}$ represent variations in the entry efficiency for a given MOI.

**(B)** Linear regression of PI fluorescence on MOI in individual cells, performed as described in **Methods, Section 6.4**. Black markers, mean ± SE (from the same dataset as Panel A). Magenta line, linear fit. Inset, the residuals $\text{PI fluorescence}|\text{MOI}$ represent variations in the intracellular PI fluorescence for a given MOI.

The correlation between the residuals in Panel A and Panel B is shown in **Figure 3E**.

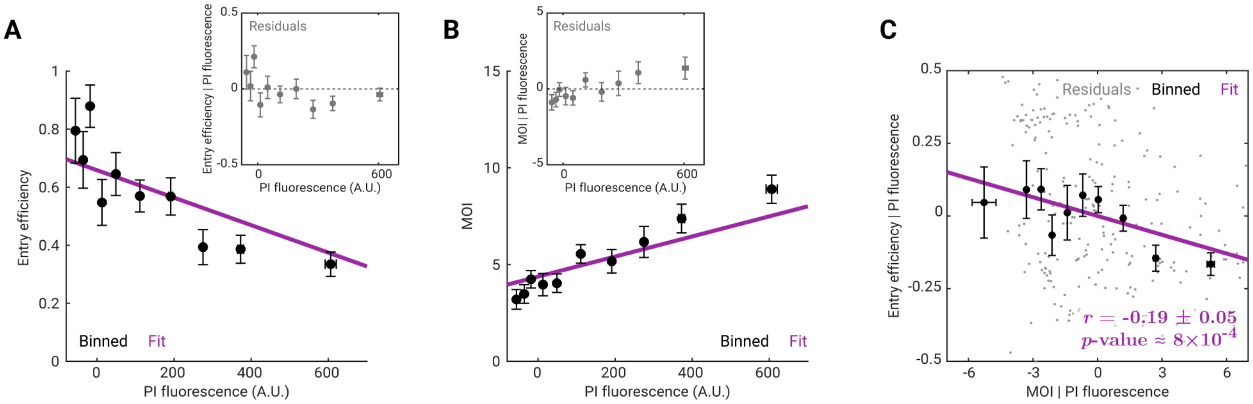


**Figure S18. Inferring additional causal links between phage adsorption and impaired phage entry.**

**(A)** Linear regression of entry efficiency on the intracellular PI fluorescence, performed as described in **Methods, Section 6.4**. Black markers, mean ± SE (30 cells per bin, $N$ = 298 cells, pooled from data at 5 and 20 minutes following infection). Magenta line, linear fit. Inset, the residuals $\text{Entry efficiency}|\text{PI fluorescence}$ represent variations in the entry efficiency for a given degree of membrane permeabilization (indicated by PI fluorescence).

**(B)** Linear regression of the number of adsorbed phages per cell (MOI) on the intracellular PI fluorescence, performed as described in **Methods, Section 6.4**. Black markers, mean ± SE (30 cells per bin, from the same dataset as Panel A). Magenta line, linear fit. Inset, the residuals $\text{MOI}|\text{PI fluorescence}$ represent variations in MOI for a given degree of membrane permeabilization (indicated by PI fluorescence).

**(C)** Phage entry efficiency and MOI, conditioned on PI fluorescence, are negatively correlated. Gray markers, residuals obtained by linear regression of the entry efficiency and of MOI on the PI fluorescence (shown in Panels A and B). Black markers, mean ± SE of the residuals (30 cells per bin). Magenta line, linear fit.

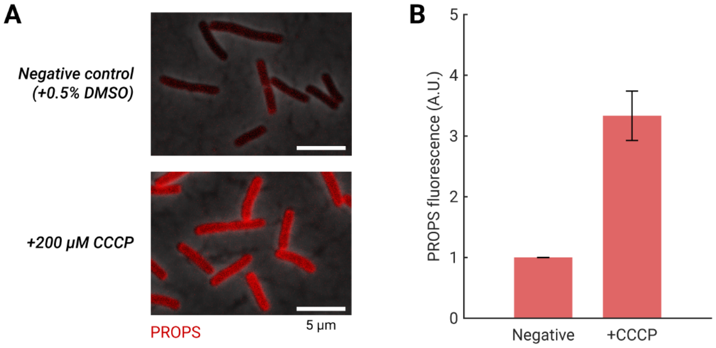


**Figure S19. Validating the CCCP treatment protocol using PROPS.**

**(A)** Fluorescence of PROPS (red signal) in cells treated with 0.5% DMSO (serving as a negative control) and in cells depolarized using 200 μM CCCP, measured as described in **Methods, Section 10**.

**(B)** Following CCCP-induced depolarization, PROPS fluorescence increased ~3-fold. This fold-change was consistent with the reported sensitivity of PROPS in LB, 2.5-fold per 100 mV (*10*), and the resting membrane potential of *E. coli*, approx. -140 mV (*42*). Error bars indicate SE from *N* = 2 independent experiments, in which 102–210 cells were analyzed for each sample in each experiment.

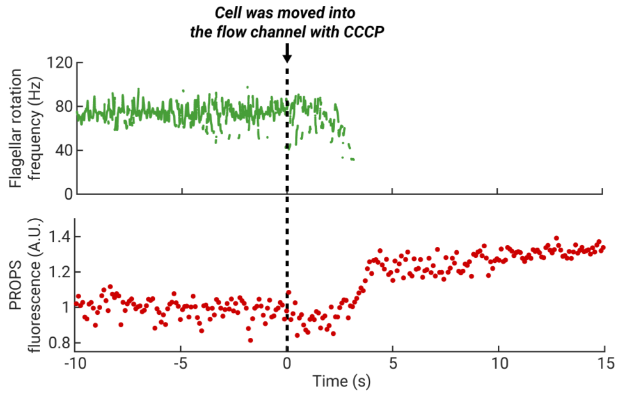


**Figure S20. Response of optically-trapped cells to CCCP treatment.**

Individual *E. coli* cells were tracked using optical traps and fluorescence microscopy, as described in **Methods, Section 13**. Under CCCP treatment, the flagellar rotation frequency (upper panel) decreased to zero, and the PROPS fluorescence (lower panel) increased by ~1.3-fold. Data from one representative cell is shown here; in total, $N$ = 9 cells, in 2 independent experiments, all exhibited similar behavior (see also **Figure S22**). Cells were grown in TB, trapped in a flow chamber, and moved into the channel with 200 μM CCCP at $t$ = 0.

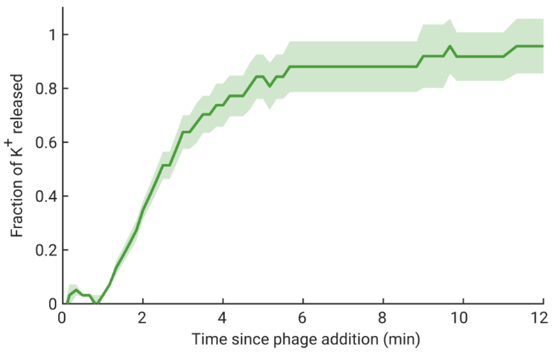


**Figure S21. Potassium efflux following phage infection.**

Efflux of potassium ions following lambda infection was measured as described in **Methods, Section 15**. Green line and shading, mean ± SE from $N$ = 2 independent experiments. Phages were added to the cell suspension at $t$ = 0. The fraction of potassium efflux was calculated using the concentrations of potassium ions in the solution (measured using an electrode) before and after phage addition.

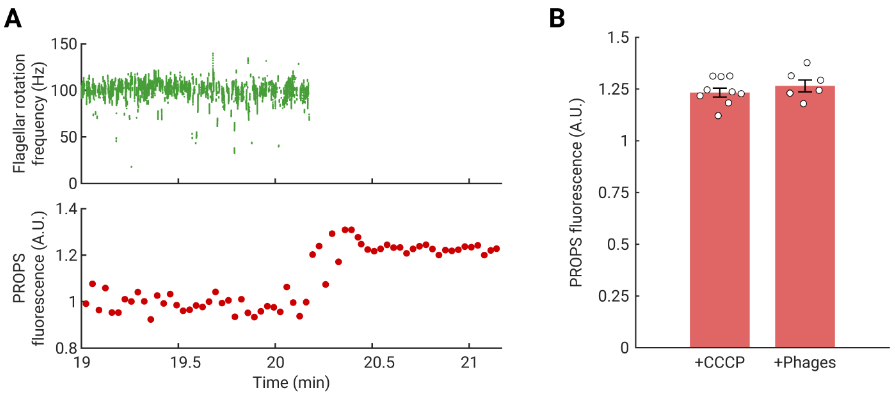


**Figure S22. Membrane depolarization of optically-trapped cells following phage adsorption, measured using PROPS.**

**(A)** Optically-trapped cells in a flow channel with lambda phages exhibited a loss of the flagellar rotation (upper panel) and an increase in PROPS fluorescence (lower panel), measured as described in **Methods, Section 13**. One representative cell is shown here; in total, $N$ = 6 cells all exhibited similar behavior. Cells were grown in TB, trapped in a flow chamber, and moved into the channel with phages at $t$ = 0.

**(B)** The increase in PROPS fluorescence following phage adsorption was similar to that following CCCP treatment. Red bars, PROPS fluorescence in cells in the flow channel containing 200 μM CCCP ($N$ = 9 cells, data for one of which is shown in **Figure S20**) and in cells in the flow channel containing phages ($N$ = 6 cells, data for one of which is shown in Panel A); error bars indicate SE. White circles, single-cell values. Cells were observed for at least 30 minutes or until motility was lost. The fluorescence of PROPS in individual cells was normalized to that before treatment or infection as described in **Methods, Section 13**.

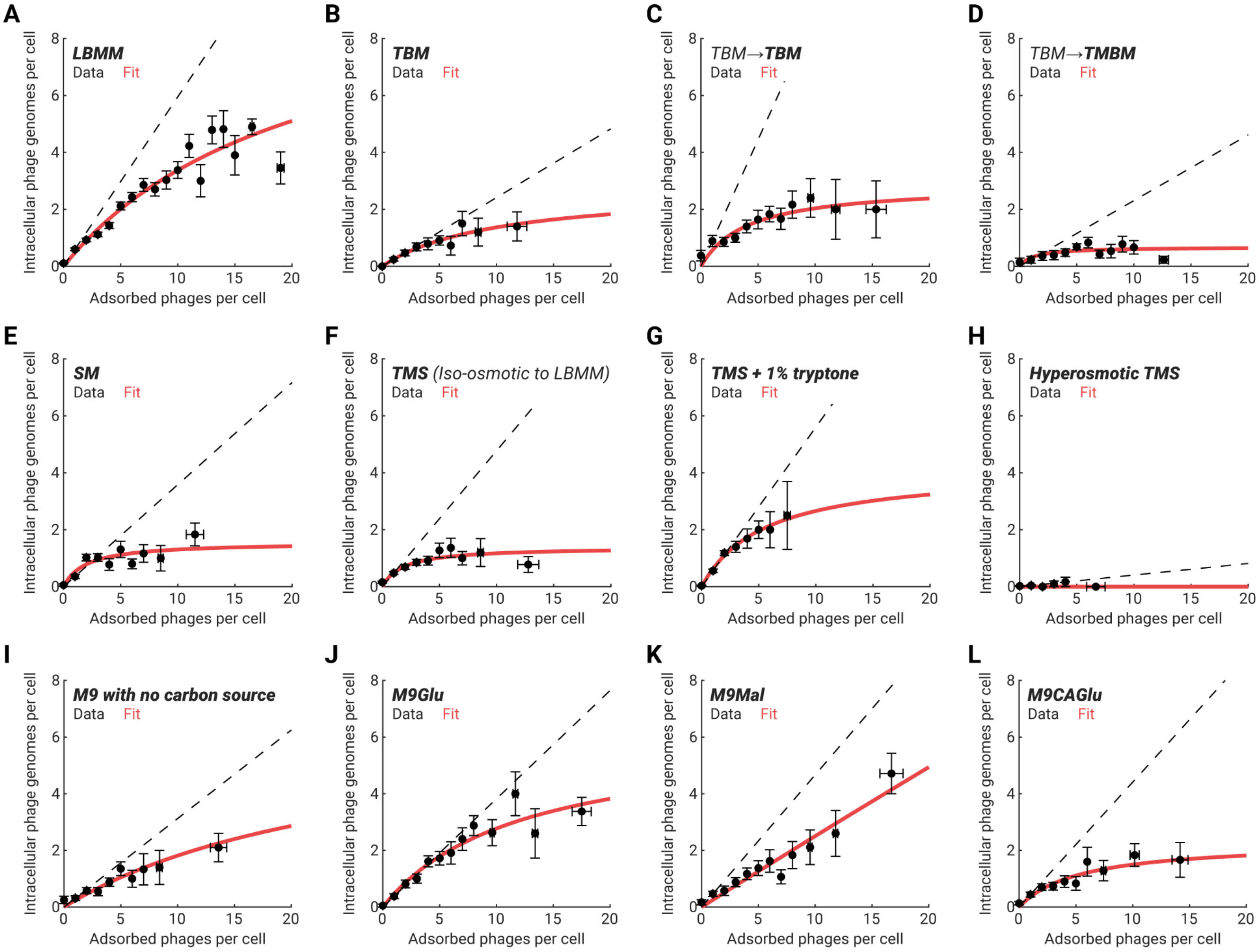


**Figure S23. Relation between the numbers of intracellular phage genomes and adsorbed phages in various infection media.**

See **Table S3** for the full names and compositions of all infection media. For all panels, cells were grown in LBMM before infection, except for Panels C and D in which cells were grown in TBM. Markers, mean ± SE, obtained using the bulk infection assay (**Methods, Section 5**); cells at higher MOI were binned together to allow for at least 10 cells per bin in Panel A, and 5 cells per bin in all other panels. Dashed line, linear scaling extrapolated from cells with one adsorbed phage. Red curve, fits to a Michaelis-Menten function (**Equation 5.1**), serving as a guide to the eye. The sample size and the fitted parameter values for each infection medium are as follows. **(A)** Infection in LBMM (reproduced in **Figures 1A** and **4A**), *N* = 1437 cells, $a$ = 10.53, $K$ = 21.24. **(B)** Infection in TBM, *N* = 223 cells, $a$ = 2.76, $K$ = 10.20. **(C)** Growth and infection in TBM, *N* = 146 cells, $a$ = 2.87, $K$ = 4.10. **(D)** Growth in TBM, infection in TMBM, *N* = 194 cells, $a$ = 0.68, $K$ = 1.25. **(E)** Infection in SM (reproduced in **Figure 4A**), *N* = 242 cells, $a$ = 1.56, $K$ = 1.97. **(F)** Infection in TMS, *N* = 514 cells, $a$ = 1.39, $K$ = 1.86. **(G)** Infection in TMS supplemented with 1% tryptone, *N* = 288 cells, $a$ = 4.13, $K$ = 5.56. **(H)** Infection in hyperosmotic TMS, *N* = 178 cells, $a$ = 0.08, $K$ = 1.39. **(I)** Infection in M9 broth with no carbon source, *N* = 188 cells, $a$ = 6.76, $K$ = 27.25. **(J)** Infection in M9Glu, *N* = 223 cells, $a$ = 6.16, $K$ = 12.23. **(K)** Infection in M9Mal, *N* = 218 cells, $a$ = 190, $K$ = 747. **(L)** Infection in M9CAGlu, *N* = 213 cells, $a$ = 2.31, $K$ = 5.39.

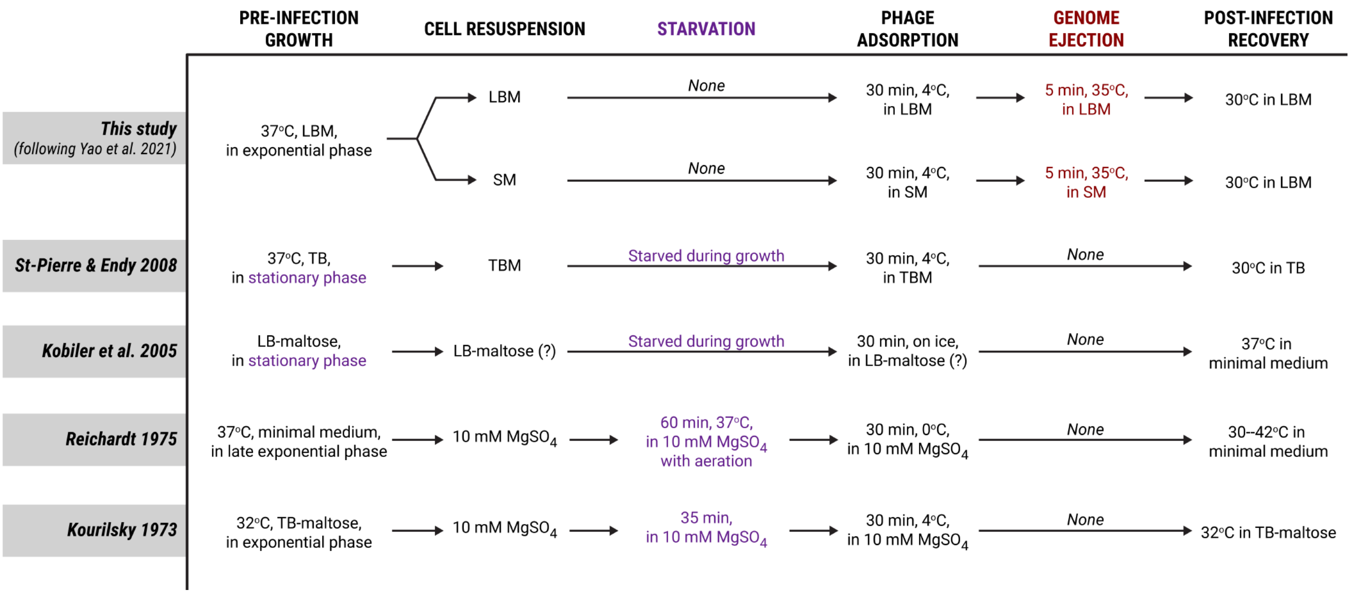


**Figure S24. Comparing the lysogenization assay in this study vs. the literature.**

Our bulk lysogenization assay (with the entry step performed in either LBM or SM, as described in **Methods, Section 16**) is compared against the procedures in Kourilsky 1973 (*27*), Reichardt 1975 (*28*), Kobiler et al. 2005 (*29*), and St-Pierre & Endy 2008 (*23*). In our assay, cells in the exponential growth phase were not starved prior to infection (text in purple). In addition, our assay included an explicit step to trigger phage ejection in the same medium as the adsorption step (text in red). For more details, see **Methods, Section 16.1**.

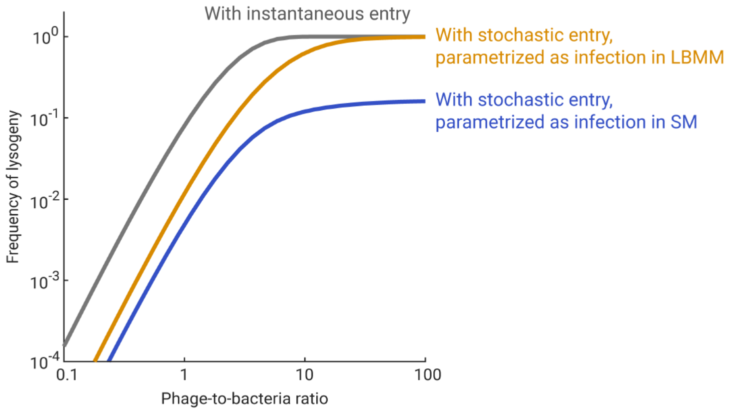


**Figure S25. Frequency of lysogeny as a function of phage-to-bacteria ratio, predicted using different models of entry dynamics.**

The frequency of lysogeny was predicted as a function of the phage-to-bacteria ratio, using **Equations 17.6** and **17.7**, with $a$ = 1, $\mathrm{MOI}^*$ = 3, and $q_{\max}$ = 1. Gray curve, model prediction when phage entry is instantaneous. Orange and blue curves, model predictions when stochastic phage entry, parametrized as infection in LBMM and SM, respectively, is incorporated into the model. Parametrization of the entry dynamics in LBMM and SM was done using the bulk infection assay (**Methods, Section 5**) as described in **Figure S23**.

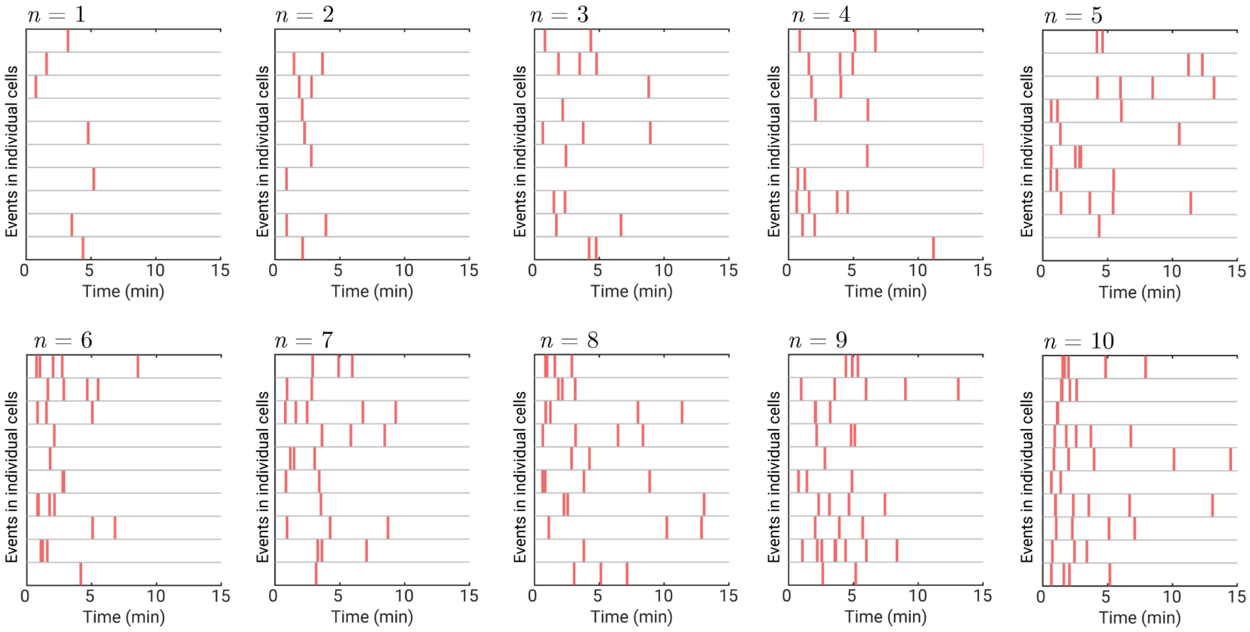


**Figure S26. Simulated time series of phage entry events.**

Time series of phage entry events, obtained using the stochastic simulation described in **Methods, Section 19**. Here, 10 cells are shown for each value of the number of adsorbed phages per cell, $n$, from 1 to 10. The parameters used for simulation were described in **Methods, Section 8.3.3**. In the simulation, phage adsorptions occurred synchronously at $t$ = 0. Vertical lines in red, entry events.

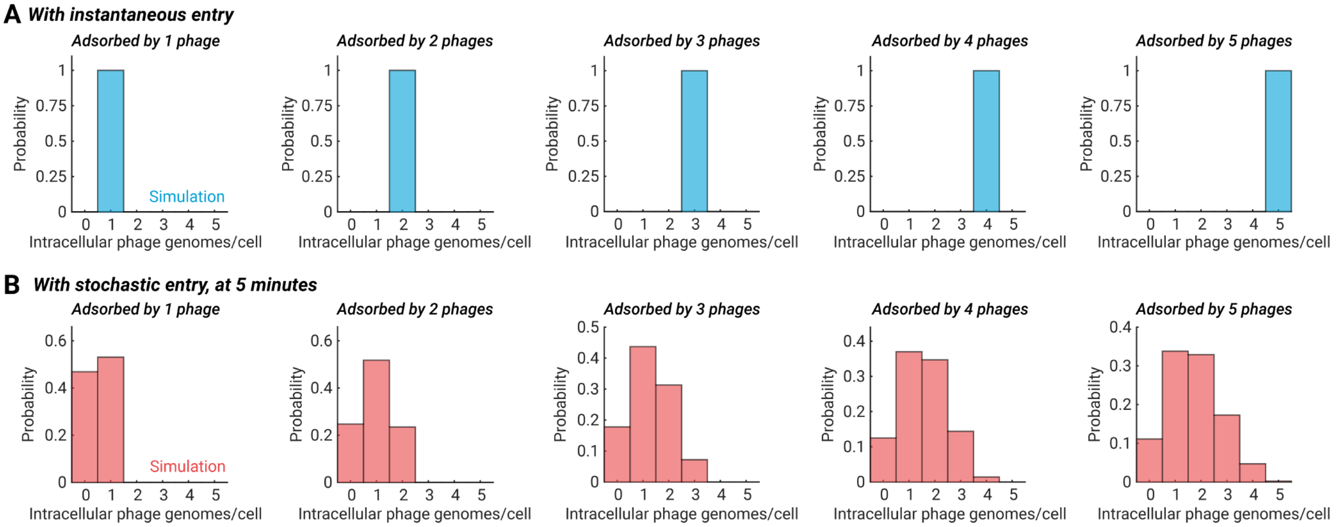


**Figure S27. Distributions of intracellular phage numbers, simulated using different models of entry kinetics.**

In all graphs, the bars indicate the distribution of the intracellular phage numbers in cells adsorbed by 1–5 phages, obtained using the stochastic simulation described in **Methods, Section 19** (*N* = 1000 cells for each value of adsorbed phage numbers). **(A)** Simulation assuming all adsorbed phages enter the cell instantaneously. These distributions were used as the MOI to simulate the infection outcome following instantaneous entry in **Figure 4E**. **(B)** Simulation assuming stochastic entry; the number of intracellular phage genomes was counted at 5 minutes. These distributions were used as the MOI to simulate the infection outcome following stochastic entry in **Figure 4E**.

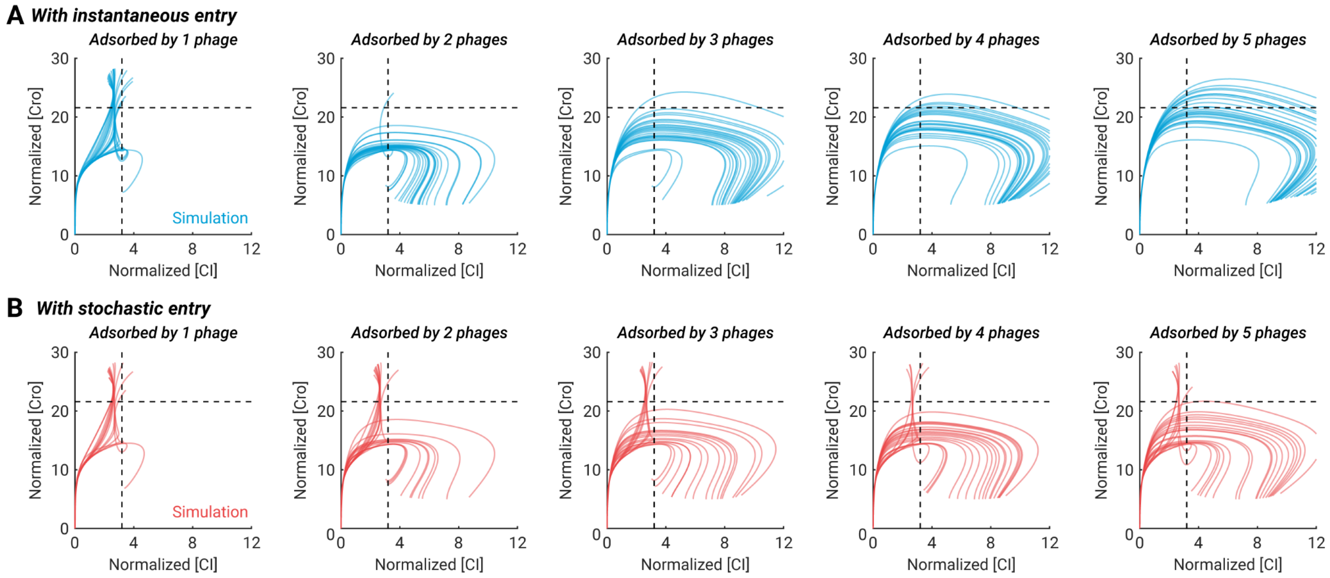


**Figure S28. Trajectories of Cro and CI concentrations, simulated using different models of entry kinetics.**

In all graphs, the curves indicate the predicted single-cell trajectories of Cro and CI concentrations in cells adsorbed by 1–5 phages, obtained using the stochastic simulation described in **Methods, Section 20** ($N$ = 1000 cells for each value of adsorbed phage numbers). Trajectories of 50 cells are shown in each graph; some trajectories for cells with 2 adsorbed phages are reproduced in **Figure 4D**. Dashed lines, the decision thresholds in Cro and CI concentrations, used to determine one of four possible outcomes (failed infection, lysis, lysogeny, and mixed outcome) as described in (*1*). **(A)** Simulation assuming all adsorbed phages enter the cell instantaneously, and the single-cell MOI is the number of adsorbed phages (the distributions of which are shown in **Figure S27A**). **(B)** Simulation assuming stochastic phage entry, and the single-cell MOI is the number of intracellular phage genomes at 5 minutes (the distributions of which are shown in **Figure S27B**).

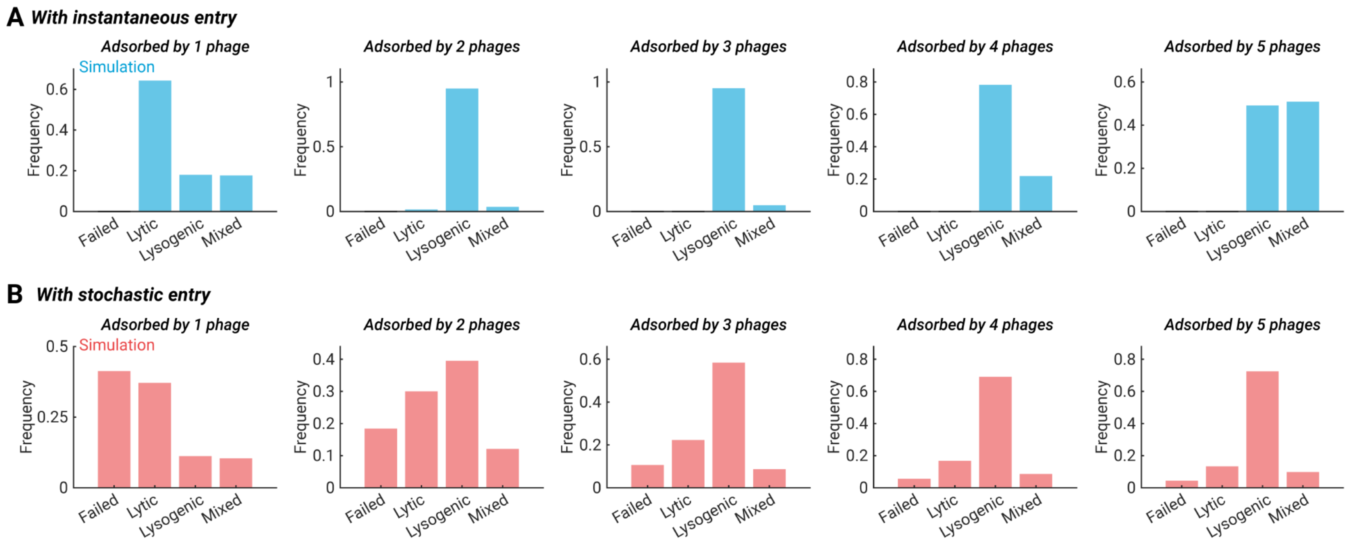


**Figure S29. Distributions of infection outcomes, simulated using different models of entry kinetics.**

In all graphs, the bars indicate the predicted distributions of infection outcomes in cells adsorbed by 1–5 phages, obtained using the stochastic simulation described in **Methods, Section 20** (*N* = 1000 cells for each value of adsorbed phage numbers). Infection outcome was determined using the trajectories of Cro and CI concentrations (shown in **Figure S28**) as described in (*1*). Only the lytic and lysogenic fates were used to calculate the decision curve (shown in **Figure 4E**). **(A)** Simulation assuming all adsorbed phages enter the cell instantaneously. **(B)** Simulation assuming stochastic phage entry.

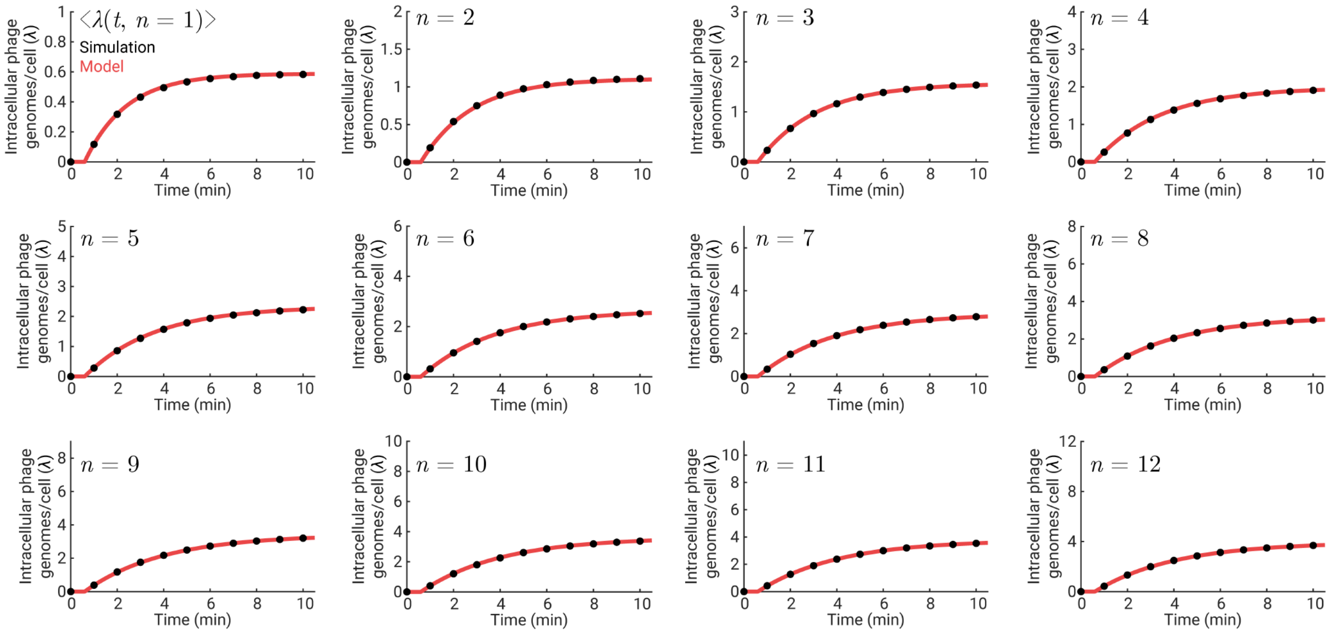


**Figure S30. Comparison of the stochastic simulation and analytical predictions for the time-dependent average number of intracellular phage genomes.**

In all graphs, the markers indicate the time-dependent average number of intracellular phage genomes, $\langle\lambda(t)\rangle$, in cells adsorbed by $n$ = 1–12 phages. These values were obtained using the stochastic simulation described in **Methods, Section 19** (*N* = 10000 simulated cells for each $n$ value). Red curves, analytical predictions by the stochastic model of phage entry kinetics, using **Equation 18.3**. The parametrization of $(\eta_n, k_n, \tau_n)$ used for both the stochastic simulation and the analytical prediction was described in **Methods, Section 8.3.3**.

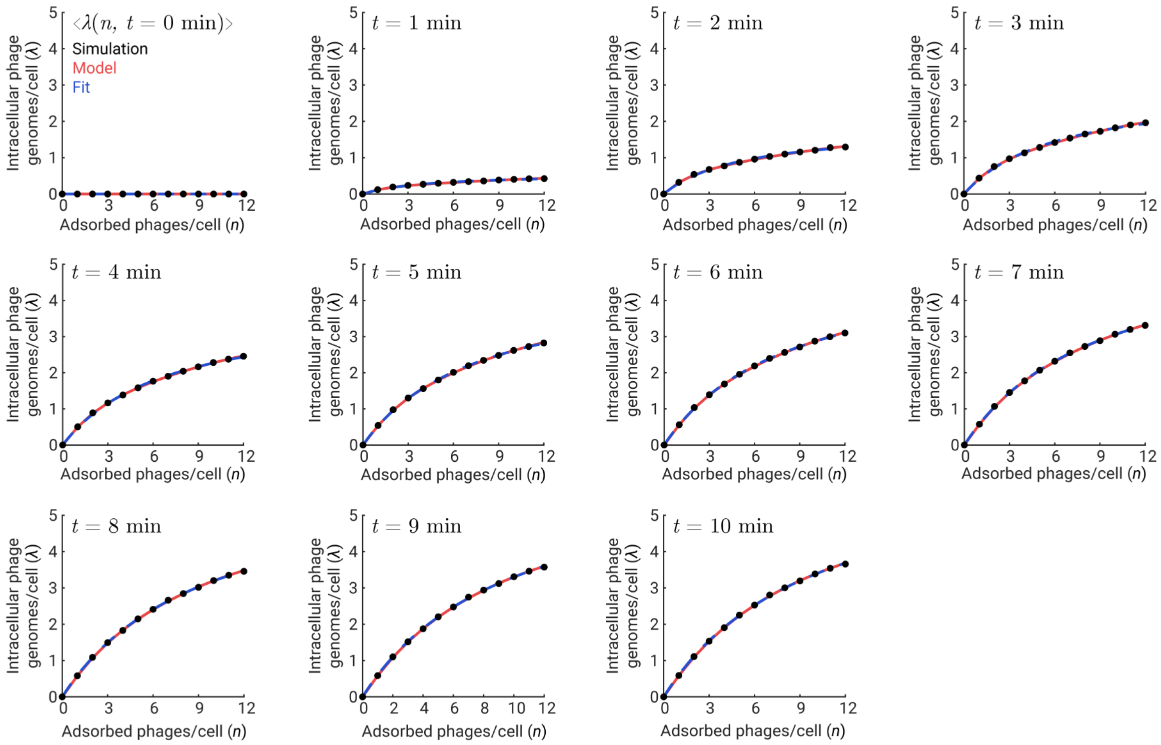


**Figure S31. Comparison of the stochastic simulation, analytical predictions, and phenomenological fit for the relation between the numbers of intracellular phage genomes and adsorbed phages.**

In all graphs, the markers indicate the average number of intracellular phage genomes, $\langle \lambda \rangle$, as a function of the number of adsorbed phages, $n$, at $t$ = 0–10 min. These values were obtained using the stochastic simulation described in **Methods, Section 19** (*N* = 10000 simulated cells for each $n$ value). Red curves, analytical predictions by the stochastic model of phage entry kinetics, using **Equation 18.3**. The parametrization of $(\eta_n, k_n, \tau_n)$ used for both the stochastic simulation and the analytical prediction was described in **Methods, Section 8.3.3**. Blue curves, fit of **Equation 5.1** to the simulated data.

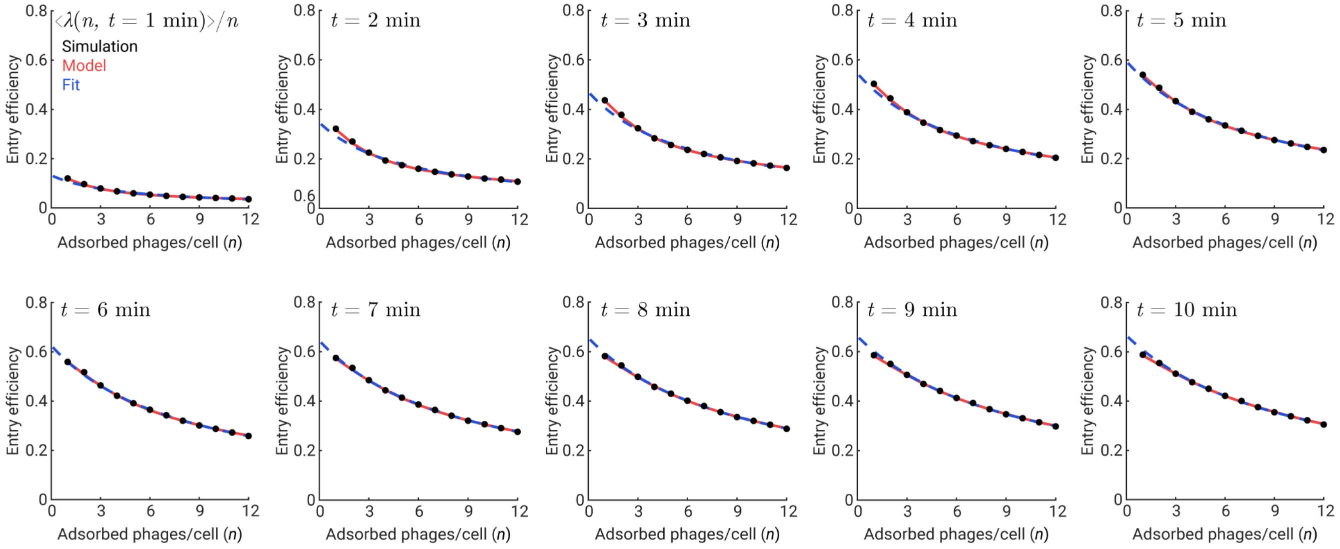


**Figure S32. Comparison of the stochastic simulation, analytical predictions, and phenomenological fit for the efficiency of phage entry.**

In all graphs, the markers indicate the efficiency of phage entry, $\langle\lambda\rangle/n$, as a function of the number of adsorbed phages, $n$, at $t$ = 1–10 min. These values were obtained using the stochastic simulation described in **Methods, Section 19** (*N* = 10000 simulated cells for each $n$ value). Red curves, analytical predictions by the stochastic model of phage entry kinetics, using **Equation 18.3**, divided by the corresponding value of $n$. The parametrization of $(\eta_n, k_n, \tau_n)$ used for both the stochastic simulation and the analytical prediction was described in **Methods, Section 8.3.3**. Blue curves, fit of **Equation 5.2** to the simulated data.

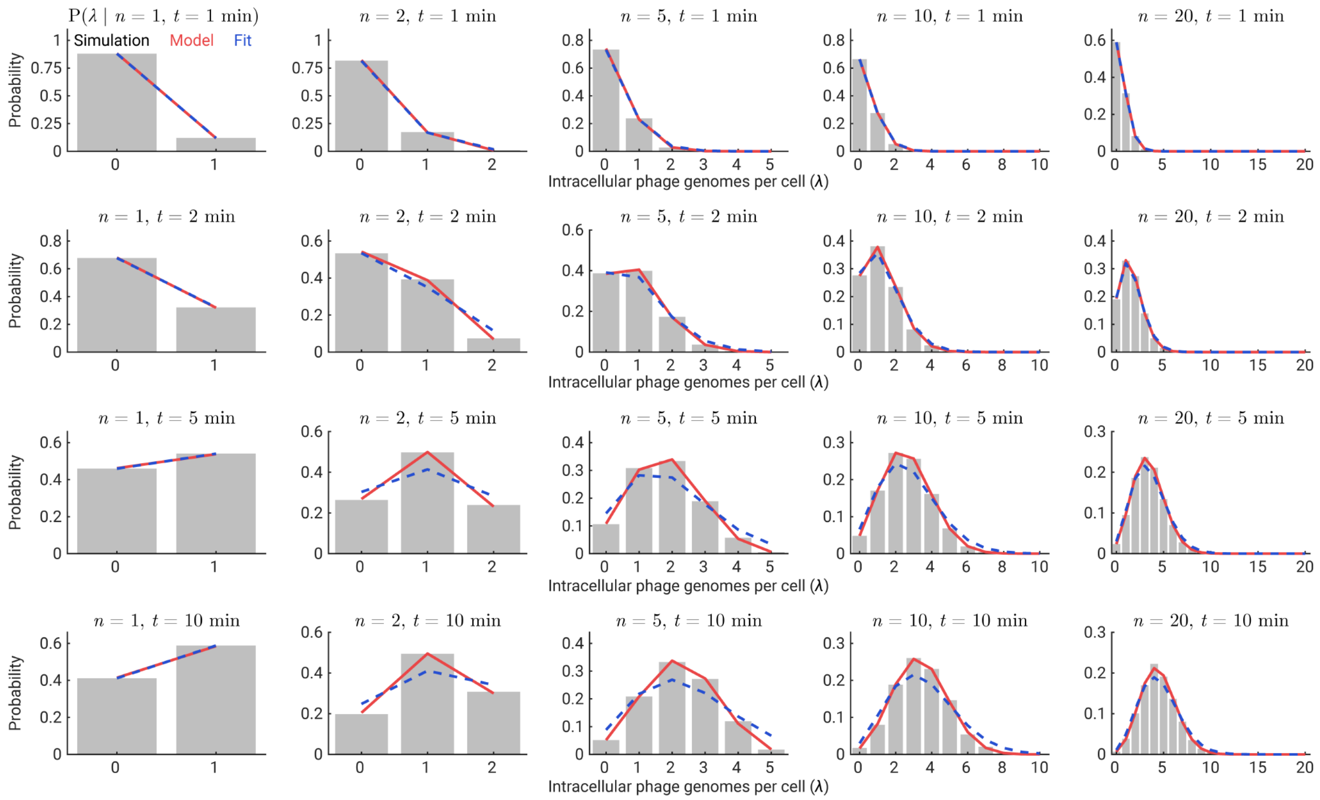


**Figure S33. Comparison of the stochastic simulation, analytical predictions, and phenomenological fit for the distribution of the intracellular phage number.**

In all graphs, the gray bars indicate the histogram of intracellular phage numbers, $\lambda$, at time $t$ in cells adsorbed by $n$ phages. Histograms for $n$ = 1, 2, 5, 10, and 20 at $t$ = 1, 2, 5, and 10 min are shown. These data were obtained using the stochastic simulation described in **Methods, Section 19** (*N* = 10000 simulated cells for each *n* value). Red curves, analytical predictions by the stochastic model of phage entry kinetics, using **Equation 18.29**. The parametrization of $(\eta_n, k_n, \tau_n)$ used for both the stochastic simulation and the analytical prediction was described in **Methods, Section 8.3.3**. Blue curves, fit of **Equation 5.3** to the simulated data.

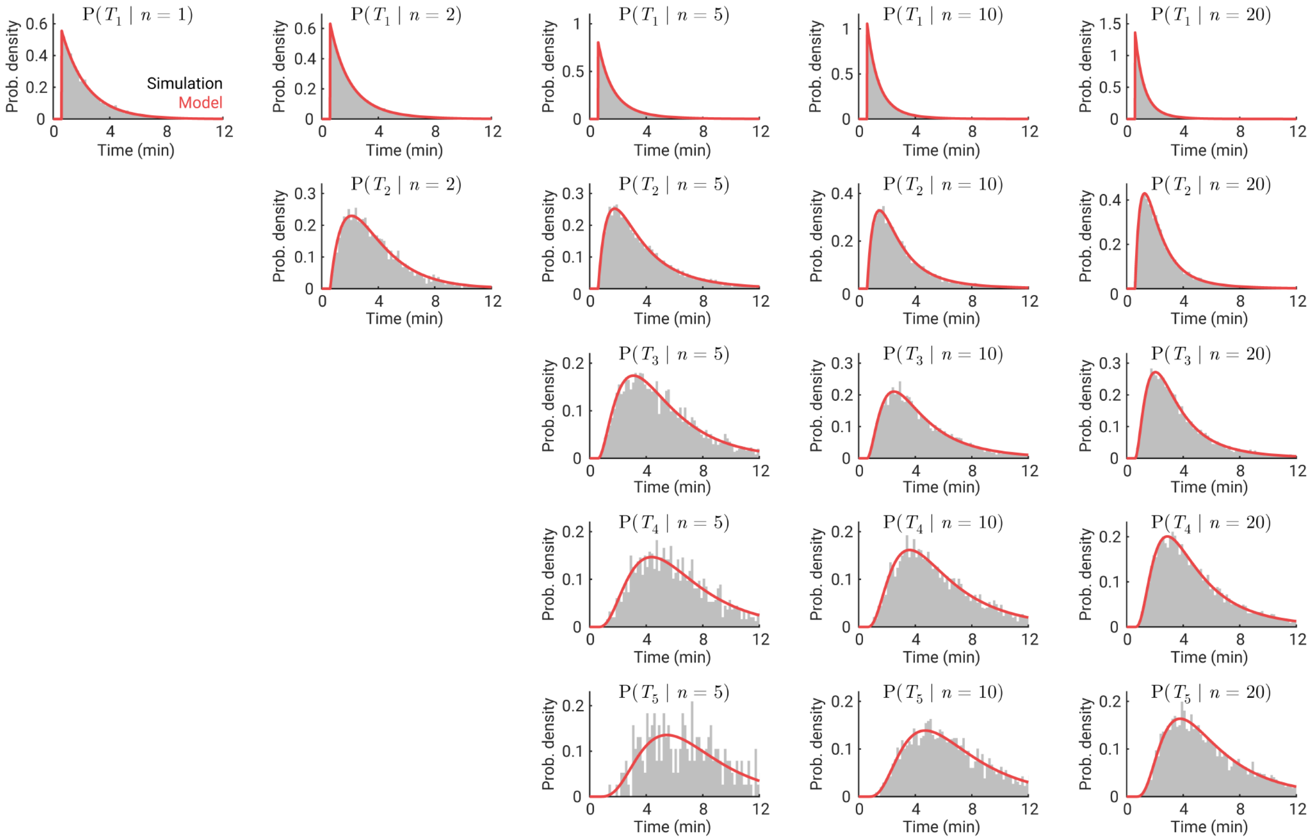


**Figure S34. Comparison of the stochastic simulation and analytical predictions for the distribution of phage entry time.**

In all graphs, the gray bars indicate the histogram of the entry time of the $i$-th phage, $T_i$, in cells adsorbed by $n$ phages. Histograms for $n$ = 1, 2, 5, 10, and 20 and $i$ = 1–5 are shown ($T_i(n)$ is undefined for $i > n$). These data were obtained using the stochastic simulation described in **Methods, Section 19** (*N* = 10000 simulated cells for each $n$ value). Red curves, analytical predictions by the stochastic model of phage entry kinetics, using **Equation 18.25**. The parametrization of $(\eta_n, k_n, \tau_n)$ used for both the stochastic simulation and the analytical prediction was described in **Methods, Section 8.3.3**.

### SUPPLEMENTARY MOVIE

**Caption for Movie S1: Phage adsorption results in membrane depolarization.**

The motility of phage-adsorbed cells was tracked using dual-trap optical tweezers as described in **Methods, Section 13**. Fluorescence imaging was performed continuously, and the cell's flagellar rotation frequency was inferred from the trap position (graph on the bottom). Dotted line, approximate boundary of the cell body. At the beginning of the movie, the cell was trapped in a flow channel with phage perfusion. Phages were visualized using SYTOX Orange (green signal), which stains encapsidated phage DNA. The motility of the cell was indicated by the oscillations in the trap positions. At $t$ = 0, a phage adsorbed to the cell; the rotation of the fluorescent spot corresponded to the rotation of the cell body. From $t$ = 6 to 16 s, the cell was moved to a flow channel containing blank medium to prevent additional phage adsorptions. The cell remained motile with a flagellar rotation frequency of approx. 100 Hz for the next ~30 s. At $t$ = 45 s, the cell motility was lost, and the inferred flagellar rotation frequency decreased to zero, indicating membrane depolarization. The phage particle remained stained for encapsidated DNA, indicating that phage-induced depolarization precedes phage entry. This movie was recorded and played at 10 frames per second. A snapshot from this movie is shown in **Figure 3F**. Scale bar, 1 μm.

### SUPPLEMENTARY TABLES

#### Table S1: Bacterial strains, phages, and plasmids used in this study

| Strain | Description | Use in work | Source |
|---|---|---|---|
| **BACTERIA** | | | |
| MG1655 | Wild-type *E. coli* | All infection experiments | Lab stock |
| LE392 | *glnV* (*supE*44), *tryT* (*supF*58); amber suppressor (*43*). | Production of $\lambda_{TY11}$ phages; phage titering | Lab stock |
| **PHAGE** | | | |
| $\lambda_{TY11}$ | λ *cI*857 *Pam*80 *stf*::P1*parS-kan*$^R$<br>Temperature-sensitive mutation in the *cI* gene.<br>Amber mutation in the *P* gene; replication-deficient when infecting MG1655 cells.<br>Containing a *parS* sequence and a kanamycin-resistance cassette at 21029–22557 bp (*stf* locus). | All experiments | (*1*) |
| **PLASMIDS** | | | |
| p2973 | mCherry-P1Δ30ParB and YGFP-pMT1Δ23ParB under the control of $P_{trc}$ (*amp*$^R$).<br>mCherry-P1Δ30ParB is denoted as mCherry-ParB in this study. YGFP-pMT1Δ23ParB, though expressed, is not imaged in this study. | Labeling of intracellular phage genomes | (*3*) |
| pALA3047 | CFP-P1Δ30ParB under the control of $P_{trc}$ (*amp*$^R$).<br>CFP-P1Δ30ParB is denoted as CFP-ParB in this study. | Labeling of intracellular phage genomes | Gift of Stuart Austin |
| pACYC177-$P_{late}$*D-mTurquoise2 | gpD-mTurquoise2 under the control of $P_{R'}$ (*amp*$^R$). | Labeling of phage capsids | This work; constructed as described in (*2*) |
| pACYC177-$P_{late}$*D-EYFP | gpD-EYFP under the control of $P_{R'}$ (*amp*$^R$). | Labeling of phage capsids | (*2*) |
| pJMK001 | Proteorhodopsin optical proton sensor (PROPS) under the control of $P_{araBAD}$ (*amp*$^R$). | Measurements of the cell's membrane potential | (*10*); acquired from Addgene (#33780) |

#### Table S2: Chemical reagents used in this study

| Solution | Manufacturer | Stock concentration | Storage condition & Notes |
|---|---|---|---|
| Ampicillin | Fisher Scientific | 100 mg/mL | -20$^{o}$C |
| Kanamycin | Fisher Scientific | 50 mg/mL | -20$^{o}$C |
| NaCl | Fisher Scientific | 5 M | Room temperature (RT) |
| NaOH | Fisher Scientific | 1 M | RT |
| $MgSO_4$ | Fisher Scientific | 1 M | RT |
| Tris-Cl, pH 7.5 | Fisher Scientific | 1 M | RT |
| Maltose | Fisher Scientific | 20% | RT |
| Glucose | Fisher Scientific | 20% | RT |
| *L*-arabinose | Sigma-Aldrich | 20% | RT |
| Glycerol | Fisher Scientific | N/A | Diluted to 15% for cell storage at -80$^{o}$C |
| Isopropyl-β-thiogalactoside (IPTG) | Sigma-Aldrich | 500 mM | -20$^{o}$C |
| Carbonyl cyanide *m*-chlorophenyl hydrazone (CCCP) | Acros Organics | 40 mM in DMSO | -20$^{o}$C |
| SYTOX Orange | Invitrogen | 5 mM in DMSO | -20$^{o}$C in the dark |
| 4′,6-diamidino-2-phenylindole (DAPI) | Invitrogen | 500 mg/mL | -20$^{o}$C in the dark |
| All-*trans* retinal | Sigma-Aldrich | 20 mM in ethanol | -20$^{o}$C in the dark; for each experiment, a 5 mM substock is prepared in water, stored on ice in the dark, and used within the day |
| Propidium iodide (PI) | Invitrogen | 1.5 mM | 4$^{o}$C |
| Formaldehyde | Fisher Scientific | 37% | RT |
| Ethanol | Decon Labs | N/A | RT |
| Dimethyl sulfoxide (DMSO) | Fisher Scientific | N/A | RT |

#### Table S3: Growth media and buffers used in this study

| Medium or buffer | Recipe<br>*(Including manufacturer information if not in Table S2)* | Storage condition & Notes |
|---|---|---|
| LB (Lennox) | By w/v, 1% tryptone (BD Biosciences), 0.5% yeast extract (BD Biosciences), and 0.5% NaCl; pH adjusted using 1 mM NaOH. | RT after autoclaving |
| LBM | LB and 10 mM $MgSO_4$. | Made fresh for each experiment |
| LBMM | LB, 0.2% maltose, and 10 mM $MgSO_4$. | Made fresh for each experiment |
| LBGM | LB, 0.2% glucose, and 10 mM $MgSO_4$. | Made fresh for each experiment |
| Tryptone broth (TB) | By w/v, 1% tryptone and 0.8% NaCl. | RT after autoclaving |
| TBM | TB and 10 mM $MgSO_4$. | Made fresh for each experiment |
| NZYM | By w/v, 2.2% NZYM media (Teknova); pH adjusted using 10 mM NaOH. | 4$^o$C after autoclaving |
| M9 with no carbon source | M9 minimal salts broth media without carbon source (Teknova). | RT |
| M9Glu | M9 minimal salts broth and 0.4% glucose. | Made fresh for each experiment |
| M9Mal | M9 minimal salts broth and 0.4% maltose. | Made fresh for each experiment |
| M9CAGlu | M9 minimal salt broth with 1% glucose and 0.1% casamino acids (Teknova). | RT |
| SM | 100 mM NaCl, 10 mM $MgSO_4$, 50 mM Tris-Cl pH 7.5, and 0.01% (w/v) gelatin. When used in small amount, purchased from Teknova. | RT |
| PBS | 1×, prepared from a 10× stock (Invitrogen). | RT |
| PBSM | 1× PBS and 10 mM $MgSO_4$. | Made fresh for each experiment |
| Motility buffer (MB) | 10 mM $K_3PO_4$ pH 7.0 (EMD Millipore), 70 mM NaCl, and 0.1 mM EDTA (Promega Life Sciences). | RT |
| Trap motility buffer (TMB) | 70 mM NaCl, 100 mM Tris-Cl (pH 7.5), and 2% (w/v) glucose, supplemented with an oxygen-scavenging system (290 µg/mL pyranose oxidase, Sigma-Aldrich, and 65 µg/mL catalase, EMD Millipore). | -20$^o$C |
| TMBM | TMB and 10 mM $MgSO_4$. | Made fresh for each experiment |
| Iso-osmotic Tris-Magnesium-Sodium buffer (TMS) | 10 mM Tris-Cl pH 7.5, 10 mM $MgSO_4$, and 125 mM NaCl; osmolality of approx. 280 mOsm/kg, similar to LB (*44*) when supplemented with 10 mM $MgSO_4$. | RT |
| TMS + 1% tryptone | Iso-osmotic TMS and 1% tryptone (w/v). | Made fresh for each experiment |
| Hyperosmotic TMS | 10 mM Tris-Cl pH 7.5, 10 mM $MgSO_4$, and 250 mM NaCl (additional 250 mOsm/kg compared to LBM). | RT |

**Table S4: Filter sets used for fluorescence microscopy**

| **Filter sets** *(Semrock catalog #)* | **Fluorophore** |
|---|---|
| DAPI-3060A | DAPI |
| CFP-2432C | gpD-mTurquoise2, CFP-ParB |
| YFP-2427B | gpD-EYFP |
| LED-TRITC-A | SYTOX Orange |
| mCherry-C | mCherry-ParB, Propidium iodide (PI) |
| LED-Cy5-5070A | PROPS |

**Table S5: Variables and parameters in the mathematical models**

| Symbol | Description | Section or equation |
|---|---|---|
| $M$ | Phage-to-bacteria ratio in the infection mixture | Section 17 |
| $f_{\text{lysogeny}}(M)$ | Frequency of lysogeny as a function of the phage-to-bacteria ratio | Equations 17.6 and 17.7 |
| $n$ | Number of adsorbed phages for the cell of interest | Section 18.1 |
| $\lambda$ | Number of intracellular phage genomes for the cell of interest | Section 18.1 |
| $t$ | Time of interest | Section 18.1 |
| $\eta$ | Entry probability; the probability that an adsorbed phage is capable of entering the cell (at infinite time) | Section 18.1 |
| $k$ | Rate (or probability per unit of time) of entry initiation by each entry-capable phage | Section 18.1 |
| $\tau$ | Time for an entry-initiated phage to translocate into the cell and get detected | Section 18.1 |
| $(\eta_n, k_n, \tau_n)$ | Parameters specific to cells adsorbed by $n$ phages | Section 8.3.3 |
| $m$ | Number of entry-capable phages (among the adsorbed phages) | Section 18.2 |
| $a_{\text{ejection}}$ | Propensity of ejection (in the simulation) | Equation 19.1 |
| $t_{i-1,i}$ | Waiting time between the ($i$-1)-th and $i$-th entry initiations | Section 18.2 |
| $T_i$ | Time of the $i$-th phage entry; the time between phage adsorption and when the $i$-th phage genome is detected inside the cell | Section 18.2 |
| $f_{T_i\|m}(t)$ | Probability density that the $i$-th phage entry occurs at time $t$, given $m$ entry-capable phages | Equation 18.19 |
| $F_{T_i\|m}(t)$ | Cumulative probability that the $i$-th phage entry has occurred by time $t$, given $m$ entry-capable phages; | Equation 18.20 |
| $\langle T_i(m) \rangle$ | Average time of the $i$-th phage entry, in cells with $m$ entry-capable phages | Equation 18.21 |
| $P(m\|n)$ | Probability that a cell has $m$ entry-capable phages, given $n$ adsorbed phages | Equation 18.22 |
| $f_{T_i\|n}(t)$ | Probability density that the $i$-th phage entry occurs at time $t$, given $n$ adsorbed phages | Equation 18.25;<br>for $n = 1$, Equation 18.31 |
| $\langle T_i(n) \rangle$ | Average time of the $i$-th phage entry, in cells with $n$ adsorbed phages | Equation 18.26;<br>for $n = 1$, Equation 18.32 |
| $P(\lambda = i\|n, t)$ | Probability that a cell has $\lambda = i$ intracellular phage genomes at time $t$, given $n$ adsorbed phages | Equation 18.29;<br>for $n = 1$, Equation 18.33 |
| $\langle \lambda(n, t) \rangle$ | Average number of intracellular phage genomes at time $t$, in cells with $n$ adsorbed phages | Equation 18.3;<br>for $n = 1$, Equation 18.34 |
